# A novel beta-adrenergic like octopamine receptor modulates the audition of malaria mosquitoes and serves as insecticide target

**DOI:** 10.1101/2022.08.02.502538

**Authors:** M Georgiades, CA Alampounti, J Somers, M Su, D Ellis, J Bagi, W Ntabaliba, S Moore, JT Albert, M Andres

**Author notes:** These authors share first authorship (M Georgiades, CA Alampounti).

## Abstract

Hearing is an essential sense in the life cycle of malaria mosquitoes. Within large swarms formed transiently at dusk, mosquitoes acoustically recognize their mating partners by their wingbeats. Indeed, malaria mosquitoes only respond to the flight tones of mating partners during swarm time. This phenomenon implies a sophisticated context- and time-dependent modulation of mosquito audition, the mechanisms of which are still largely unknown. Using transcriptomics, we identify a complex network of candidate neuromodulators regulating mosquito hearing. Among them, octopamine stands out as regulator of the auditory performance during swarm time. To explore octopamine’s roles in mosquito hearing, we carried out an in-depth analysis of octopamine-mediated effects on auditory function. We found that octopamine affects the properties of the mosquito ear on multiple levels: it modulates the tuning and stiffness of the flagellar sound receiver and it controls the erection of antennal fibrillae in males. We found that two different receptors are driving octopamine’s auditory roles, including a novel beta octopamine receptor. We also demonstrate that the octopaminergic auditory control system can be targeted by insecticides. Our findings identify octopamine signalling as a key component of hearing and mating partner detection in malaria mosquitoes, and as a potential novel target for mosquito control.

## Introduction

Sensory organs extract information from the environment. As environmental conditions are dynamic, sensory organs have evolved modulatory mechanisms that enable sensory plasticity to adapt their physiology to these external changes. The acoustic detection of mating partners in swarms by malaria mosquitoes constitutes a superb example of the adaptation of a sensory organ -the mosquito ear-to a transient change in the sensory ecology. Malaria mosquito swarms are transitory aggregations of up to thousand mosquitoes that take place every sunset and last for a brief period of time of around 30 minutes (1,2). Within the swarm, mosquitoes are exposed to an acoustically challenging, potentially very noisy, environment. It is against the acoustic backdrop of the swarm that mosquitoes identify and locate the flight tones of the mating partners (3–5).

A remarkable feature of Johnston’s organ (JO), the mosquitoes’ ‘inner ear’, is its efferent control system (6), which – across insects - has so far only been reported for mosquitoes. The neuromodulatory efferent network seems to be an intrinsic component of mosquito audition, as the ears of three distinct disease-transmitting species (*Culex quinquefasciatus, Anopheles gambiae and Aedes aegypti)* - evolutionarily separated by more than 150 million years - are endowed with efferent input (7). The efferent fibers release several neurotransmitters including the biogenic amines octopamine and serotonin and the inhibitory neurotransmitter GABA (6). Efferent input may hold the key to understanding the extraordinary performance of mosquito ears in the swarm. Indeed, ablating efferent activity causes the onset of self-oscillations (SSOs) in males, which are believed to act as amplifiers of female wingbeats in the swarm and to be essential for its detection (7–9). We hypothesize that the efferent input and other potential neuromodulators adjusts the auditory performance of malaria mosquitoes for mating partner detection in the swarm. One of the most prominent candidates to modulate *An. gambiae* audition during swarm time is the efferent neurotransmitter octopamine. *An. gambiae* males present phonotactic responses towards female flight tones that are restricted to the circadian time when mosquitoes swarm and coincide in time with the erection of the antennal fibrillae (10), a phenomenon that is believed to increase the male sensitivity to female flight tones. In *An. stephensi* mosquitoes, fibrillae erection has been shown to be induced by octopamine (11). Octopamine is the counterpart of the vertebrate norepinephrine and plays similar functions to the noradrenergic system in invertebrates, modulating a plethora of insect behaviours and senses (12,13). It has also been shown to convey circadian clock information to insect sensory organs, modulating pheromone mating responses in moths (14–17) and attraction to conspecific secreted volatiles during the gregarious phase in locusts (18). Moreover, as octopamine signalling is restricted to invertebrates (12,19), it is a valid target for insecticide development (20–22). Indeed, octopamine receptors are currently the only type of G-protein coupled receptors (GPCRs) targeted by commercially available insecticides, although they are not used yet for mosquito control.

In this study, we profile the transcriptome of the malaria mosquito ear across the day to identify potential neuromodulators of auditory physiology. From the candidates identified, the α-adrenergic-like octopamine receptor *AGAP000045* shows a peak of expression during swarm time. We also identify a novel β-adrenergic-like octopamine receptor, *AGAP002886*, highly expressed in the malaria mosquito ear. We thoroughly investigate the auditory role of octopamine in the malaria mosquito *An. gambiae*. We find that octopamine modulates hearing mostly in male mosquitoes, with females showing lesser auditory changes after octopamine exposure. Our results suggest that in males octopamine plays multiple auditory roles; it controls the erection of the antennal fibrillae, sets flagellar stiffness and modulates auditory tuning, likely enhancing the male’s ability to detect the female in the swarm. Moreover, we determine that AGAP002886 is the primary octopamine receptor in the mosquito ear as mutant males completely fail to erect their fibrillae and present minimal changes in their audition upon octopamine exposure. We also show that the octopaminergic signalling in the malaria mosquito ear can be targeted by insecticides and is therefore a potential novel target for mosquito population control. Together, our results suggest that octopamine acts as an important modulator of hearing in malaria mosquitoes, tuning their auditory physiology to the acoustic environment of the swarm, thus facilitating the detection of mating partners.

## Results

### Transcriptomics suggest a complex neuromodulatory control of the mosquito audition

Mosquito ears are one of the most complex sensory organs in insects and are composed of a sound receiver, the antennal flagellum, and the auditory organ *per se*, the JO (1). To characterize the neuromodulatory network of the malaria mosquito ear and its potential implication in the regulation of the auditory physiology during swarm time, we undertook RNA-sequencing analysis of male and female ears at six different time points along the day (lights on or ZT0, ZT4, ZT8, swarm time or ZT12, ZT16 and ZT20). We collected exclusively second antennal segments (hosting JO) without flagella, to avoid ‘pollution’ from chemosensory neurons located in the flagellum (Fig.1a). Identified transcripts (Supplementary Tables 1 and 2) were assigned to GO accession numbers and those with molecular function potentially related to neuromodulation were selected (including biogenic amine, classical neurotransmitter and neuropeptide receptors, as well as other GPCRs or receptors related to other sensory modalities). Within these categories, we identified 173 and 152 genes expressed in the male and female JOs, respectively (Fig. 1c, d; Table 1; complete dataset in Supplementary Tables 3 and 4).

**Table 1:**
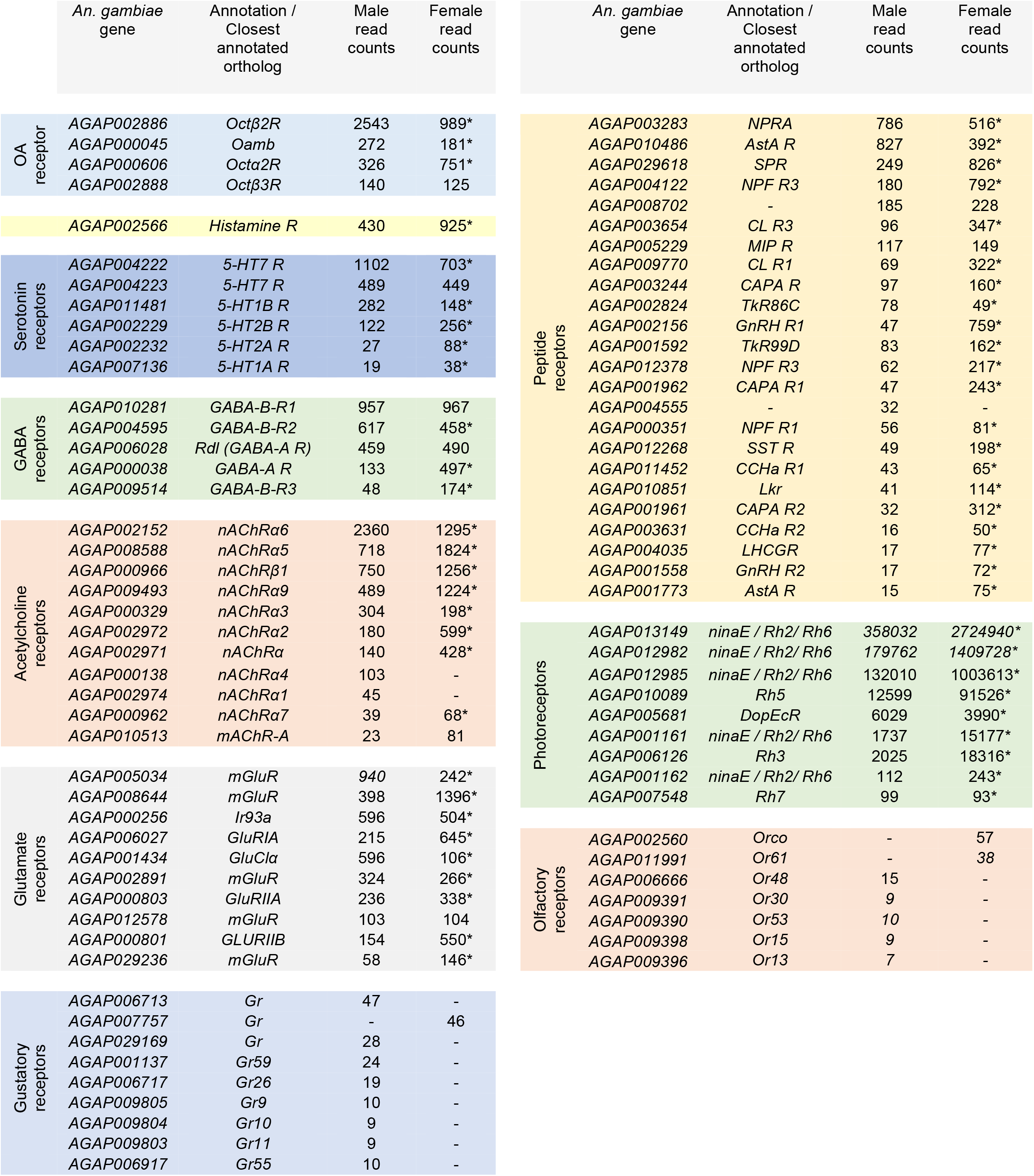
Genes expressed in the male and female JO potentially related to auditory neuromodulation, including receptors for octopamine, histamine, serotonin, GABA, acetylcholine, glutamate and peptides. Receptors involved in other sensory modalities (olfaction, gustation and photorreception) are also shown. Only genes with higher read counts in each category are shown (complete dataset in Supplementary Tables 3 and 4). Read count corresponds to the normalized average across all time points of sample collection. * in the last column indicates that the gene was differentially expressed between males and females, p < 0.05. OA: octopamine.

**Fig. 1:**
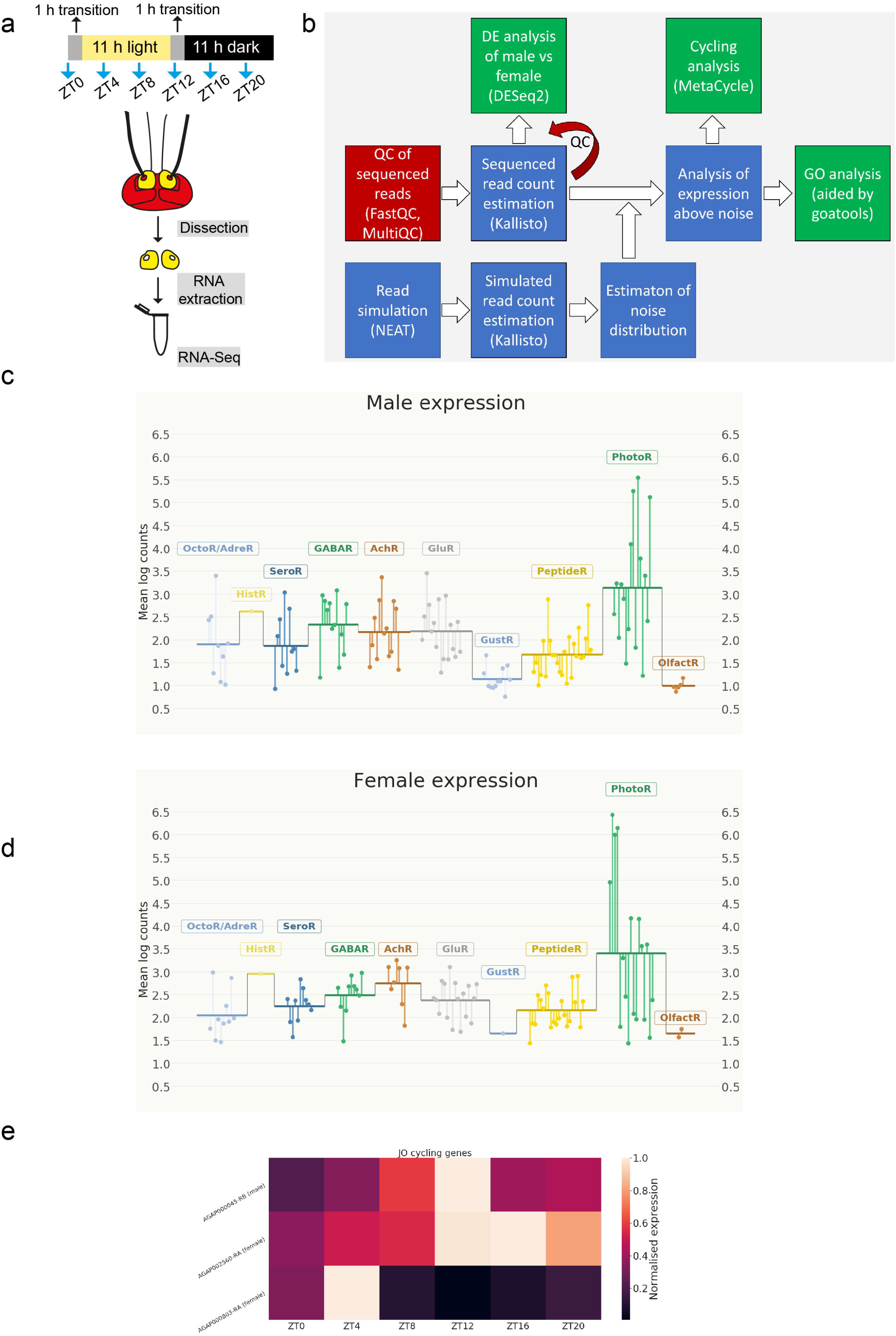
Transcriptomics of potential neuromodulatory genes in the JOs of male and female *An. gambiae* mosquitoes. **a)** Sample collection and preparation for RNA-sequencing. Mosquitoes were entrained to 12 h: 12 h light/dark cycle with one hour light transition between light and dark phases. Male and female mosquito were collected every four hours, their ears dissected and RNA extracted for RNA-sequencing. **b)** Schematic workflow of the analysis of RNA-sequencing data. **c, d**) Plots summarising the expression levels of transcripts belonging to the selected GO categories in males (**c**) and females (**d**) potentially involved in neuromodulation. The transcriptomic data support the existence of an extensive neuromodulatory network acting on the mosquito JO. Each point represents a transcript and depicts the average of the log counts of the ZT point showing the highest expression for that gene. Horizontal lines represent the mean of the log counts represented by each point within a GO category. **e**) Heatmap representing normalised expression per time point of genes showing cycling expression in males and females. A single gene in males (AGAP000045) and two genes in females (AGAP002560 and AGAP00803) show cyclic expression in the mosquito ear.

Biogenic amines are important sensory organ modulators in insects (17). Our results support that this is also the case in the malaria mosquito ear as several biogenic amine receptors were expressed in both sexes (Fig. 1c,d). An ortholog of the octopamine β2 receptor in *Drosophila, AGAP002886*, was the highest expressed biogenic amine receptor. Other α and β octopamine receptors were also expressed (*AGAP000045, AGAP000606, AGAP002888*). Several serotonin receptor orthologs (including 5-HT1a, 5-HT1b, 5-HT2a and 5-HT7 receptor orthologs) were also expressed in both male and female JOs. A histamine receptor ortholog (*AGAP002566*) was also identified. Most biogenic amine receptors were differentially expressed in males and females (Table 1, Supplementary Table 5).

We also examined the expression of classical neurotransmitter receptors. GABA acts as efferent neurotransmitter in the ear of *Cx. quinquefasciatus* mosquitoes (6). Multiple GABA receptors were expressed in both sexes, consistent with the extensive GABAergic innervation observed in the auditory nerve of *An. gambiae* (unpublished *data, not shown*). Three metabotropic GABA-B receptor orthologs (*AGAP010281, AGAP004595, AGAP009514*) were highly expressed in both sexes. The ionotropic GABA-A receptor *Rdl, AGAP006028*, which causes resistance to the insecticide dieldrin in *Anopheles* populations (23), was also identified. We also found several nicotinic acetylcholine receptor alpha subunits and a single beta subunit. Interestingly, the expression of nicotinic acetylcholine receptor subunits was higher in females (Table 1). Nicotinic acetylcholine receptors are critical components of the efferent auditory system in vertebrates (24). Moreover, a single muscarinic acetylcholine receptor was found. Several glutamate receptors were also identified. Metabotropic glutamate receptors (mGluR) were highly expressed and different ionotropic (including AMPA -*AGAP006027*- and kainate receptor -*AGAP000803*-orthologs) were also found. An ortholog of *Drosophila* IR93a glutamate receptor (*AGAP000256*), involved in thermo- and hygrosensation (25), was also identified.

Neuropeptides act as important neuromodulators of insect sensory neurons (26). Our dataset exhibited a broad repertoire of neuropeptide receptors in both sexes (Table 1). Orthologs of the natriuretic peptide, the sex peptide, calcitonin and allatostatin 3 receptors were highly expressed in both sexes. We also identified putative tachykinin receptors 1 and 2, which have been implicated in the modulation of olfactory neurons in *Drosophila* (27), as well as several neuropeptide F receptor orthologs. Neuropeptide F modulates the locomotor plasticity of swarming migratory locusts (28). Two putative gonadotrophin-releasing hormones receptors were expressed in both sexes, although interestingly the expression was higher in females. Other orthologs of neuropeptide receptors also found included crustacean cardioactive peptide, leukokinin, CCHamide-1, CCHamide-2, allatostatin 2, capability and myosuppressin and pirokinin receptors.

We also studied the expression of receptors involved in other sensory modalities to investigate modulatory roles of auditory responses following unconventional signalling pathways (29). Visual rhodopsins were particularly highly expressed, particulary in the female JO. Rhodopsins have been previously shown to mediate auditory and mechanosensory roles in *Drosophila* (30,31). Some gustatory and olfactory receptors were also expressed, yet those mostly in males.

As we are particularly interested in the auditory modulation at swarm time, we interrogated our dataset for genes whose expression cycled in the 24-hour period of data collection and identified one gene in males and two genes in females with rhythmic expression. Interestingly, these genes differed between sexes (Fig.1e). In males, the octopamine alpha receptor *AGAP000045*, ortholog of the OAMB receptor in *Drosophila* (32), showed cyclic expression that peaked at ZT12, the laboratory swarm time. In females, an ortholog of a kainate glutamate receptors, *AGAP000803*, showed a rhythmic expression that peaked at ZT12. The expression of the Orco receptor AGAP002560 was also cyclic in females, but higher during the day, and lower at night (33,34). A list of all genes cycling in the male and the female JO (independently of the GO term) can be found in Supplementary Tables 6 and 7.

### Octopamine modulates malaria mosquito audition in a time-dependent, diel manner

Our transcriptomic analysis suggested a high plasticity of the mosquito auditory physiology and a primary role of octopamine as auditory modulator at swarm time. We explored the auditory role of octopamine by injecting octopamine in the mosquito thorax and quantifying resulting changes in auditory function (Fig. 2). We used two different octopamine concentrations (1 mM and 10 mM) to explore sensitivity and dynamic range differences. We repeated the tests at two different circadian zeitgeber times (ZTs); during a phase of inactivity for *An. gambiae* mosquitoes (ZT4 or four hours after lights-on) and during the laboratory swarm time (ZT12 or the laboratory dusk) (Fig. 2). Control injections were performed using a ringer solution.

**Fig. 2:**
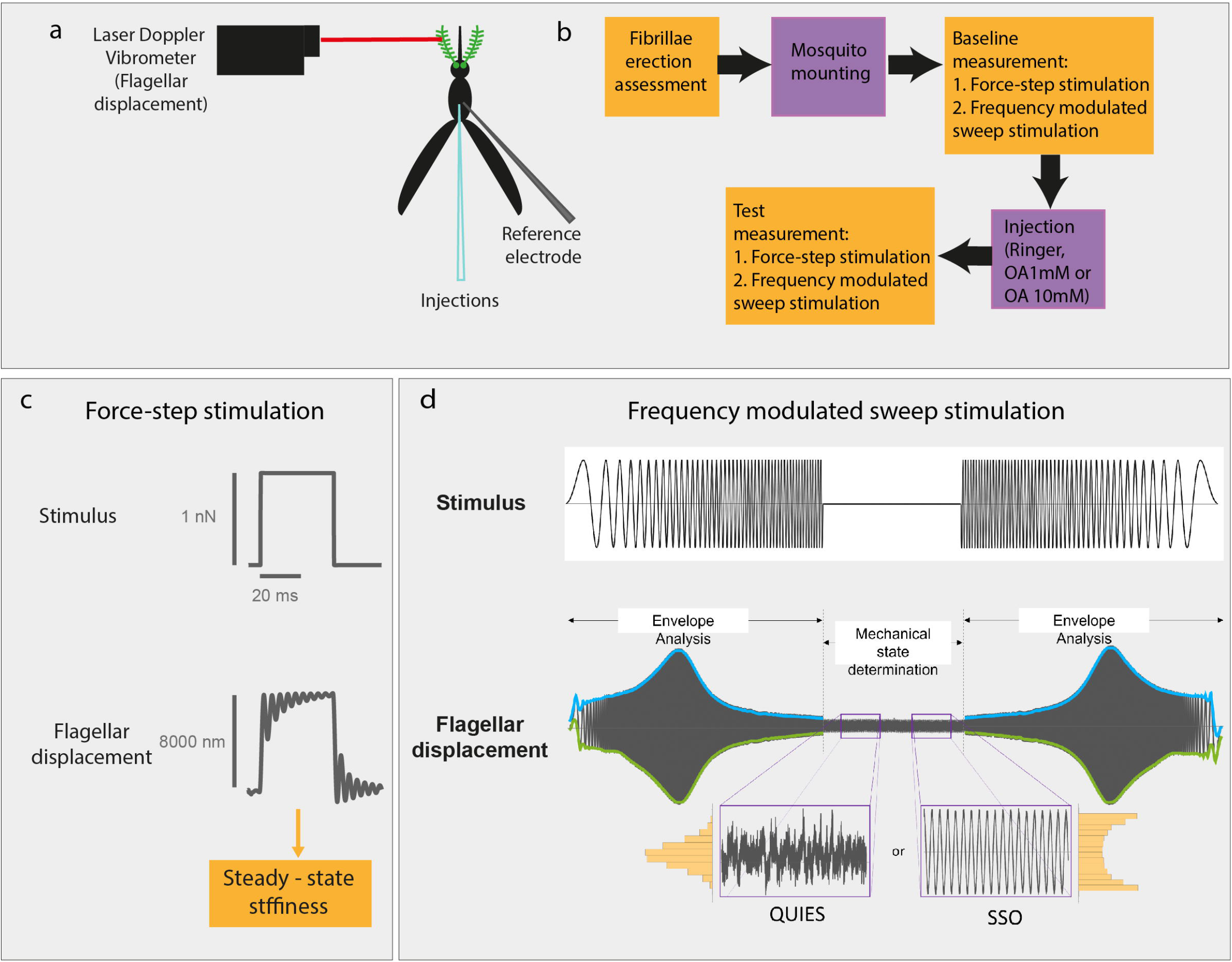
Experimental setup and analysis pipeline for auditory analysis. **a)** Experimental paradigm of laser Doppler vibrometry (LDV) recordings. **b)** Auditory tests working pipeline. **c)** Force-step stimulation and flagellar responses. d) Frequency-modulated sweep stimulation and flagellar responses.

### Octopamine causes erection of male antennal fibrillae

A well-characterised circadian clock signature of *An. gambiae* audition is the rhythmic pattern of erection of the fine hairs – or fibrillae-that cover the male antennal flagellum (35) and that become erected before swarm time. The extension of the antennal fibrillae is believed to assist the acoustic detection of mating partners in the swarm by increasing the flagellum’s effective acoustic surface and thus the male’s sensitivity to female wingbeats (36). Octopamine has been shown to induce fibrillae erection in *An. stephensi* mosquitoes (11). We tested whether this was also the case in *An. gambiae*.

A visual assessment of the fibrillae state of mosquitoes kept in the glass vials inside circadian incubators confirmed that 100% of male mosquitoes presented erected fibrillae at ZT12 and 0% did at ZT4 (Fig. 3a). After mounting the mosquitoes in preparation for auditory tests, and likely because of gluing the JO base for auditory recordings, all mosquitoes collapsed the fibrillae, irrespective of the circadian time. This allowed us to study the effects of octopamine from a baseline state in which all mosquitoes (both at ZT4 and ZT12) had collapsed fibrillae. Injecting 1 mM octopamine caused the full erection of the antennal fibrillae in 33% mosquitoes at ZT4 and 62.5% at ZT12. These results suggest circadian time-dependent changes in the octopamine effects. Increasing the octopamine concentration to 10 mM caused increased rates of fibrillae erection (60% of males at ZT4 and 83% at ZT12) while control ringer injections caused erection in 17% mosquitoes at ZT4 and 0% at ZT12.

**Fig. 3:**
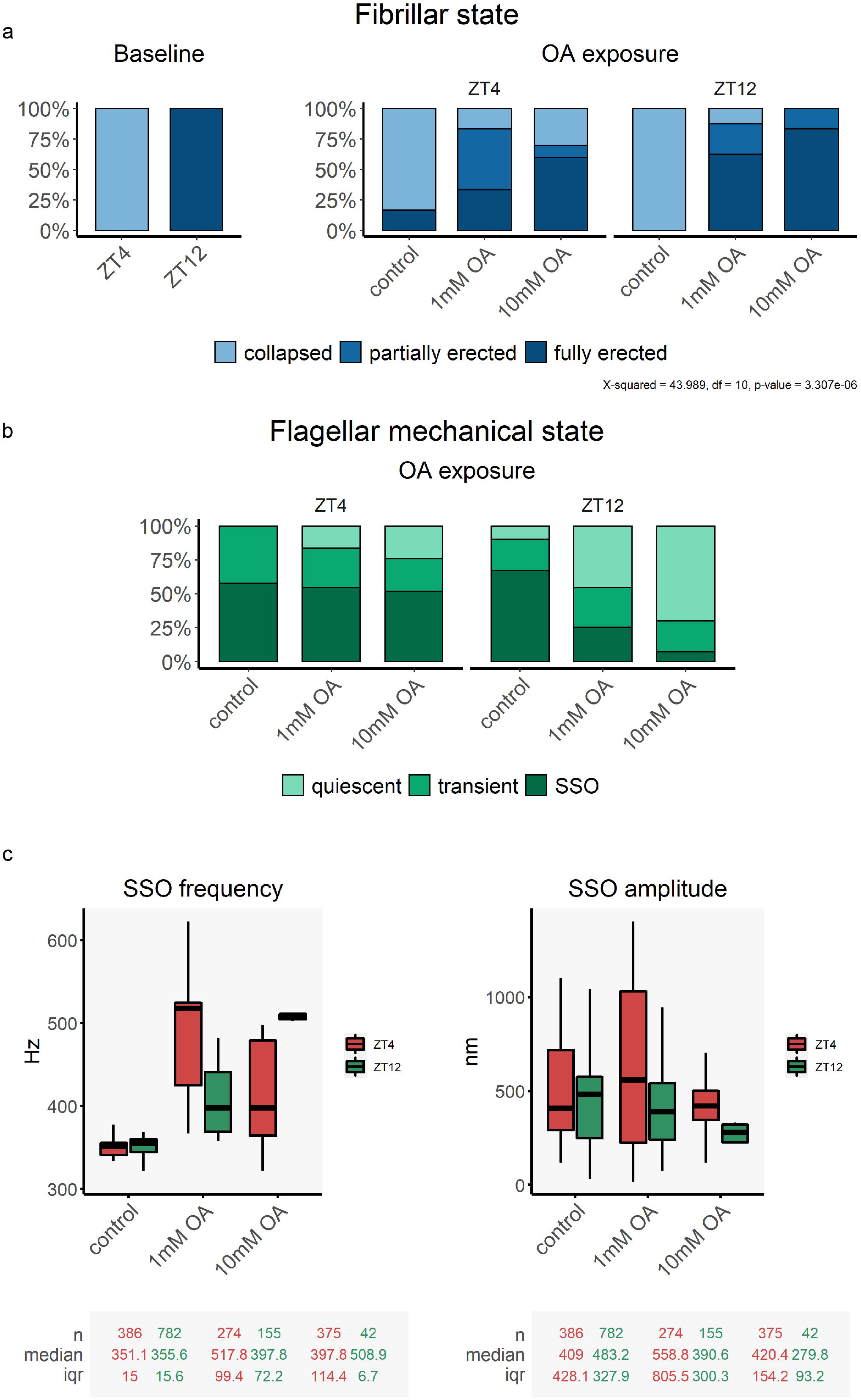
Octopamine injection causes the erection of the antennal fibrillae and influences the flagellar mechanical state in male *An. gambiae* mosquitoes. **a**) Fibrillae erection state under different experimental conditions. Left panel, fibrillae erection baseline levels before mosquito mounting. At ZT12, during the laboratory swarm time, all male mosquitoes presented erected fibrillae, while at ZT4 -when *An. gambiae* mosquitoes are inactive-, all mosquitoes presented collapsed fibrillae. Right panel, fibrillae erection state upon octopamine injections. Please note that as a result of the mounting process, all mosquitoes collapsed their fibrillae, so prior to injecting octopamine all mosquitoes had collapsed fibrillae. Injecting both 1 mM and 10 mM octopamine caused the erection of the antennal fibrillae. **b**) Flagellar mechanical state based on the analysis of unstimulated free fluctuations of the flagellum. At ZT4, only mosquitoes exposed to octopamine will present quiescent flagella. At ZT12, some quiescent mosquitoes can be observed in the control, and octopamine exposure causes a shift to quiescent states and a sharp reduction in the amount of SSO mosquitoes. **c**) SSO frequency and amplitude in different experimental conditions. Tables underneath each graph display the n numbers, median and interquartile range (iqr) for each of the categories shown in the graph above. Red boxplots show ZT4 data, and green ZT12. Octopamine injections caused a shift in the SSO frequency to higher values and a decrease in the amplitude of the oscillations. OA: octopamine, SSO: self-sustained oscillations.

### Octopamine prompts a quiescent state in the male mosquito flagellum

We first analysed the sound receiver (flagellum) behaviour of males in unstimulated conditions (free fluctuations). Male mosquito sound receivers have been previously reported to exhibit two different mechanical states. One of them presents spontaneous, large and mono-frequent self-sustained oscillations (SSOs) of around 350 Hz and with velocity amplitudes around 1mm/s, which is ∼1000-fold above baseline levels (7). By contrast, quiescent receivers show best frequencies of ∼500 Hz and substantially lower velocity amplitudes of ∼ 1µm/s. Although the ecological relevance of these two states is not fully understood, SSOs seem to act as amplifiers of the female wingbeats in the swarm (4,5) and thus likely to be relevant for partner detection. We speculated that the time of the day would influence the flagellar mechanical state.

We first developed an analytical framework to provide a quantitative definition of the different mechanical states (see Methods). We analysed the free fluctuations of the flagellum and appreciated that apart from being either in quiescent or SSO state, mosquitoes could also display a transient state in between the two extremes (Fig. 3b). During the inactivity phase ZT4, mosquitoes displayed either SSO or were in a transient state, but none was quiescent ([ZT4]: 0% quiescent, 42.5% transient, 57.5% SSO). Instead, during swarm time, the mechanical state distribution changes: there were fewer transient mosquitoes, and more displaying SSOs or being quiescent ([ZT12]: 9.9% quiescent, 23.2% transient, 67% SSO). Our data support that the flagellar mechanical state changes across the day.

We examined changes in the mechanical state caused by octopamine exposure. Interestingly, injecting 1 mM octopamine at ZT4 pushed transient mosquitoes into a quiescent state, while the proportion of SSO mosquitoes remained unaltered (Fig. 3b). Injecting 10 mM octopamine caused a more pronounced effect ([ZT4]; OA1mM: 16.5% quiescent, 28.9% transient, 54.6% SSO; OA10mM: 24.3% quiescent, 24% transient, 51.7% SSO). By contrast, at ZT12, octopamine exposure induced a shift from SSO to quiescent state, as there was an observed decrease in the number of mosquitoes displaying SSO and a concurrent increase in the proportion of quiescent mosquitoes ([ZT12]; OA1mM: 45.5% quiescent, 29.3% transient, 25.2% SSO; OA10mM: 69.8% quiescent, 22.7% transient, 7.5% SSO). We conclude that octopamine drives the flagellum into a quiescent state, and that this effect is circadian-time dependent. While at ZT4 octopamine affects only transient mosquitoes, at ZT12 the sensitivity to octopamine is higher and hence also SSO mosquitoes are pushed into quiescent states.

### Octopamine causes an increase in SSO frequency

The vibrations of unstimulated flagella, i.e. their ‘free fluctuations’ have previously been used to assess the frequency tuning of the mosquito ear (7). Free fluctuation recordings were used to extract the oscillator frequency, *f*_*0*_, and displacement amplitudes of the flagellum’s spontaneous vibrations. We first explored potential circadian time-dependent changes in free fluctuation values (Fig. 3c, Supplementary Figure 8 and 9). Control injections revealed small, yet significant circadian time-dependent changes in both the SSO frequency and amplitude, which were higher at ZT12 compared to ZT4 (Fig. 3c, [ZT4]; control, Frequency: 351 ± 4 Hz, Amplitude: 409±230 nm; [ZT12]; control, Frequency: 356 ± 7 Hz, Amplitude: 483±119 nm). 351+-15

Octopamine injections caused an overall increase of the oscillator frequency and a decrease of the amplitude of the SSOs (Fig. 3c) (quiescent mosquitos do not produce sinusoidal or periodic signals and a frequency could not be conventionally determined within the resolutions of our paradigms. For this reason, they are not represented here, see Methods). This effect was more pronounced at ZT12 for 10 mM octopamine injections where the average SSO frequency values were above 500 Hz, compared to around 350 Hz for controls. The damping effect of octopamine on the SSO amplitude was also stronger at ZT12 upon 10 mM octopamine injections, showing values approximately half of control amplitude values. The stronger responses to octopamine at ZT12 depicted both an increase in the sensitivity to octopamine and in the dynamic range of octopamine effects in the free fluctuations during the laboratory swarming time ZT12.

### Octopamine affects the auditory tuning of stimulated male mosquito ears

The flagellar sound receiver acts as an inverted pendulum, which vibrates as a forced damped harmonic oscillator in response to the sound-induced motion of surrounding air particles produced by the wingbeats of mating partners (37). To study how octopamine affected flagellar responses, we stimulated the flagellum using frequency-modulated sweeps (0-1000 Hz to cover the frequency range of *An. gambiae* male and female flight tones) and measured flagellar displacements. We fitted a forced damped harmonic oscillator model to the resulting flagellar responses and extracted different biophysical parameters to quantify flagellar biomechanics. We studied both quiescent receivers and receivers undergoing SSOs.

We first analysed potential circadian time-dependent changes of these biophysical parameters. We did not observe clear circadian time-dependent differences in flagellar responses to sweep stimulation between the control cases of ZT4 and ZT12 (Fig. 4a-d, Supplementary tables 10 and 11). However, octopamine exposure affected the oscillator at multiple levels and these changes were circadian-time dependent overall (Fig. 4a-d). In quiescent animals, octopamine exposure caused a decrease in the acceleration, an increase in the oscillator frequency and a decrease in the damping ratio. These effects were circadian-time dependent, and overall, stronger at ZT12 upon exposure to 10mM octopamine, indicating higher sensitivity and dynamic range during the laboratory swarm time.

**Fig. 4:**
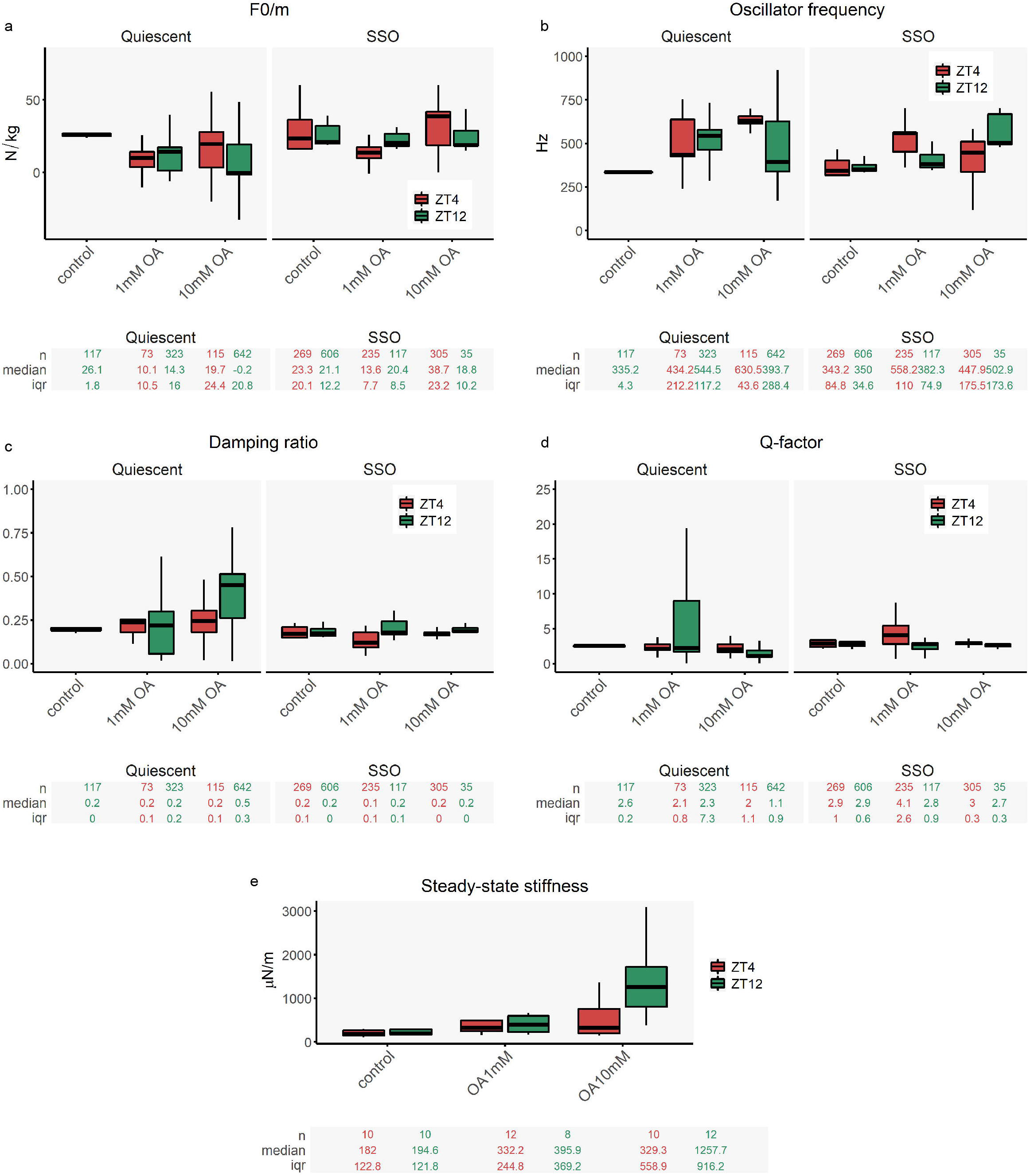
Octopamine injection causes acute changes in auditory responses of male malaria mosquitoes. Auditory tests were conducted after injecting three different solutions: control ringer and two different octopamine concentrations (1 mM and 10 mM). Red boxplots show ZT4 data, and green ZT12. (**a-d**) Biophysical parameters were extracted from fitting damped harmonic oscillator functions to the flagellar responses to frequency-modulated sweeps, including acceleration (F0/m), the oscillator frequency, the damping ratio and Q-factor. Flagella were assigned to either displaying a quiescent state or SSOs. No quiescent mosquitoes were observed at ZT4 in control conditions, so only one boxplot is shown. Tables underneath each graph display the n numbers, median and interquartile range (iqr) for each of the categories shown in the graph above. Octopamine exposure affected most biophysical parameters, and remarkably, a sharp increase in the oscillator frequency. **e**) Steady-state stiffness values extracted from force-step stimulation responses. Octopamine injections induced an increase in flagellar stiffness values that was acuter during swarm time ZT12.

Octopamine caused fewer changes in the biophysical parameters of SSO flagella, except for the oscillator frequency that strongly shifted to higher frequencies upon octopamine exposure. The shift was strongest at ZT12 after 10mM octopamine injections ([ZT4]; control: 343± 28Hz; OA10mM: 448 ± 76Hz; [ZT12]; control: 350 ± *11* Hz; OA10mM: *5*0*3* ± *11* Hz).

### Octopamine causes an increase in the flagellar stiffness of male mosquitoes

Changes in flagellar steady-state stiffness upon octopamine injections were analysed as an indication of the ear’s sensitivity to mechanical stimulation. Flagellar steady-state stiffness were calculated from force-step a ctuations (7). In short, the flagellar steady-state stiffness describes the force required to hold the flagellum a t a certain steady-state displacement (‘holding stiffness’), it is indicative of the flagellum’s ‘passive mechanics’. Injecting 1mM octopamine caused a sharp increase in flagellar stiffness (Fig. 4e, Supplementary tables 1 2 and 13). The effect was circadian time-dependent and strongest at ZT12 upon 10mM octopamine injections, where stiffness values were 7-fold higher than for control injections ([ZT4]; control: 182 ± 41 μN/m; OA1 mM: 332.2±105.3 μN/m; OA10mM: 329.3±149.8 μN/m; [ZT12]; control: 194.6 ± 106.2 μN/m; OA1mM: 395. 9±129.7 μN/m; OA10mM: 1257.7±250.6 μN/m).

### Octopamine exposure has milder effects on female *Anopheles gambiae* mosquito audition

Our transcriptomic data showed expression of octopamine receptors also in female ears, although the expression did not cycle along the day (Table 1, Fig. 1e). To examine the effect of octopamine exposure in female auditory function, we examined the responses to frequency-modulated sweep and force-step stimulation. At the baseline, there were some circadian time-dependent differences in the responses to sweep stimulation: the oscillator frequency was lower at ZT12, while the damping ratio was higher (Fig.5a-d). Overall, octopamine exposure caused small changes in the biophysical parameters studied, and the oscillator frequency did not change during the laboratory swarm time between a control and a 1mM octopamine injection. By contrast, flagellar stiffness values extracted from force-step stimulation changed between control and octopamine injections (Fig. 5e, Supplementary tables 14-17).

**Fig. 5:**
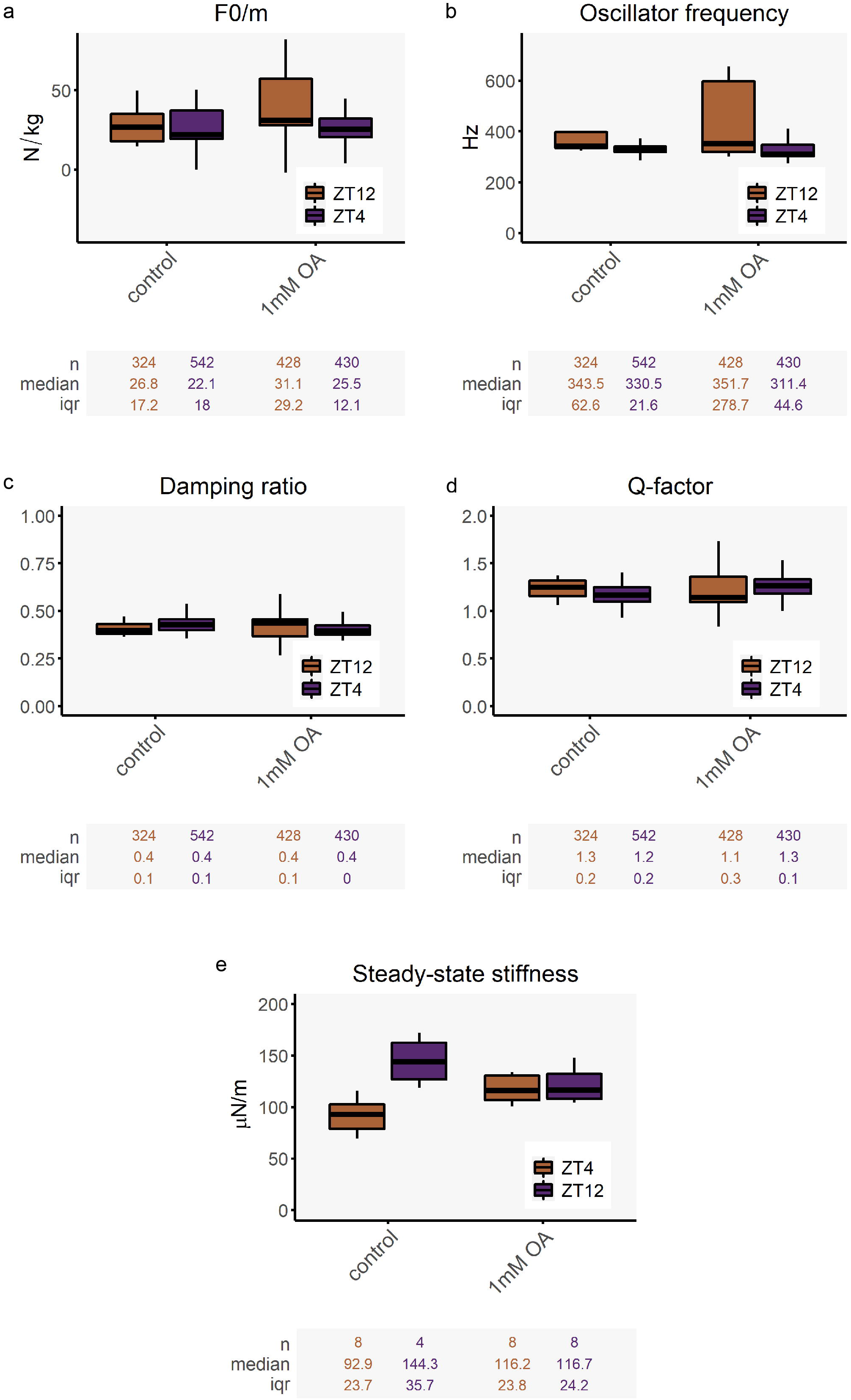
Octopamine injection causes mild effect in female malaria mosquitoes. Tests of auditory biophysics were conducted after performing injecting two different solutions: control ringer and 1 mM octopamine concentrations. Orange boxplots show ZT4 data, and purple ZT12. (**a-d**) Biophysical parameters were extracted from fitting damped harmonic oscillator functions to the flagellar responses to frequency-modulated sweeps, including acceleration (F0/m), the oscillator frequency, the damping ratio and Q-factor. Flagella were assigned to either displaying a quiescent state or SSOs. The tables underneath each graph display the n numbers, median and interquartile range (iqr) for each of the categories shown in the graph above. Octopamine exposure did have very mild effects on female mosquito hearing and the oscillator frequency did not change upon octopamine injections. **e**) Steady-state stiffness values extracted from force-step stimulation responses. Octopamine injections did not cause any change in flagellar stiffness in malaria mosquito females.

### AGAP002886 and AGAP000045 as novel octopamine receptor mediating octopamine auditory roles in malaria mosquitoes

To gain insight into octopamine auditory function, we generated knock-out mosquitoes of two octopamine receptors expressed in the mosquito ear. We selected the most highly expressed octopamine receptor, A*GAP002886*, and the receptor showing a peak of expression during swarm time, *AGAP000045*. We used CRISPR/Cas9 genome editing to target regions of predicted octopamine binding or GPCR activity and highly conserved across mosquito strains (1^st^ coding exon for AGAP0000045 and 3^rd^ coding exon for AGAP002886) (32,38,39). The presence of a GFP fluorescent marker inserted in the targeted site allowed for tracking mutant alleles (Fig. 5a). Both mosquito mutant lines were tested for the unstimulated and stimulated auditory behaviours.

### Octopamine receptor mutants present severe defects in the pattern of fibrillae erection

We first observed the fibrillae erection pattern of the octopamine receptor mutants compared to wild-type animals during the laboratory swarm time. As shown above, 100% of wild-type males erected their fibrillae at ZT12 (Fig 3a). Almost 90% of *AGAP000045* homozygous mutant males also did so. By contrast, none of *AGAP002886* knock-out animals erected their fibrillae during the laboratory swarming time. We then performed 1 mM octopamine injection after mounting the mosquitoes (mounting the mosquitoes causes fibrillae collapse in all mosquitoes). Although octopamine exposure caused the partial or full erection of the antennal fibrillae in almost 90% of wild-type males at swarming time, it did not cause any erection in *AGAP002886* mutants and only 50% of *AGAP000045* knock-outs responded to it (Fig. 6b).

**Fig. 6.**
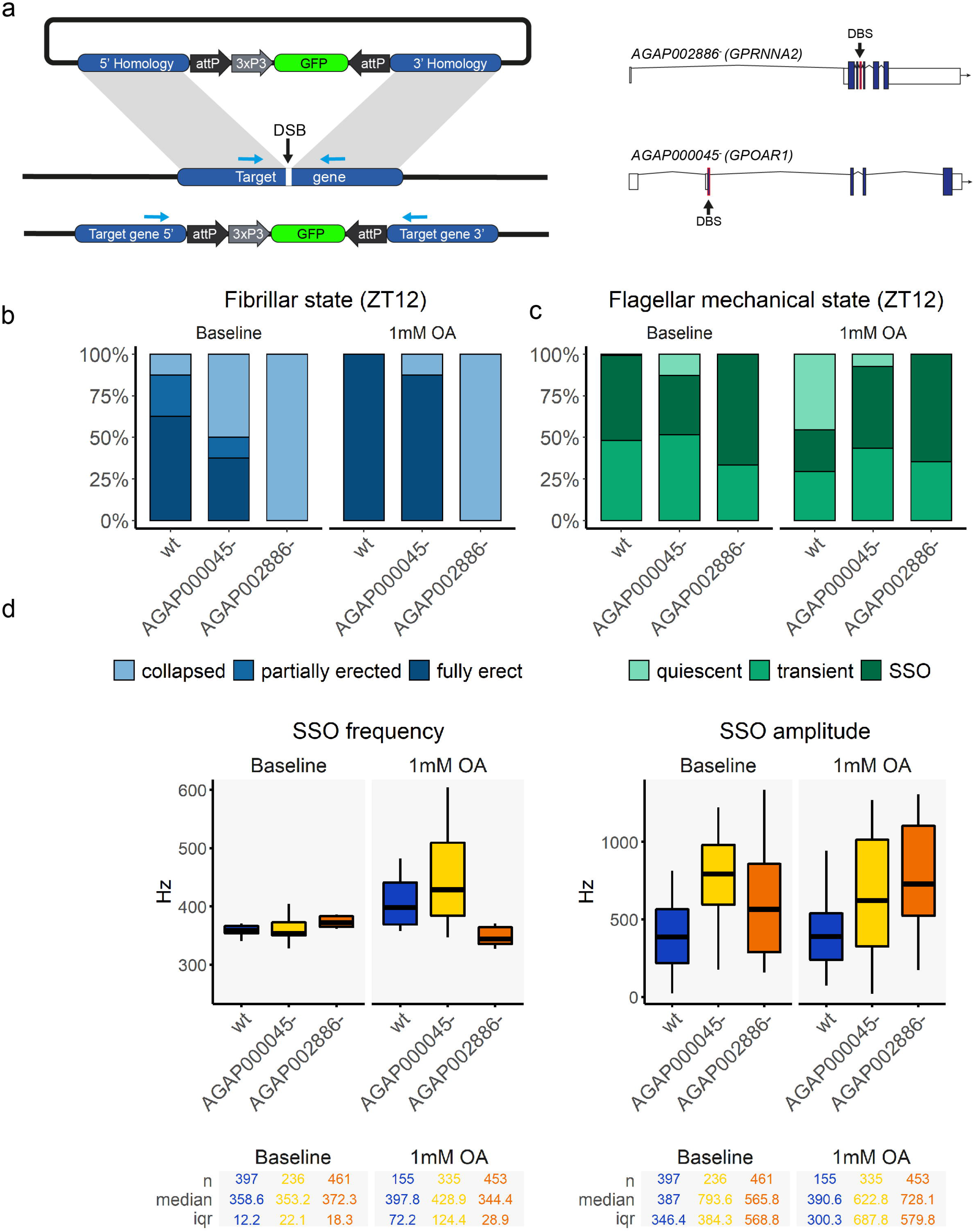
Auditory behaviour of unstimulated ears in octopamine receptor mutants. (**a**) Schematic of the knock-out generation approach. A double-strand break (DSB) was introduced into specific locations of the target gene using CRISPR-Cas9 mediated cleavage. A cassette containing an attP-flanked 3xP3::GFP marker construct was inserted into the target gene DBS site by homology directed repair. GFP expression was used to screen for transformants. The DSB site was located in the third exon of the receptor *AGAP002886* and the first exon of the receptor *AGAP000045*. (**b**) Fibrillae erection state in octopamine receptor mutants compared to wildtypes. Baseline levels before mounting the mosquitoes, show that none of AGAP002886 knock-out mosquitoes erect their fibrillae during the laboratory swarm time, while almost all wildtype mosquitoes erect them and half of *AGAP000045*^*-*^ mosquitoes. Upon octopamine exposure, *AGAP002886*^*-*^ mosquitoes keep the fibrillae collapsed, while almost all wildtype and *AGAP000045*^*-*^ animals erect them. **c**) Flagellar mechanical state based on the analysis of unstimulated free fluctuations of the flagellum during swarm time. At the baseline, almost all mosquitoes exhibit exclusively SSO (except for a few *AGAP000045* mutants). Octopamine exposure pushes around half of wildtype flagella to a quiescent state, while the mutants do not change the flagellar mechanical state and maintain the SSOs. **d**) SSO frequencies and displacement amplitudes in the different genotypes during swarm time. The tables underneath each graph display the n numbers, median and interquartile range (iqr) for each of the categories shown in the graph above. Baseline levels show an increase SSO frequency in *AGAP002886* mutants, but these mutants do not increase the oscillator frequency upon octopamine exposures. *AGAP000045*^*-*^ animals respond with a sharper increase in the SSO frequency values than wildtypes after octopamine injections. OA: octopamine, SSO: self-sustained oscillations.

### Octopamine receptor mutants do not shift to quiescent flagellar states upon octopamine injections

We then studied the mechanical state of unstimulated flagella in mutant animals during the laboratory swarm time (ZT12). Data are shown only for SSO animals, as only very few *AGAP000045* mutants and none of the *AGAP002886* mutants exhibited quiescent states of their flagella. At the baseline, all genotypes presented a similar distribution, with most animals either presenting SSOs or a transient state (Fig. 6c). Although in wild-type animals octopamine prompts a shift to quiescent states, this was not the case for any of the mutants that maintained the SSO or transient states despite being exposed to octopamine. Regarding the SSO frequency (Fig. 5d), *AGAP002886* mutants exhibited SSO of higher frequency compare to wild-type animals at the baseline ([wt], baseline: 358.6±12.2 Hz; [*AGAP000045*^*-*^], baseline: 353.2±22.1 Hz; [*AGAP002886*^*-*^], baseline: 372.3±18.3 Hz; p-value [wt] vs [*AGAP002886*^*-*^] <0.05). Octopamine exposure caused a shift to higher frequencies in the SSO of both wild-type and *AGAP000045*^*-*^ animals, being the effect stronger for the mutants. By contrast, *AGAP002886* mutants reduce the SSO frequency upon octopamine injections. The SSO amplitude was higher at the baseline for both octopamine receptor mutants compared to wild-type mosquitoes (Fig. 5 d). Although octopamine did not induce changes in the SSO amplitude in wild-type mosquitoes, it did affect them in both mutants.

We subjected the octopamine receptor knock-outs to frequency-modulated sweep stimulation. At the baseline, wild-type and mutant animals presented differences in all parameters studied, except the damping ratio between wild-type and *AGAP000045*^*-*^ (Fig.7a-d, Supplementary tables 18-19). Of interest, mutant mosquitos exhibited higher oscillator frequency than wild type ([wt], baseline: 344.4±41.3 Hz; [*AGAP000045*^*-*^ ], baseline: 369.1±23.9 Hz; [*AGAP002886*^*-*^], baseline: 397.6±22.8 Hz; p-value<0.05). Injecting 1 mM octopamine caused distinct effect in both mutant lines. In *AGAP000045* mutants, the effects of octopamine exposure are more extreme than in wild-type animals (Fig.7a-d). The shift of the oscillator frequency and the acceleration to higher values is more pronounced. By contrast, the shift in the parameters values is less pronounced for *AGAP002886* knock-outs and, opposite to what is observed for the other phenotypes, the oscillator frequency decreases upon octopamine injections.

**Fig. 7.**
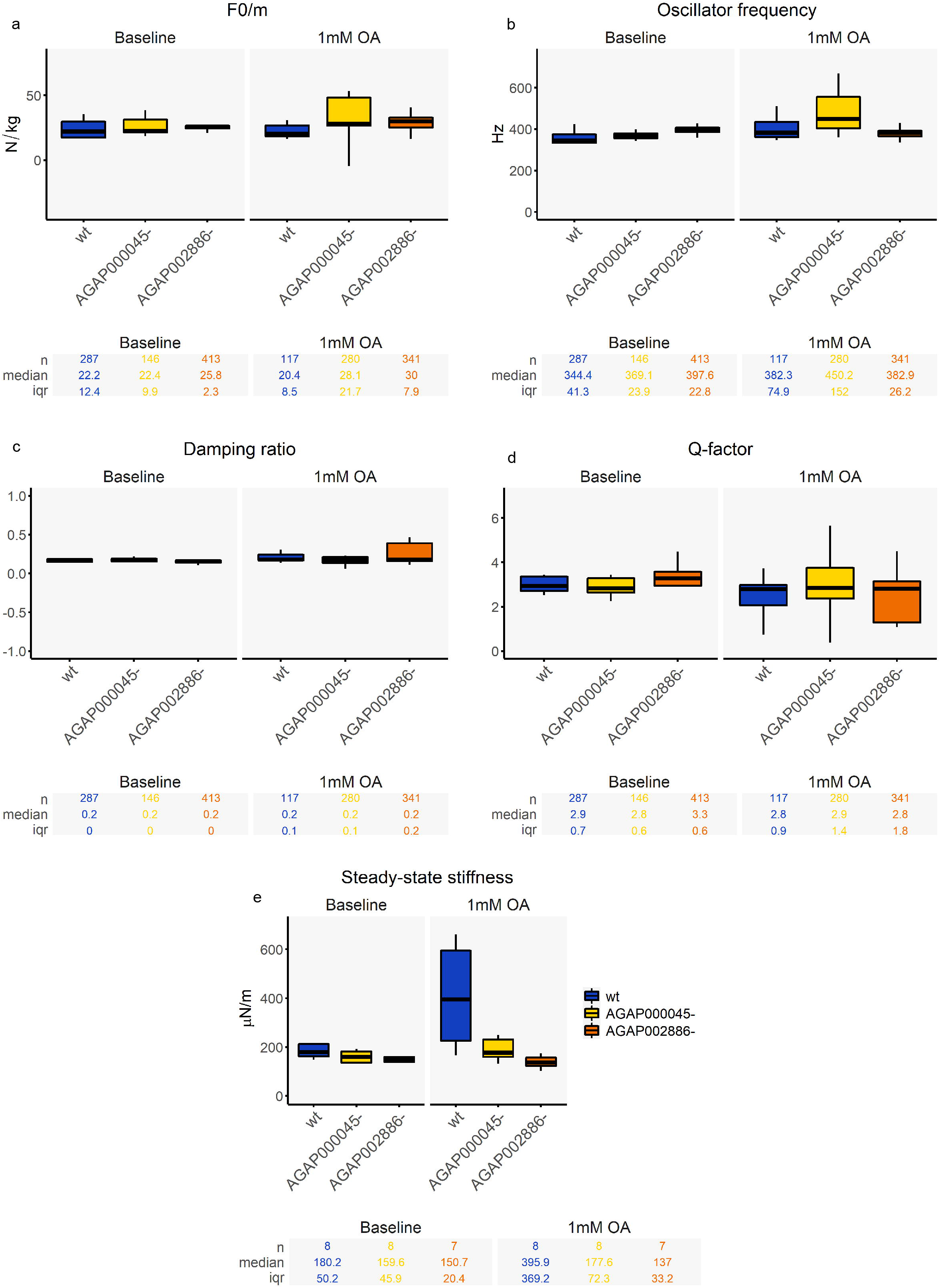
Different phenotypes in the auditory stimulated responses of octopamine receptor mutants upon octopamine exposure. Auditory tests were conducted after injecting two different solutions: control ringer and 1 mM octopamine concentrations. Auditory tests were only performed at ZT12 (**a-d**) Biophysical parameters were extracted from fitting damped harmonic oscillator functions to the flagellar responses to frequency-modulated sweeps, including acceleration (F0/m), the oscillator frequency, the damping ratio and Q-factor. As most mutant flagella were constantly exhibiting SSO, only SSO values are shown. Tables underneath each graph display the n numbers, median and interquartile range (iqr) for each of the categories shown in the graph above. Octopamine exposure affected most biophysical parameters in *AGAP000045* mutants, but the effects on AGAP002886 knock-out were very mild. Interestingly, the changes in values in *AGAP000045* mutants were stronger than in wildtype animals (we are only showing results for SSO mosquitoes). The increase in the oscillator frequency of *AGAP000045* mutants reached values of up to 450 Hz. **e**) Steady-state stiffness values extracted from force-step stimulation responses. The increase in the stiffness values was very faint in octopamine receptor mutants compared to wildtypes. In *AGAP002886* knock-outs the stiffness values decreased upon octopamine injections. OA: octopamine.

Responses to force-step stimulation were also analysed to extract steady-state stiffness values (Fig. 7e, Supplementary tables 20-21). At the baseline, *AGAP002886* mutants presented lower values compared to both the wild-type and *AGAP000045* knock-outs ([wt], baseline: 180.2±50.2 μN/m; [*AGAP000045*^*-*^], baseline: 159.6±45.9 μN/m; [*AGAP002886*^*-*^], baseline: 150.7±20.4 μN/m; p-value<0.05). Although wild-type animals showed a striking increase in the stiffness upon octopamine exposure, this was not the case for neither of the mutans that did not significantly change their stiffness values.

### The AGAP002886 mediated octopaminergic signalling in the mosquito ear can be targeted for mosquito control

Octopamine receptors are promising insecticide targets as they are found exclusively in invertebrates. The insecticide amitraz is an agonist of octopamine receptors (40) and is widely used as pesticide in agriculture to control ticks (41,42)and parasitic mites (43). Its potential to control mosquito populations has been scarcely explored up to date (44–46). To investigate if the octopaminergic signalling in the mosquito ear is affected by amitraz, we tested its effects on the antennal fibrillae erection. Amitraz activates both α- and β-adrenergic-like receptor octopamine receptors in other insects (47,48), so we would expect it to induce the erection of the antennal fibrillae through its activation of the β-adrenergic-like octopamine receptor AGAP002886.

We exposed male mosquitoes to five min of 0.025%, 0.1% and 0.4% amitraz at ZT4 –when the antennal fibrillae are collapsed- and quantified the proportion of males with erected fibrillae in a time series -5, 10, 30, 60 minutes and 1 day after amitraz exposure (Fig. 8a). Five minutes after exposing the mosquitoes to any amitraz concentration, 98% of male mosquitoes erected their fibrillae. Moreover, thirty minutes after the exposure, more than 80% of mosquitoes still presented erected fibrillae ([0.025%] 80%; [0.1%] 100%; [0.4%] 90%;) and for the highest concentration (0.4%), 89% of mosquitoes still presented erected fibrillae after 1 hr (Fig. 8a). We also exposed *AGAP002886*^*-*^ homozygous mutant males to 0.1% amitraz to test if this receptor was necessary for the amitraz induction of fibrillae erection. As expected, we observed than none of the mutant mosquitoes erected the fibrillae upon amitraz exposure (Fig. 8b).

**Fig. 8:**
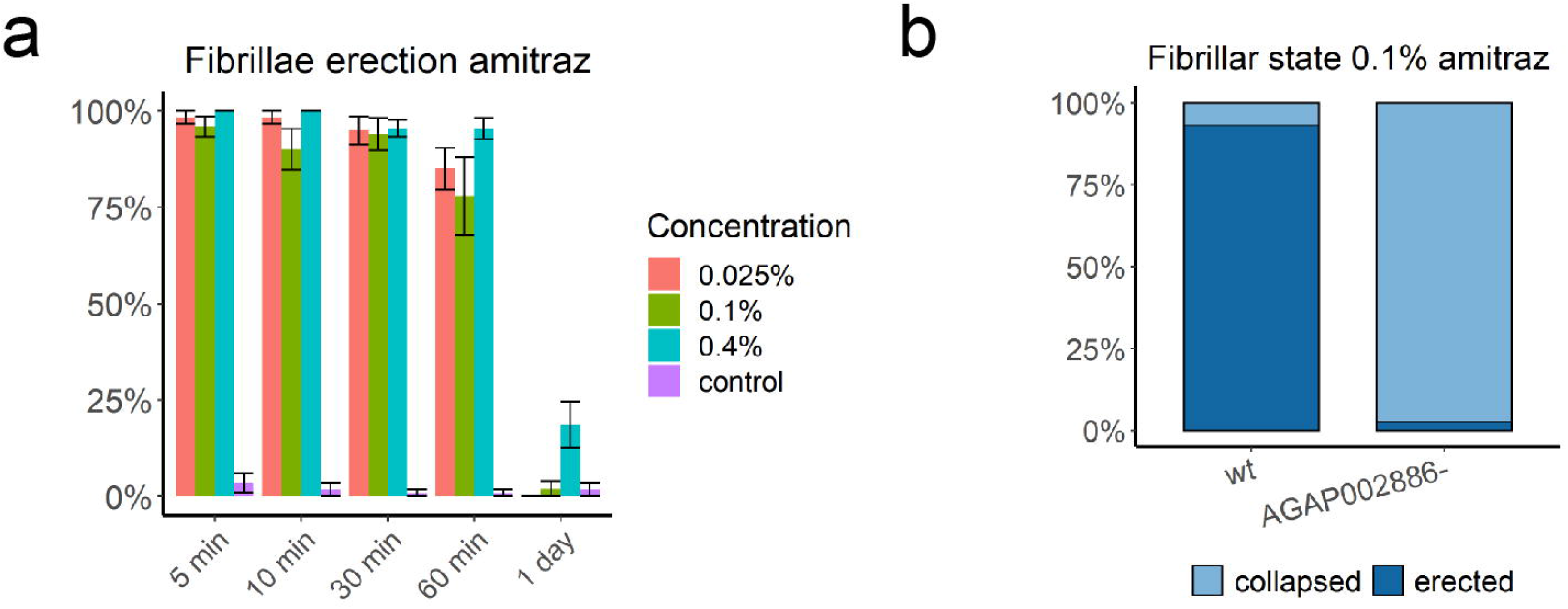
Amitraz targets the octopaminergic signalling in the malaria mosquito ear. **a)** Time series of the effects of different amitraz concentrations on the fibrillae erection of malaria mosquitoes. Amitraz exposure causes the erection of the antennal fibrillae that last 24 hours and even one day in almost 25% of mosquitoes for the highest amitraz concentration 0.4%. **b**) AGAP002886 knockouts exposed to amitraz do not erect their fibrillae.

## Discussion

Mosquito audition mediates the acoustic recognition of mating partners within crepuscular swarms. The highly transient nature of mosquito swarming behaviour suggests that its audition might be subject to plastic modulation to adapt it to the sensory requirements of the swarm. In this paper, we found evidence of an extensive auditory neuromodulatory network in malaria mosquitoes, with an impressive variety of neurotransmitter and neuropeptide receptors being expressed in the mosquito JO. Among those, we identified octopamine as potential auditory modulator during swarming as the expression of an α-adrenergic-like octopamine receptor (AGAP000045) peaks at this time. To confirm this, we examined the auditory role of octopamine in detail and demonstrated that it plays a sexually dimorphic role and modulates mostly male hearing. We showed that octopamine affects different auditory parameters that determine the male’s ability to detect the female flight tone including the auditory tuning and the flagellar stiffness. We also identified the receptors mediating octopamine auditory role: apart from AGAP000045, the novel β-adrenergic-like octopamine receptor AGAP002886 played a primary role. Knocking-out both mutants caused different auditory phenotypes and critically, AGAP002886 mutant mosquitoes fail to erect the antennal fibrillae. We also demonstrated that the octopaminergic pathways in the mosquito ear can be targeted by currently available insecticides, opening new paths for mosquito control.

The broad repertoire of neurotransmitters and neuropeptide receptors identified in our RNA-sequencing analysis suggests a high plasticity of mosquito hearing. This sensory plasticity likely enables the dynamic adaptation of auditory physiology to environemental (e.g. daytime-dependent) changes of auditory needs (male malaria mosquitoes are attracted to female tones only during swarm time); it could help the male mosquito ear to adjust to changes in the swarm’s sensory ecology (related e.g. to mosquito numbers, noise levels or temperature-dependent variations) and thus facilitate the detection and localisation of female mating partners. As the male mosquito ear harbours more than 16,000 sensory neurons, the neuromodulatory innervation should also contribute to establish neuronal subtypes that differ in their sensitivity to sound and their adaptation capabilities (49). Investigating the distribution and function of the neuromodulatory input seems essential to understand the outstanding performance of the mosquito ear as sound detector.

Our transcriptomic analysis showed that, as previously described in *Cx. quinquefasciatus* mosquitoes (6), the biogenic amines octopamine and serotonin also innervate the malaria mosquito ear, as we detected expression of several octopamine and serotonin receptors. In insects, the biogenic amines octopamine and serotonin modulate various aspects of the sensory organ physiology (17). Serotonin has been shown to modulate mating behaviour (50) and the entrainment of the circadian clock to visual stimuli (51) in *Drosophila*. Several GABA-A, GABA-B and nicotinic acetylcholine receptors were also highly expressed, suggesting a complex modulation of auditory physiology. Interestingly, acetylcholine is the main neurotransmitter of the vertebrate auditory efferent system. Auditory efferents innervating the cochlea facilitate the discrimination of signals from noise and contribute to sharpening of tuning curves (52). Both mechanisms would be relevant in the auditory environment of the swarm to detect the mating partner flight tones above the noise caused by hundreds of mosquitoes flying together. We also identified several neuropeptide receptors, including calcitonin receptors, which have been identified in the cochlea (53). Neuropeptide expression has been previously identified in the antenna of other insects (54) suggesting its implication in the modulation of the sensory function.

It has been known for decades that malaria mosquito males are attracted to female sounds (55,56) only during the circadian swarm time (10). Our initial hypothesis was that local changes in the auditory physiology should explain the temporal restriction of phonotactic responses to this circadian time. When we interrogated our male transcriptomic dataset for genes with cycling expression that peaked at swarm time, the single outcome was the octopamine receptor AGAP000045, suggesting the implication of octopamine in the modulation of hearing during swarming (Fig. 1). We then investigated whether the auditory function is at all subject to circadian control and what role octopamine plays in this potential sensory plasticity. The first indication of swarming-related changes in auditory physiology is provided by the erection of the antennal fibrillae. We have been able to demonstrate that octopamine controls this behaviour and that its effect is circadian time-dependent, as mosquitoes were more likely to erect fibrillae upon octopamine exposure during swarm time (Fig. 3a). We also detected circadian time-dependent changes in the mechanical state of the flagellum. During swarm time, male mosquito flagella were more likely to exhibit more extreme behaviours (either quiescent or SSO) rather than intermediate “transient” vibrations. Moreover, quiescent flagella were only observed at swarm time under control conditions (Fig. 3b). These results should be taken with caution as SSOs have been linked to the amplification and detection of female flight tones in the swarm (4,5,7). Our results do not contradict this view, they simply reveal a more dynamic flagellar mechanics during swarm time, which might switch between different oscillatory states (quiescent or SSO), without precluding a role of SSOs in female detection. We should also consider that manipulating the mosquito ear for our auditory tests might induce stress responses that can affect the flagellum, as SSOs have been previously shown to be induced by CO_2_ induced bouts of hypoxia (37). What our results clearly show is that injecting octopamine during the inactive phase at ZT4 causes similar flagellar effects as the circadian swarming time (reduction of transient and increase of quiescent flagella). At ZT12, this effect is more pronounced, and most flagella will stop SSOing and will remain quiescent. We cannot rule out that these effects are too severe compared to the natural role of octopamine as injecting octopamine might overflow the system. What we can certainly state is that octopamine action can influence the mechanical state of the flagellum and that sensitivity of the system to this effect is stronger during the circadian swarm time, likely reflecting the higher expression of octopamine receptors.

Acoustic mating partner detection in mosquitoes is based on the recognition of their wingbeat frequencies. Current theories on mosquito partner recognition support that the detection of the female mating partner by the male is based on a distortion product (DP) system, in which low frequency DPs generated in the male flagellum as the non-linear mixing of the male and female wingbeats (57) are detected by the male. It is also hypothesized that SSOs of the male flagellum act as amplifiers of the female wingbeats to increase the power of the DPs generated (7), and as such, facilitate the detection of the female. Moreover, recent research shows that males increase their wingbeat frequencies at swarm time thereby enhancing the audibility of females (57). Our data show that the SSO frequency increases at swarm time ZT12 and that octopamine exposure also causes a shift of the SSO frequency to higher frequencies (Fig. 3c). We therefore propose that endogenous octopamine is the modulator that drives the observed increase in SSO frequency at swarm time. This increase of the SSO frequency would be part of the male mechanisms that, together with the increase in the wingbeat frequency, will increase female audibility in the swarm. To clarify this, more experiments are required that involve stimulating the auditory nerve with two tones mimicking the male and female wingbeats and studying the effects on the nerve responses of changing the SSO frequency after octopamine exposure. These experiments will shed light on how mosquito partner detection takes place, and which is the role of octopamine in this system. We speculate that by modulating the SSO frequency, octopamine might contribute to adapting the male auditory system to optimize the female detection. Octopamine has also been shown to innervate flight muscles (58) and modulate flight performance (12). It is plausible that octopamine plays a multifunctional role in the male “swarming physiology” to enable the acoustic detection of the female mating partner in the swarm, by modulating the pattern of fibrillae erection, the auditory tuning and the frequency of the flight tones.

Octopamine exposure also causes an increase of the flagellar steady-state stiffness, which is sharper at ZT12 (Fig. 4e). This increase in the stiffness could subsequently cause effects on the other biophysical parameters studied, explaining some of the multimodal effects of octopamine action. But the flagellar ears of mosquitoes, especially those of males, are complex oscillators and the interrelation - and interdependencies - of their functional parameters are not well understood. Neuropharmacological manipulations, such as those presented here, will be a vital tool to dissect – and deconstruct – the underlying oscillatory complexity. Octopamine terminals have been observed at the base of the auditory cilia in *Cx. quinquefasciatus* mosquitoes (6). Steady-state flagellar stiffness is dependent on the combined elasticity of the components that suspend the sound receiver, but are not directly involved in the transmission of the sound stimulus (59,60). The location of octopamine terminals at the base of the auditory cilia could indicate that octopamine increases the tension of the cilia, stiffening the flagellum and modulating other auditory parameters such as auditory tuning, which would influence the capacity of the male mosquito to detect the female.

The role of the female hearing in mosquito partner detection remains elusive. Up to date, only males have been shown to be attracted to female sounds (10,55,56). There are very few examples of sound-induced behavioural responses reported in females (61). Our transcriptomic analysis showed nonetheless that most receptors identified in males were also expressed in females, supporting a fine plasticity of female hearing. Previous research on the anatomy of *An. gambiae* female JO shows that its efferent pattern is very reduced (7). This suggests that in females the ligands of the receptors identified in the RNA-Seq analysis might reach the ear though the hemolymph acting as neurohormones. The effect of octopamine in mosquito audition is also highly sexually dimorphic and females seems to respond very faintly to octopamine exposure. This agrees with previous studies that support a predominant role of the male in the acoustic detection of the mating partner in the swarm (9,57). The complexity of the female ear, both from an anatomical and transcriptomic perspective makes us hypothesize that we are missing something regarding the role of female hearing in the mosquito mating system. This is especially true for Culicid mosquitoes (e.g. *Aedes spp* or *Culex spp*), which have anatomically more complex ears than Anopheles (7).Developing finer behavioural assays might be necessary to understand if and how female audition contributes to mating partner detection.

In this study, we have also identified the receptors mediating octopamine auditory role. AGAP002286 is a novel β-adrenergic-like octopamine receptor orthologue of Octb2R in other insects, which are coupled to intracellular increases in cyclic AMP (62). Most octopamine-induced auditory responses studied, including the erection of the antennal fibrillae and the mechanical changes of the flagellum, were nearly abolished in *AGAP002886*^*-*^ mutants, supporting its role as the primary octopamine receptor in the mosquito ear (Fig. 5). The other octopamine receptor detected, the α-adrenergic-like octopamine receptor AGAP000045, showed a peak of expression at swarm time. *AGAP000045* displayed an intermediate phenotype, with octopamine-mediated auditory effects being reduced compared to wildtype animals, both regarding the fibrillae erection and the flagellar stiffness. Interestingly, none of the receptor mutants ceased SSOs upon octopamine exposure (Fig. 5b). Indeed, the biophysical parameters extracted from frequency-modulated sweep stimulation show stronger shifts in the values upon octopamine exposure in *AGAP000045* mutants than in wildtypes. In the mutants, octopamine signalling was not able to push the flagella from SSO to quiescent state (Fig.5b). However, in AGAP000045 mutants, SSO animals displayed greater shifts in the oscillator frequency and acceleration values than wildtype mosquitoes. This is possibly due to the fact that in wildtypes, the stronger effect of octopamine exposure would push flagella to quiescent states as the increase in the flagellar stiffness would reach too high values to maintain SSO. In AGAP000045 mutants, because the stiffness shift is weaker, mosquitoes can continue SSOs, but with greater shifts in the biophysical parameters. In AGAP002886 mutant mosquitoes, SSO values did barely change before and after octopamine injection. It seems therefore that AGAP002886 modulate more fundamental mechanisms than AGAP000045, which is involved in finer responses. Studying the anatomical distribution of these receptors might shed some light as to how the functional differences are established.

We finally tested whether the octopaminergic signalling in the mosquito ear could be targeted using pharmacological interventions to potentially disrupt mosquito partner recognition by disrupting mosquito heraing and thus eventually help control mosquito populations. We used the insecticide amitraz, an octopamine receptor agonist, as proof-of-concept of this approach. We found that amitraz activates AGAP002886 in the mosquito ear to induce the erection of the antennal fibrillae. This opens new pathways as to whether insecticides can be applied in the field to disrupt mosquito reproduction. Providing new molecular targets for insecticide development is indeed of highest priority to sustain malaria control and prevention in light of the emergence of resistance to currently available insecticides. The exquisitely complex mosquito auditory system might provide us with novel opportunities to do so.

## Methods

### Mosquito rearing and entrainment

*An. gambiae* Kisumu strain mosquitoes were used in the RNA-Seq analysis and were provided by Shahida Begum from the London School of Hygiene and Tropical Medicine. All other experiments used *An. gambiae* G3 strain mosquitoes, initially provided by Andrea Crisanti (Imperial College London) (63)until we started our own colony at the UCL Ear Institute insectary. Mosquitoes were reared in the insectary under a 12 h: 12 h light-dark cycle, 80% relative humidity and 28 °C. Mosquitoes were blood fed with horse blood (Fisher, Oxoid) using a Hemotek. All mosquitoes used for the experiments were 3 to 7 days old.

Experimental pupae were sex-separated and males and females were hatched in different cages to keep them virgin. Mosquitoes were then sorted into groups of five in glass vials and transferred to environmental incubators (Percival I-30 VL Multipurpose Plant Breeding Chamber, CLF PlantClimatics GmbH, Germany) for circadian entrainment. Entrainment conditions consisted of 12 h: 12 h light/dark cycle with a constant rate of light-intensity change from minimum to maximum from ZT12 to ZT13 and from maximum to minimum from ZT0 to ZT1 – where ZT stands for Zeitgeber Time, and ZTX is the formalized chronobiological notation of an entrained circadian cycle’s phase. Temperature and relative humidity were set constant throughout the circadian day, at 28 °C and 80% respectively. Mosquitoes were exposed to the circadian entrainment in incubators for three days before performing any experiment.

### RNA-Seq

#### Sample collection and preparation

On day four of circadian entrainment, 35 male and female mosquitoes were removed from the incubators at six different circadian time points (ZT0, ZT4, ZT8, ZT12, ZT16, ZT20), sedated with CO_2_ and transferred to Eppendorfs, which were in turn immediately transferred to liquid nitrogen to minimize sample degradation. The samples were kept in a storage freezer at -80 °C prior to dissections. During dissections, using a pair of forceps, flagella were first dissected off the JOs, which were subsequently removed from the mosquito heads and placed in eppendorfs immersed in ice. The mosquito mouthparts were also removed and the remainder of the heads were too transferred to (separate) eppendorfs.

#### Library preparation

RNA was extracted (Quiagen) and its integrity was confirmed using Agilent’s 2200 Tapestation.

Samples were processed using the SMART-Seq v4 Ultra Low Input RNA Kit (Clontech Laboratories, Inc.). Briefly, cDNA libraries were generated using the SMART (Switching Mechanism at 5’ End of RNA Template) technology which produces full-length PCR amplified cDNA starting from small amounts (500pg) of total RNA. 9 cycles of PCR were used to generate cDNA. The amplified cDNA was checked for integrity and quantity on the Agilent Bioanalyser using the High Sensitivity DNA kit.

150pg of cDNA was then converted to sequencing library using the Nextera XT DNA protocol (Illumina, San Diego, US). This uses a transposon able to fragment and tag double-stranded cDNA (tagmentation), followed by a limited PCR reaction (12 cycles) which adds sample specific indexes to allow multiplex sequencing.

#### Sequencing

Libraries to be multiplexed in the same run were pooled in equimolar quantities, calculated from Qubit and Tapestation fragment analysis.

Samples were sequenced on the NextSeq 500 instrument (Illumina, San Diego, US) using a 38 paired end run, generating 400M read pairs in total.

#### Data analysis

Fig. 1b shows a schematic of the data analysis workflow. FastQC (64)and MultiQC (65) were used to perform quality control of sequencing reads. These were subsequently classified, and read counts were estimated, with Kallisto (66), based on the the *Anopheles gambiae* transcriptome (https://vectorbase.org/vectorbase/app/downloads/release-53/AgambiaePEST/fasta/data/VectorBase-53_AgambiaePEST_AnnotatedTranscripts.fasta). Following a preliminary principal component analysis of sample counts, a male JO sample collected at ZT12 was deemed unsuitable for use in further analysis and discarded.

##### 1. Identifying transcript expression counts above the quantification noise floor

The remaining samples were used in classifying what we hereby call the male and female JO *expressed* genes. The idea behind this analysis was to identify the noise distribution of estimated transcript counts within the context of our RNA-sequencing experiment: A quantification of falsely and randomly assigned read counts. Contrasted to this distribution, then, we could make statistical claims of whether transcripts’ estimated expressions a) exceed the count noise floor and can thus be regarded as biologically expressed, or b) do not exceed the count noise floor and thus be regarded as noise. The formulation of the noise distribution was conducted as follows:

First, 50 transcripts were randomly selected, noted, and removed from the *Anopheles gambiae* transcriptome. Then, the remaining transcripts were used to simulate paired end reads with lengths and coverage similar to our sequenced reads. Simulation of reads was conducted with NEAT (67). The 50 transcripts that had been initially removed were subsequently re-introduced into the dataset to recover the whole transcriptome, and read counts for the simulated reads were estimated using Kallisto. Count estimates for the originally removed transcripts were extracted. Note that any counts obtained from these genes can be regarded as noise/false assignements, as they weren’t part of the dataset used to simulate reads to start with. The mean expression counts for these 50 transcripts – the mean noise of counts – was calculated. The whole process was repeated ten times, each time removing a different set of randomly selected transcripts, to obtain ten means for noise counts. These means were normally distributed (Shapiro-Wilk test), and their mean was calculated, assumed to represent the average of the underlying population of count noise, and used to define a Poisson distribution for the counts’ noise. This Poisson distribution served as the reference distribution against which transcript expression counts were compared.

Of the six timepoints in total, at which samples were collected (three biological replicates per timepoint) for each sex, the one with the highest mean expression counts for a specific gene was selected for comparison against the noise distribution. If expression counts for that gene were high enough such that the probability of them arising solely by noise, given the noise distribution, was < 0.05, then that gene was regarded as expressed. Probability values for these expression counts were examined after adjustment for multiple comparisons (Benjamini-Hochberg method).

##### 2. Classifying transcripts by gene ontology (GO)

Transcripts identified as expressed were further investigated in search for those that were assigned with any of the following GO terms:

| GO term | Definition |
| --- | --- |
| GO:0004952 | dopamine neurotransmitter receptor activity |
| GO:0004969 | histamine receptor activity |
| GO:0008226 | tyramine receptor activity |
| GO:0004930 | G protein-coupled receptor activity |
| GO:0008227 | G protein-coupled amine receptor activity |
| GO:0001653 | peptide receptor activity |
| GO:0015464 | acetylcholine receptor activity |
| GO:0016917 | GABA receptor activity |
| GO:0008066 | glutamate receptor activity |
| GO:0008066 | glutamate receptor activity |
| GO:0004984 | olfactory receptor activity |
| GO:0099589 | serotonin receptor activity |
| GO:0008527 | olfactory receptor activity |
| GO:0009881 | photoreceptor activity |

If any genes were assigned with a GO term ‘child’ (=subterm) of any of the above, but not with the ‘parental terms’ themselves, then we assigned it to the parental GO ourselves. Perusal through the hierarchy of GO terms for GO inheritance was conducted with the Python package goatools (68).

GO annotations of each of the transcripts of *Anopheles gambiae* were compiled from two sources:

1. The *Anopheles gambiae* gene association file (gaf) available for download on VectorBase (https://vectorbase.org/vectorbase/app/downloads/release-53/AgambiaePEST/gaf/VectorBase-53_AgambiaePEST_GO.gaf)
2. GO annotations of *Anopheles gambiae* transcript orthologs. This approach was only applied to instances where the transcript had not itself been annotated, but had a close ortholog that were assigned with any of the GO terms of interest. We focused on other mosquito species and drosophilids for finding orthologous genes.

##### 3. Differential expression analysis between male and female JOs

Normalization of read counts for comparability across, and comparison between male and female samples was conducted in R, with the package DESeq2 (69). The normalization method used was the default (median of ratios) method of DESeq2.

The circadian time of sampling introduces variation in transcript abundances, but the focus of this analysis was the source of variation due to the sex factor (male vs female). To adjust the read counts accordingly, and increase the sensitivity of finding transcriptional differences specific to the sex factor, the time factor was included in the design formula (i.e. ∼ time + sex).

The significance condition applied to comparisons was a False Discovery Rate (FDR) of < 0.05.

##### 4. Cycling expression analysis of male and female JOs

The cycling expression analysis was conducted individually for male and female samples. Normalization of read counts for comparability across the circadian time-points and a first round of statistical filtering of potential cycling candidates was conducted in R, with DESeq2. For the analysis of the diel pattern of gene expression (the aforementioned statistical filtering step), briefly, a “full” model of the counts was constructed including a term for time, and compared to a “reduced” model (lacking the time term) with the likelihood ratio test function of DESeq2 (conceptually similar to ANOVA) – null hypothesis being that a model of the data incorporating time offers no improvement over one which doesn’t. Significance condition applied was FDR < 0.05.

Transcripts that passed the first round of filtering were further investigated for cycling via the R-package MetaCycle (70), using the JTK_CYCLE function for rhythmicity detection. Once again, significance condition applied was FDR < 0.05.

### CRISPR-Cas 9 mediated mutant generation

#### Generation of gRNA constructs

We designed gRNAs targeting coding regions of AGAP000045 and AGAP002886 using CRISPOR (http://crispor.tefor.net/). We selected gRNAs with no SNPs reported in *An. gambiae* G3 strain (AGAP000045 gRNA sequence: GTGGACGGATCCGACCAATC; AGAP002886 gRNA sequence: GGTCGTTCATGTGTGACGTG). gRNAs were cloned by Golden Gate cloning (NEB) into a BSaI digested p165 vector (kindly donated by Roberto Galizi). The plasmid p165 contains the vasa2 promoter driving the expression of a human codon-optimized version of the *Streptococcus pyogenes* Cas9 gene (hCas9) and vasa 3’ UTR regulatory sequence upstream a U6::gRNA cassette containing a spacer cloning sites, all flanked by attB recombination sites (71). The full sequence of vector p165 has been deposited to GenBank (accession ID: KU189142).

#### Generation of donor constructs

Homology-directed repair was used to disrupt the coding sequences of the targeted genes. Homology arms were cloned by Gibson cloning into a donor vector (kindly provided by Roberto Galizi) that had been designed to contain an *attP*-flanked 3xP3::GFP marker construct enclosed within the homology arms, as well as an external *3xP3::RFP* marker. Homology arms extended 2kb in either direction of the expected CRISPR-Cas9 cleavage site and were amplified from genomic DNA using the primers that include overlapping ends for Gibson cloning (<u>underlined</u>):

AGAP00045: AGAP000045_5’_attP_F (<u>CTCGAGTTTTTCAGCAAGAT</u>GTGTACCGCTCGAATCCAAC); AGAP000045_5’_attP_F (<u>CCAGTTGGGG</u>CCACTGTACGGACGCGAG); AGAP000045_5’_3’_F (<u>CGTACAGTGG</u>CCCCAACTGGGGTAACCTTT); AGAP000045_5’_3’_R (<u>GGCGAGCACC</u>CCCCAACTGGGGTAACCTTT); AGAP000045_3’_attP_F (<u>CCAGTTGGGG</u>GGTGCTCGCCTTCATCAAC) and AGAP000045_3’_attP_R (<u>AGGAGATCTTCTAGAAAGAT</u>TTACTCCTCCAGACCCCGTA) AGAP002886: AGAP002886_5’_attP_F (<u>CTCGAGTTTTTCAGCAAGAT</u>GTCGTGTGCCTGGCCTC); AGAP002886_5’_attP_F (<u>CCAGTTGGGG</u>ACATCCAGACGTCGACCG); AGAP002886_5’_3’_F (<u>GTCTGGATGT</u>CCCCAACTGGGGTAACCTTT); AGAP002886_5’_3’_R (<u>GAGGCCGTGC</u>CCCCAACTGGGGTAACCTTT); AGAP002886_3’_attP_F (<u>CCAGTTGGGG</u>GCACGGCCTCGATCCTG) and AGAP002886_3’_attP_R (<u>AGGAGATCTTCTAGAAAGAT</u>GAGAGAGCGAGAGAGCAAGA)

##### Microinjection of mosquito embryos and selection of transformants

Freshly laid *Anopheles gambiae* G3 embryos were aligned and used for microinjections as described (66). The donor construct (200 ng/μl) containing regions of homology to the relevant target locus was injected together with the relevant CRISPR plasmid (200 ng/μl). All surviving G0 larvae were crossed to wild-type mosquitoes and F1 positive transformants were identified using a fluorescence microscope (Eclipse TE200) as eGFP+ larvae. To confirm that candidate transformants were knock-outs of the gene of interest, a PCR was performed with primers binding at both sites of the homology arms in the genomic DNA region.

### Tests of auditory function

#### Laser Doppler vibrometry (LDV)

Glass vials containing five mosquitoes were extracted from the environmental incubators at the required circadian time points. Mosquitoes were glued to a Teflon rod using blue-light-cured dental glue (as reported previously for both mosquitoes (7)and *Drosophila* (60)). The mosquito body was immobilised via glue application to minimise disturbances caused by mosquito movements, but leaving thoracic spiracles free for the mosquito to breathe. The left flagellum was glued to the head and further glue was applied between the pedicels, with only the right flagellum remaining free to move.

Following this gluing procedure (7), the rod holding the mosquito was placed in a micromanipulator on a vibration isolation table, with the mosquito facing the laser Doppler vibrometer at a 90° angle. To minimise mechanical disturbances, the laser was focused at the second flagellomere from the flagellum tip in males and the third flagellomere from the tip in females. All recordings were done using a PSV-400 laser Doppler vibrometer (Polytec) with an OFV-70 close up unit and a DD-500 displacement decoder. Fig. 2 shows a sketch of the laser Doppler vibrometry (LDV) experimental paradigm. All measurements were taken in a temperature-controlled room (22°C) at the different circadian times specified in each experiment.

#### Compound injection procedure

One millimolar and 10mM octopamine hydrochloride (Sigma) solutions in Ringer (72)were prepared fresh on the day of the injection experiment. Sharpened micro-capillaries were filled with the either octopamine or control ringer solutions. The tip of the micro-capillary was inserted into the thorax of a mounted mosquito and the solution was injected to flood the entire inset body and reach the JO. In all injection experiments, the protocol included creating recordings at three distinct stages: 1) baseline prior to any injections; 2) following injection of a control ringer solution; 3) following injection of either 1mM octopamine, 10mM octopamine or a 2^nd^ ringer control injection. This protocol allowed us to collect baseline and compound injection data in the same experiment for comparative purposes, as well as ensuring that the mosquito was healthy prior to any injections.

#### Fibrillae erection assessment

Theantennal fibrillar state (collapsed vs erected) was assessed at the baseline or after the compound injection.

### LDV Data Analysis

All data have undergone a curation process to ensure that the laser was appropriately positioned on the mosquito flagellum during an experimental run. This is done through a diagnostic channel which measures and records the laser backscatter of the LDV. All experimental runs are segregated into two large categories: free-fluctuation runs -where the flagellum is allowed to operate in its natural state unstimulated, and stimulus runs – where an electrostatic waveform is imposed on the flagellum to act as a driving force. The experimental structure interleaves unstimulated and stimulated periods, with unstimulated and stimulated parts continually alternating. Each ‘run’ contains one unstimulated and one stimulated dataset with a duration of 1 second each. The pipeline for each of those experimental runs as well as a novel formalism of the auditory mechanical state is detailed as below.

#### Unstimulated free-fluctuation analysis

The unstimulated data form the basis of the LDV analysis as the frequency, amplitude and mechanical state can be determined prior to the stimulated counterpart. The frequency and amplitude of the oscillator is extracted though a Fast Fourier transform (FFT), whereas the mechanical state is determined by looking at the amplitude distribution of the flagellar motion. The amplitude distribution of a fully quiescent flagellum follows a standard normal distribution, whereas an SSOing animal would instead follow an arcsine (73). An added complication comes from the appearance of transient mechanical states which tend to lie somewhere in between a QUIES and SSO state. To account for those states, we generate a mixed distribution comprised of an arcsine and normal distribution, controlled by a weight parameter α which ranges between 0 (denoting purely QUIES) and 1 (denoting purely SSO).

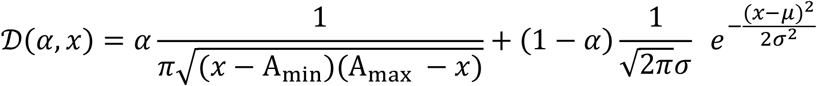

Where α denotes the weight parameter, *A*_*min*_ and *A*_*max*_ the amplitude extremes of the input signal, 𝜎 the width of the normal distribution and 𝜇 the center of the distribution. Due to the extreme variability of transient states in terms of their distribution profile, they are excluded from the majority of the LDV analysis. To permit some level of flexibility on the mechanical state, the cut-off condition for QUIES and SSO animals is set between [0 - 0.1] and [0.9 - 1.0] respectively and require a goodness of fit to account for 99.7% of the data.

We further formalise the SSO definition by adding another criterion where an animals is only considered as SSO if it sustained its oscillatory behaviour for three consecutive experimental runs, corresponding to a total span of six seconds. This condition was set to avoid attributing the SSO label to a briefly oscillating animal.

#### Force-step stimulation recordings and analysis

The force-step stimulation protocol was used to calibrate the maximum flagellar displacement to approximately ±8,000 nm and to extract some principal parameters of flagellar mechanics. Here, the flagellum was stimulated using electrostatic force-step stimuli. LDV measurements of flagellar displacements and electrophysiological activity were recorded simultaneously. For analysis, mosquito apparent flagellar mass estimates were calculated as in (7). Force-step stimulation analysis was performed as in (74).

#### Frequency-modulated sweep stimulation and analysis

The stimulated half of each experimental run is performed by electrostatically driving the flagellum (insert citation from past work). The waveform applied is a linear frequency sweep which ranges from 0 Hz to 1 kHz and its mirrored version of 1 kHz to 0 Hz (Fig.9a). Each experimental run is always followed by its mirrored version (Fig. 2d) to determine any latency due to hysteresis.

The profile of the stimulated data follows a well-known line shape that is defined by the driven damped harmonic oscillator. The so-called ‘envelope’ of the waveform is extracted using a statistic-sensitive nonlinear iterative peak clipping (SNIP) method of background estimation. The envelope is then fitted through the theoretical expectation from the driven harmonic oscillator.

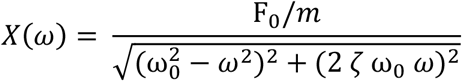

Where *F*_0_ the driving force, m the effective mass of the flagellum, *ω*_0_ the natural frequency of the oscillator, 𝜁 the damping ratio and *ω* the driving frequency. To reduce the degrees of freedom of the system for fitting purposes, the force and effective mass terms are absorbed into a singular parameter which effectively represents the acceleration experienced by the system.

These fitted parameters convey important biophysical information which fully characterises the system, however they may not necessarily be intuitive. A commonly used derived parameter that describes a - damped oscillator is the Q-factor, or quality factor, defined as *Q* = 1/2𝜁, which can be used to qualitatively interpret the level of dampening experienced by the flagellum. It is important to note that when the system is undergoing SSO, the system does not experience any effective damping (as it actively compensates for it) indicating that the above equations do not apply, however in the limit where 𝜁 → 0, the Q-factor tends to infinity (a perfect, undamped oscillator).

Another useful derived parameter is that of the optimum frequency of the driven damped oscillator 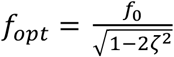which describes the frequency at which the oscillatory system achieves its maximum amplitude as a function of the driving frequency. It is simple enough to see that in the limit case where *ζ* → 0, *f*_*opt*_ = *f*_0_ where the maximum amplitude of the oscillator occurs at exactly the natural frequency of the oscillator. Any deviation of the damping ratio from 0 would lead to an apparent shift of the observed maximum amplitude at frequencies higher than the natural frequency of the oscillator.

##### Amitraz exposure experiments

Glass vials were coated with 0.2 ml of 0.025%, 0.1% or 0.4% amitraz (Sigma, 45323) in acetone or acetone alone as control. Vials were allowed to dry for two hours in a dark and cool room. Mosquitoes were collected from experimental cages at ZT4 and placed in paper cups. Single mosquitoes were carefully aspirated into the glass vials and exposed for 5 min to amitraz. Upon exposure, mosquitoes were transferred back to the paper cups and provided with 10% sucrose solution. Parafilm was used to cover the cup and keep the moisture for 24 hrs. Fibriallae erection was assessed 5 min, 10 min, 30 min, 60 min and 1 day after exposure.

##### Statistical analysis

Samples sizes for LDV experiments were determined based on published data on Dipteran antennal LDV measurements (7,73,75). Within-group variation estimates were calculated as part of standard statistical tests and were reasonable for the type of recordings.

Statistical test for normality (Shapiro-Wilk test with a significance level of *p < 0*.*05*) were used for each LDV dataset. These were generally found to be non-normally distributed; thus, median and standard error values are reported throughout.

For the free fluctuations data, non-parametric ANOVA on rank test was used to compare the flagellar BF obtained from free fluctuations fits across the different circadian time points in female and male mosquitoes. Hochberg correction was used to control for multiple comparisons.

For the fibrillae erection experiments and amitraz exposure experiments, Chi-squared test was used to compare the frequencies of fibrillae erection and mechanical states across the different compound injections and circadian time points categories.

For the force-step and frequency-modulated sweep stimulation, ANOVA on ranks or Wilcoxon signed-rank test was used to compare the stiffness and peak mechanical frequency between different compound injection and circadian time points in female and male mosquitoes.

All statistical tests were done in R 3.6.0.

## Acknowledgements

**Shahida Begum, Imperial, France Kisumu eggs. Infravec. Roberto Galizi for support on mutant generation design and Andrea Crisanti for initial provision with mosquitoes.**

This work received funding from UK Research and Innovation under the Future Leaders Fellowship scheme (Grant reference MR/S015493/1 to M.A.), from European Union’s Horizon 2020 research and innovation programme (Fellowship 752472 to M.A.), from European Research Council (ERC) under the Horizon 2020 research and innovation programme (Consolidator Grant agreement No 648709, to J.T.A.), two UCL Global Challenges Research Fund (GCRF) small grants (to J.T.A. and M.A., respectively), a Wellcome ISSF grant (204841/Z/16/Z) to M.A., a grant from the Biotechnology and Biological Sciences Research Council, UK (BBSRC, BB/V007866/1 to J.T.A.) and a grant from The Human Frontier Science Program (HFSP grant RGP0033/2021 to J.T.A.). This publication was also supported by the project Research Infrastructures for the control of vector-borne diseases (Infravec2), which has received funding from the European Union’s Horizon 2020 research and innovation program under grant agreement No 731060. Embryo microinjections and molecular characterisation of transformants were performed by Crisanti Lab at Imperial College London, for which we thank Prof Andrea Crisanti, Dr Roya Haghighat-Khah, Dr Carla Siniscalchi, Dr Rocco D’Amato, Louise Marston and Bathsheba L Gardner.

## Figure legends

**Table 1: Potential neuromodulatory genes expressed in the male and female *An. gambiae* mosquito JO**. Expression of neurotransmitter and neuropeptide receptors, as well as receptors associated with other sensory modalities. Genes have been arranged according to their molecular function based on GO terms assigned to them. Complete gene names can be found in Supp. Table 1.

## REFERENCES

1. Clements AN. The biology of mosquitoes. Sensory reception and behaviour. 1999. Wallingford: CABI Publishing Google Scholar.

2. Diabaté A, Yaro AS, Dao A, Diallo M, Huestis DL, Lehmann T. Spatial distribution and male mating success of Anopheles gambiaeswarms. BMC Evolutionary Biology. 2011;11:184.

3. Cator LJ, Arthur BJ, Harrington LC, Hoy RR. Harmonic convergence in the love songs of the dengue vector mosquito. Science. 2009 Feb 20;323(5917):1077–9.

4. Pennetier C, Warren B, Dabir KR, Russell IJ, Gibson G. ‘Singing on the wing’ as a mechanism for species recognition in the malarial mosquito Anopheles gambiae. Curr Biol. 2010 Jan;20(2):131–6.

5. Warren B, Gibson G, Russell IJ. Sex Recognition through midflight mating duets in Culex mosquitoes is mediated by acoustic distortion. Curr Biol. 2009 Mar;19(6):485–91.

6. Andrés M, Seifert M, Spalthoff C, Warren B, Weiss L, Giraldo D, et al. Auditory Efferent System Modulates Mosquito Hearing. Current Biology. 2016 Aug 8;26(15):2028–36.

7. Su MP, Andrés M, Boyd-Gibbins N, Somers J, Albert JT. Sex and species specific hearing mechanisms in mosquito flagellar ears. Nature Communications. 2018 Sep 25;9(1):3911.

8. Warren B, Russell I. Mosquitoes on the wing “tune in” to acoustic distortion. In AIP; 2011. p. 479–80.

9. Simões PMV, Ingham RA, Gibson G, Russell IJ. A role for acoustic distortion in novel rapid frequency modulation behaviour in free-flying male mosquitoes. The Journal of Experimental Biology. 2016;219(13):2039–47.

10. Charlwood JD, Jones MDR. Mating in the mosquito, Anopheles gambiae sl. II Swarming behaviour. Physiological Entomology. 1980;5(4):315–20.

11. Nijhout HF. Control of antennal hair erection in male mosquitoes. The Biological Bulletin. 1977 Dec 1;153(3):591–603.

12. Roeder T. Tyramine and octopamine: ruling behavior and metabolism. Annu Rev Entomol. 2005;50:447–77.

13. Farooqui T. Octopamine-Mediated Neuromodulation of Insect Senses. Neurochem Res. 2007 Sep 1;32(9):1511–29.

14. Linn CE, Campbell MG, Poole KR, Wu WQ, Roelofs WL. Effects of photoperiod on the circadian timing of pheromone response in male Trichoplusia ni: Relationship to the modulatory action of octopamine. Journal of Insect Physiology. 1996 Sep 1;42(9):881–91.

15. Schendzielorz T, Schirmer K, Stolte P, Stengl M. Octopamine Regulates Antennal Sensory Neurons via Daytime-Dependent Changes in cAMP and IP3 Levels in the Hawkmoth Manduca sexta. PLOS ONE. 2015 Mar 18;10(3):e0121230.

16. Flecke C, Stengl M. Octopamine and tyramine modulate pheromone-sensitive olfactory sensilla of the hawkmoth Manduca sexta in a time-dependent manner. Journal of Comparative Physiology A. 2009 Jun 1;195(6):529–45.

17. Zhukovskaya MI, Polyanovsky AD. Biogenic Amines in Insect Antennae. Front Syst Neurosci [Internet]. 2017 [cited 2017 Jul 6];11. Available from: http://journal.frontiersin.org/article/10.3389/fnsys.2017.00045/full

18. Ma Z, Guo X, Lei H, Li T, Hao S, Kang L. Octopamine and tyramine respectively regulate attractive and repulsive behavior in locust phase changes. Sci Rep. 2015;5:8036.

19. Andrés M, Su MP, Albert J, Cator LJ. Buzzkill: targeting the mosquito auditory system. Current Opinion in Insect Science. 2020 Aug 1;40:11–7.

20. Hill CA, Sharan S, Watts VJ. Genomics, GPCRs and new targets for the control of insect pests and vectors. Current Opinion in Insect Science. 2018 Dec 1;30:99–106.

21. Pietrantonio PV, Xiong C, Nachman RJ, Shen Y. G protein-coupled receptors in arthropod vectors: omics and pharmacological approaches to elucidate ligand-receptor interactions and novel organismal functions. Current opinion in insect science. 2018;

22. Audsley N, Down RE. G protein coupled receptors as targets for next generation pesticides. Insect Biochemistry and Molecular Biology. 2015 Dec 1;67(Supplement C):27–37.

23. Du W, Awolola TS, Howell P, Koekemoer LL, Brooke BD, Benedict MQ, et al. Independent mutations in the Rdl locus confer dieldrin resistance to Anopheles gambiae and An. arabiensis. Insect Molecular Biology. 2005;14(2):179–83.

24. Lustig LR. Nicotinic acetylcholine receptor structure and function in the efferent auditory system. The Anatomical Record Part A: Discoveries in Molecular, Cellular, and Evolutionary Biology. 2006;288A(4):424–34.

25. Knecht ZA, Silbering AF, Ni L, Klein M, Budelli G, Bell R, et al. Distinct combinations of variant ionotropic glutamate receptors mediate thermosensation and hygrosensation in Drosophila. Elife. 2016 Sep 22;5.

26. Nässel DR. Neuropeptides in the nervous system of Drosophila and other insects: multiple roles as neuromodulators and neurohormones. Prog Neurobiol. 2002 Sep;68(1):1–84.

27. Ignell R, Root CM, Birse RT, Wang JW, Nässel DR, Winther ÅME. Presynaptic peptidergic modulation of olfactory receptor neurons in Drosophila. PNAS. 2009 Aug 4;106(31):13070–5.

28. Hou L, Yang P, Jiang F, Liu Q, Wang X, Kang L. The neuropeptide F/nitric oxide pathway is essential for shaping locomotor plasticity underlying locust phase transition. eLife [Internet]. [cited 2021 Aug 4];6. Available from: https://www.ncbi.nlm.nih.gov/pmc/articles/PMC5400507/

29. Leung NY, Montell C. Unconventional Roles of Opsins. Annual Review of Cell and Developmental. 2017;33(1):241–64.

30. Zanini D, Giraldo D, Warren B, Katana R, Andres M, Reddy S, et al. Proprioceptive Opsin Functions in Drosophila Larval Locomotion. Neuron. 2018 Apr 4;98(1):67-74.e4.

31. Senthilan PR, Piepenbrock D, Ovezmyradov G, Nadrowski B, Bechstedt S, Pauls S, et al. Drosophila auditory organ genes and genetic hearing defects. Cell. 2012 Aug;150(5):1042–54.

32. Kastner KW, Shoue DA, Estiu GL, Wolford J, Fuerst MF, Markley LD, et al. Characterization of the Anopheles gambiae octopamine receptor and discovery of potential agonists and antagonists using a combined computational-experimental approach. Malaria Journal. 2014 Nov 18;13:434.

33. Eilerts DF, VanderGiessen M, Bose EA, Broxton K, Vinauger C. Odor-Specific Daily Rhythms in the Olfactory Sensitivity and Behavior of Aedes aegypti Mosquitoes. Insects. 2018 Oct 23;9(4):E147.

34. Rund SSC, Bonar NA, Champion MM, Ghazi JP, Houk CM, Leming MT, et al. Daily rhythms in antennal protein and olfactory sensitivity in the malaria mosquito Anopheles gambiae. Sci Rep. 2013;3:2494.

35. Charlwood J, D. R. Jones M. Mating behaviour in the mosquito, Anopheles gambiae s.l. I. Close range and contact behaviour. Vol. 4. 1979. 111–120 p.

36. Gibson G, Warren B, Russell IJ. Humming in tune: sex and species recognition by mosquitoes on the wing. J Assoc Res Otolaryngol. 2010 Dec;11(4):527–40.

37. Gpfert MC, Robert D. Active auditory mechanics in mosquitoes. Proc Biol Sci. 2001 Feb;268(1465):333–9.

38. Wu SF, Yao Y, Huang J, Ye GY. Characterization of a ?-adrenergic-like octopamine receptor from the rice stem borer (Chilo suppressalis). J Exp Biol. 2012 Aug;215(Pt 15):2646–52.

39. Huang J. Molecular Pharmacology and Physiology of Insect Biogenic Amine Receptors. In: Advances in Agrochemicals: Ion Channels and G Protein-Coupled Receptors (GPCRs) as Targets for Pest Control [Internet]. American Chemical Society; 2017 [cited 2019 Sep 10]. p. 127–38. (ACS Symposium Series; vol. 1265). Available from: https://doi.org/10.1021/bk-2017-1265.ch007

40. Evans PD, Gee JD. Action of formamidine pesticides on octopamine receptors. Nature. 1980 Sep 4;287(5777):60–2.

41. Haigh AJB, Gichang MM. The activity of amitraz against infestations of Rhipicephalus appendiculatus. Pesticide Science. 1980;11(6):674–8.

42. Vudriko P, Okwee-Acai J, Byaruhanga J, Tayebwa DS, Okech SG, Tweyongyere R, et al. Chemical tick control practices in southwestern and northwestern Uganda. Ticks and Tick-borne Diseases. 2018 May 1;9(4):945–55.

43. Pugliese N, Circella E, Cocciolo G, Giangaspero A, Tomic DH, Kika TS, et al. Efficacy of λ-cyhalothrin, amitraz, and phoxim against the poultry red mite Dermanyssus gallinae De Geer, 1778 (Mesostigmata: Dermanyssidae): an eight-year survey. Avian Pathology. 2019 Sep 13;48(sup1):S35–43.

44. Ahmed MAI, Vogel CFA. The role of octopamine receptor agonists in the synergistic toxicity of certain insect growth regulators (IGRs) in controlling Dengue vector Aedes aegypti (Diptera: Culicidae) mosquito. Acta Tropica. 2016 Mar 1;155(Supplement C):1–5.

45. Ahmed MAI, Vogel CFA. Synergistic action of octopamine receptor agonists on the activity of selected novel insecticides for control of dengue vector Aedes aegypti (Diptera: Culicidae) mosquito. Pesticide Biochemistry and Physiology. 2015 May 1;120(Supplement C):1–6.

46. Fuchs S, Rende E, Crisanti A, Nolan T. Disruption of aminergic signalling reveals novel compounds with distinct inhibitory effects on mosquito reproduction, locomotor function and survival. Scientific Reports. 2014 Jul 2;4:srep05526.

47. Guo L, Fan X yu, Qiao X, Montell C, Huang J. An octopamine receptor confers selective toxicity of amitraz on honeybees and Varroa mites. Sen S, VijayRaghavan K, Sen S, Yamanaka N, editors. eLife. 2021 Jul 12;10:e68268.

48. Kita T, Hayashi T, Ohtani T, Takao H, Takasu H, Liu G, et al. Amitraz and its metabolite differentially activate α-and β-adrenergic-like octopamine receptors. Pest Management Science. 2017;73(5):984– 90.

49. Lapshin DN, Vorontsov DD. Frequency organization of the Johnston’s organ in male mosquitoes (Diptera, Culicidae). Journal of Experimental Biology. 2017 Nov 1;220(21):3927–38.

50. Pooryasin A, Fiala A. Identified Serotonin-Releasing Neurons Induce Behavioral Quiescence and Suppress Mating in Drosophila. J Neurosci. 2015 Sep 16;35(37):12792–812.

51. Yuan Q, Lin F, Zheng X, Sehgal A. Serotonin modulates circadian entrainment in Drosophila. Neuron. 2005 Jul;47(1):115–27.

52. Ryugo DK, Fay RR, Popper AN. Auditory and vestibular efferents. Vol. 38. Springer Science & Business Media; 2010.

53. Kitajiri M, Yamashita T, Tohyama Y, Kumazawa T, Takeda N, Kawasaki Y, et al. Localization of calcitonin gene-related peptide in the organ of Corti of the rat: an immunohistochemical study. Brain research. 1985;358(1–2):394–7.

54. Latorre-Estivalis JM, Sterkel M, Ons S, Lorenzo MG. Transcriptomics supports local sensory regulation in the antenna of the kissing-bug Rhodnius prolixus. BMC Genomics. 2020 Jan 30;21(1):101.

55. Roth LM. A study of mosquito behavior. An experimental laboratory study of the sexual behavior of Aedes aegypti (Linnaeus). The American Midland Naturalist. 1948;40(2):265–352.

56. Belton P. Attractton of male mosquttoes to sound. J Am Mosq Control. 1994;10:297–301.

57. Somers J, Georgiades M, Su MP, Bagi J, Andrés M, Alampounti A, et al. Hitting the right note at the right time: Circadian control of audibility in Anopheles mosquito mating swarms is mediated by flight tones. Sci Adv. 2022 Jan 14;8(2):eabl4844.

58. Pauls D, Blechschmidt C, Frantzmann F, Jundi B el, Selcho M. A comprehensive anatomical map of the peripheral octopaminergic/tyraminergic system of Drosophila melanogaster. Scientific Reports. 2018 Oct 17;8(1):1–12.

59. Albert JT, Nadrowski B, Göpfert MC. Drosophila mechanotransduction--linking proteins and functions. Fly (Austin). 2007 Aug;1(4):238–41.

60. Effertz T, Wiek R, Goepfert MC. NompC TRP Channel Is Essential for Drosophila Sound Receptor Function. Current Biology. 2011 Apr;21(7):592–7.

61. Bartlett-Healy K, Crans W, Gaugler R. Phonotaxis to Amphibian Vocalizations in Culex territans (Diptera: Culicidae). Ann Entomol Soc Am. 2008 Jan 1;101(1):95–103.

62. Maqueira B, Chatwin H, Evans PD. Identification and characterization of a novel family of Drosophilaβ-adrenergic-like octopamine G-protein coupled receptors. Journal of Neurochemistry. 2005 Jul 1;94(2):547–60.

63. Haghighat-Khah RE, Sharma A, Wunderlich MR, Morselli G, Marston LA, Bamikole C, et al. Cellular mechanisms regulating synthetic sex ratio distortion in the Anopheles gambiae germline. Pathogens and Global Health. 2020 Oct 2;114(7):370–8.

64. Andrews S. (2010). FastQC: a quality control tool for high throughput sequence data. Available online at: http://www.bioinformatics.babraham.ac.uk/projects/fastqc.

65. Ewels P, Magnusson M, Lundin S, Käller M. MultiQC: summarize analysis results for multiple tools and samples in a single report. Bioinformatics. 2016 Oct 1;32(19):3047–8.

66. Nl B, H P, P M, L P. Near-optimal probabilistic RNA-seq quantification [Internet]. Nature biotechnology. 2016 [cited 2020 Oct 16]. Available from: https://pubmed.ncbi.nlm.nih.gov/27043002/

67. NEAT: an efficient network enrichment analysis test | BMC Bioinformatics | Full Text [Internet]. [cited 2022 Aug 2]. Available from: https://bmcbioinformatics.biomedcentral.com/articles/10.1186/s12859-016-1203-6

68. Klopfenstein DV, Zhang L, Pedersen BS, Ramírez F, Warwick Vesztrocy A, Naldi A, et al. GOATOOLS: A Python library for Gene Ontology analyses. Sci Rep. 2018 Jul 18;8(1):10872.

69. Mi L, W H, S A. Moderated estimation of fold change and dispersion for RNA-seq data with DESeq2 [Internet]. Genome biology. 2014 [cited 2020 Oct 16]. Available from: https://pubmed.ncbi.nlm.nih.gov/25516281/

70. G W, Rc A, Me H, K K, Jb H. MetaCycle: an integrated R package to evaluate periodicity in large scale data [Internet]. Bioinformatics (Oxford, England). 2016 [cited 2020 Oct 16]. Available from: https://pubmed.ncbi.nlm.nih.gov/27378304/

71. Hammond A, Galizi R, Kyrou K, Simoni A, Siniscalchi C, Katsanos D, et al. A CRISPR-Cas9 Gene Drive System Targeting Female Reproduction in the Malaria Mosquito vector Anopheles gambiae. Nat Biotechnol. 2016 Jan;34(1):78–83.

72. Baines RA, Bate M. Electrophysiological Development of Central Neurons in theDrosophila Embryo. J Neurosci. 1998 Jun 15;18(12):4673–83.

73. Mc G, Ad H, Jt A, D R, O H. Power gain exhibited by motile mechanosensory neurons in Drosophila ears [Internet]. Proceedings of the National Academy of Sciences of the United States of America. 2005 [cited 2020 Oct 19]. Available from: https://pubmed.ncbi.nlm.nih.gov/15623551/

74. Mechanical Signatures of Transducer Gating in the Drosophila Ear: Current Biology [Internet]. [cited 2020 Oct 16]. Available from: https://www.cell.com/current-biology/fulltext/S0960-9822(07)01381-4?code=cell-site

75. Effertz T, Nadrowski B, Piepenbrock D, Albert JT, Gopfert MC. Direct gating and mechanical integrity of Drosophila auditory transducers require TRPN1. Nat Neurosci. 2012 Sep;15(9):1198–200.

