## Supplemental Table 12 and 13 for "A novel beta-adrenergic like octopamine receptor modulates the audition of malaria mosquitoes and serves as insecticide target"

**Supplementary table 12: Summary of steady-state stiffness values extracted from force-step stimulation responses in wildtype male mosquitoes upon exposure to octopamine**

|  | condition | parameters | mean | sd | median | se |
| --- | --- | --- | --- | --- | --- | --- |
| 1 | ZT4_G3_Ringer2 | Steady-state stiffness | 222.3794 | 129.4798 | 181.992 | 40.94512 |
| 2 | ZT4_G3_OA1 | Steady-state stiffness | 456.8191 | 364.7843 | 332.2196 | 105.3041 |
| 3 | ZT4_G3_OA10 | Steady-state stiffness | 531.0592 | 473.7489 | 329.3449 | 149.8126 |
| 4 | ZT12_G3_Ringer2 | Steady-state stiffness | 337.7086 | 335.975 | 194.562 | 106.2446 |
| 5 | ZT12_G3_OA1 | Steady-state stiffness | 488.5785 | 366.961 | 395.9188 | 129.7403 |
| 6 | ZT12_G3_OA10 | Steady-state stiffness | 1442.868 | 868.1435 | 1257.664 | 250.6114 |

**Supplementary table 13: Wilcoxon signed-rank test on force-step stimulation analysis in males (pairwise comparisons).**

|  | .y. | group1 | group2 | p | p.adj | p.format | p.signif | method |
| --- | --- | --- | --- | --- | --- | --- | --- | --- |
| 1 | value | ZT04\|ringer2 | ZT12\|ringer2 | 0.393048 | 1 | 0.39305 | ns | Wilcoxon |
| 2 | value | ZT04\|ringer2 | ZT04\|OA1mM | 0.024916 | 0.25 | 0.02492 | * | Wilcoxon |
| 3 | value | ZT04\|ringer2 | ZT12\|OA1mM | 0.043421 | 0.39 | 0.04342 | * | Wilcoxon |
| 4 | value | ZT04\|ringer2 | ZT04\|OA10mM | 0.052426 | 0.42 | 0.05243 | ns | Wilcoxon |
| 5 | value | ZT04\|ringer2 | ZT12\|OA10mM | 6.19E-06 | 9.30E-05 | 6.20E-06 | **** | Wilcoxon |
| 6 | value | ZT12\|ringer2 | ZT04\|OA1mM | 0.122911 | 0.86 | 0.12291 | ns | Wilcoxon |
| 7 | value | ZT12\|ringer2 | ZT12\|OA1mM | 0.14571 | 0.87 | 0.14571 | ns | Wilcoxon |
| 8 | value | ZT12\|ringer2 | ZT04\|OA10mM | 0.247451 | 1 | 0.24745 | ns | Wilcoxon |
| 9 | value | ZT12\|ringer2 | ZT12\|OA10mM | 0.000473 | 0.0066 | 0.00047 | *** | Wilcoxon |
| 10 | value | ZT04\|OA1mM | ZT12\|OA1mM | 0.792077 | 1 | 0.79208 | ns | Wilcoxon |
| 11 | value | ZT04\|OA1mM | ZT04\|OA10mM | 0.821204 | 1 | 0.8212 | ns | Wilcoxon |
| 12 | value | ZT04\|OA1mM | ZT12\|OA10mM | 0.000496 | 0.0066 | 0.0005 | *** | Wilcoxon |
| 13 | value | ZT12\|OA1mM | ZT04\|OA10mM | 0.828557 | 1 | 0.82856 | ns | Wilcoxon |
| 14 | value | ZT12\|OA1mM | ZT12\|OA10mM | 0.002159 | 0.026 | 0.00216 | ** | Wilcoxon |
| 15 | value | ZT04\|OA10mM | ZT12\|OA10mM | 0.005645 | 0.062 | 0.00564 | ** | Wilcoxon |
