## Supplemental Table 14 and 15 for "A novel beta-adrenergic like octopamine receptor modulates the audition of malaria mosquitoes and serves as insecticide target"

**Supplementary table 14: Summary of biophysical parameters extracted from frequency-modulated sweep responses in wildtype female mosquitoes upon exposure to octopamine**

|  | condition | parameters | mean | sd | median | se |
| --- | --- | --- | --- | --- | --- | --- |
| 1 | ZT4_G3_Ringer2 | F0m | 28.09141 | 10.57646 | 22.14493 | 0.454298 |
| 2 | ZT4_G3_Ringer2 | F0 | 331.3676 | 25.92163 | 330.5037 | 1.113429 |
| 3 | ZT4_G3_Ringer2 | damp_ratio | 0.426086 | 0.093004 | 0.428096 | 0.003995 |
| 4 | ZT4_G3_Ringer2 | Q-factor | -7.46E-08 | 1.61E-07 | -7.42E-08 | 6.91E-09 |
| 5 | ZT4_G3_OA1 | F0m | 26.89788 | 8.827836 | 25.52511 | 0.425716 |
| 6 | ZT4_G3_OA1 | F0 | 325.3265 | 36.00657 | 311.4013 | 1.736391 |
| 7 | ZT4_G3_OA1 | damp_ratio | 0.389959 | 0.111398 | 0.39421 | 0.005372 |
| 8 | ZT4_G3_OA1 | Q-factor | -1.19E-07 | 1.03E-07 | -1.45E-07 | 4.98E-09 |
| 9 | ZT12_G3_Ringer2 | F0m | 29.44617 | 12.26527 | 26.81448 | 0.681404 |
| 10 | ZT12_G3_Ringer2 | F0 | 388.9516 | 85.66718 | 343.4757 | 4.759288 |
| 11 | ZT12_G3_Ringer2 | damp_ratio | 0.404838 | 0.029336 | 0.399706 | 0.00163 |
| 12 | ZT12_G3_Ringer2 | Q-factor | -6.37E-08 | 1.10E-07 | -9.11E-08 | 6.12E-09 |
| 13 | ZT12_G3_OA1 | F0m | 40.33739 | 21.3275 | 31.08491 | 1.030904 |
| 14 | ZT12_G3_OA1 | F0 | 441.7661 | 138.5823 | 351.7447 | 6.698629 |
| 15 | ZT12_G3_OA1 | damp_ratio | 0.324653 | 0.27572 | 0.409373 | 0.013327 |
| 16 | ZT12_G3_OA1 | Q-factor | -1.66E-07 | 3.22E-07 | -8.23E-08 | 1.56E-08 |

**Supplementary table 15: Wilcoxon signed-rank test on frequency-modulated sweep analysis in females (pairwise comparisons).**

|  | .y. | group1 | group2 | p | p.adj | p.format | p.signif | method |
| --- | --- | --- | --- | --- | --- | --- | --- | --- |
| 1 | F0m | ZT4\|Ringer 2 | ZT4\|1mM OA | 0.981922 | 0.98 | 0.98 | ns | Wilcoxon |
| 2 | F0m | ZT4\|Ringer 2 | ZT12\|Ringer 2 | 0.428129 | 0.86 | 0.43 | ns | Wilcoxon |
| 3 | F0m | ZT4\|Ringer 2 | ZT12\|1mM OA | 3.36E-27 | 2.00E-26 | <2e-16 | **** | Wilcoxon |
| 4 | F0m | ZT4\|1mM OA | ZT12\|Ringer 2 | 0.185775 | 0.56 | 0.19 | ns | Wilcoxon |
| 5 | F0m | ZT4\|1mM OA | ZT12\|1mM OA | 3.16E-24 | 1.60E-23 | <2e-16 | **** | Wilcoxon |
| 6 | F0m | ZT12\|Ringer 2 | ZT12\|1mM OA | 9.27E-18 | 3.70E-17 | <2e-16 | **** | Wilcoxon |
| 7 | f0 | ZT4\|Ringer 2 | ZT4\|1mM OA | 3.15E-10 | 6.30E-10 | 3.20E-10 | **** | Wilcoxon |
| 8 | f0 | ZT4\|Ringer 2 | ZT12\|Ringer 2 | 1.10E-42 | 6.60E-42 | < 2e-16 | **** | Wilcoxon |
| 9 | f0 | ZT4\|Ringer 2 | ZT12\|1mM OA | 7.83E-17 | 2.30E-16 | < 2e-16 | **** | Wilcoxon |
| 10 | f0 | ZT4\|1mM OA | ZT12\|Ringer 2 | 2.02E-42 | 1.00E-41 | < 2e-16 | **** | Wilcoxon |
| 11 | f0 | ZT4\|1mM OA | ZT12\|1mM OA | 1.75E-40 | 7.00E-40 | < 2e-16 | **** | Wilcoxon |
| 12 | f0 | ZT12\|Ringer 2 | ZT12\|1mM OA | 0.169085 | 0.17 | 0.17 | ns | Wilcoxon |
| 13 | damp_ratio | ZT4\|Ringer 2 | ZT4\|1mM OA | 8.25E-27 | 4.90E-26 | < 2e-16 | **** | Wilcoxon |
| 14 | damp_ratio | ZT4\|Ringer 2 | ZT12\|Ringer 2 | 1.20E-24 | 6.00E-24 | < 2e-16 | **** | Wilcoxon |
| 15 | damp_ratio | ZT4\|Ringer 2 | ZT12\|1mM OA | 0.000643 | 0.0026 | 0.00064 | *** | Wilcoxon |
| 16 | damp_ratio | ZT4\|1mM OA | ZT12\|Ringer 2 | 0.400995 | 0.4 | 0.401 | ns | Wilcoxon |
| 17 | damp_ratio | ZT4\|1mM OA | ZT12\|1mM OA | 0.00138 | 0.0041 | 0.00138 | ** | Wilcoxon |
| 18 | damp_ratio | ZT12\|Ringer 2 | ZT12\|1mM OA | 0.002154 | 0.0043 | 0.00215 | ** | Wilcoxon |
| 19 | Q | ZT4\|Ringer 2 | ZT4\|1mM OA | 8.25E-27 | 4.90E-26 | < 2e-16 | **** | Wilcoxon |
| 20 | Q | ZT4\|Ringer 2 | ZT12\|Ringer 2 | 1.20E-24 | 6.00E-24 | < 2e-16 | **** | Wilcoxon |
| 21 | Q | ZT4\|Ringer 2 | ZT12\|1mM OA | 0.000643 | 0.0026 | 0.00064 | *** | Wilcoxon |
| 22 | Q | ZT4\|1mM OA | ZT12\|Ringer 2 | 0.400995 | 0.4 | 0.401 | ns | Wilcoxon |
| 23 | Q | ZT4\|1mM OA | ZT12\|1mM OA | 0.00138 | 0.0041 | 0.00138 | ** | Wilcoxon |
| 24 | Q | ZT12\|Ringer 2 | ZT12\|1mM OA | 0.002154 | 0.0043 | 0.00215 | ** | Wilcoxon |

**Supplementary table 12: Summary of steady-state stiffness values extracted from force-step stimulation responses in wildtype female mosquitoes upon exposure to octopamine**

|  | condition | parameters | mean | sd | median | se |
| --- | --- | --- | --- | --- | --- | --- |
| 1 | ZT4_G3_Ringer2 | Steady-state stiffness | 91.70988 | 15.74046 | 92.88549 | 5.565091 |
| 2 | ZT4_G3_OA1 | Steady-state stiffness | 117.3927 | 13.22724 | 116.227 | 4.676537 |
| 3 | ZT12_G3_Ringer2 | Steady-state stiffness | 144.9563 | 25.04838 | 144.3444 | 12.52419 |
| 4 | ZT12_G3_OA1 | Steady-state stiffness | 140.7325 | 83.51008 | 116.654 | 29.52527 |

**Supplementary table 13: Wilcoxon signed-rank test on force-step stimulation analysis in females (pairwise comparisons).**

|  | .y. | group1 | group2 | p | p.adj | p.format | p.signif | method |
| --- | --- | --- | --- | --- | --- | --- | --- | --- |
| 1 | value | ZT4\|ringer2 | ZT4\|OA1mM | 0.006993 | 0.035 | 0.007 | ** | Wilcoxon |
| 2 | value | ZT4\|ringer2 | ZT12\|ringer2 | 0.00404 | 0.024 | 0.004 | ** | Wilcoxon |
| 3 | value | ZT4\|ringer2 | ZT12\|OA1mM | 0.037918 | 0.15 | 0.038 | * | Wilcoxon |
| 4 | value | ZT4\|OA1mM | ZT12\|ringer2 | 0.153535 | 0.46 | 0.154 | ns | Wilcoxon |
| 5 | value | ZT4\|OA1mM | ZT12\|OA1mM | 0.878477 | 0.88 | 0.878 | ns | Wilcoxon |
| 6 | value | ZT12\|ringer2 | ZT12\|OA1mM | 0.214141 | 0.46 | 0.214 | ns | Wilcoxon |
