## Supplemental Table 18 and 19 for "A novel beta-adrenergic like octopamine receptor modulates the audition of malaria mosquitoes and serves as insecticide target"

**Supplementary table 18: Summary of biophysical parameters extracted from frequency-modulated sweep responses in octopamine receptor mutant male mosquitoes upon exposure to octopamine**

|  | condition | State | parameters | mean | sd | median | se |
| --- | --- | --- | --- | --- | --- | --- | --- |
| 1 | ZT12_G3_baseline | SSO | F0m | 23.51785 | 6.32611 | 22.22667 | 0.373418 |
| 2 | ZT12_G3_baseline | SSO | F0 | 335.0872 | 132.7153 | 344.4433 | 7.833938 |
| 3 | ZT12_G3_baseline | SSO | damp_ratio | 0.122903 | 0.155253 | 0.151577 | 0.009164 |
| 4 | ZT12_G3_baseline | SSO | Q-factor | 2.11E-07 | 2.18E-07 | 2.18E-07 | 1.29E-08 |
| 5 | ZT12_G3_baseline | Quiesc | F0m | 33.28834 | 1.694667 | 33.55677 | 0.691845 |
| 6 | ZT12_G3_baseline | Quiesc | F0 | 380.9039 | 3.764656 | 380.8853 | 1.536914 |
| 7 | ZT12_G3_baseline | Quiesc | damp_ratio | 0.331114 | 0.008712 | 0.334231 | 0.003557 |
| 8 | ZT12_G3_baseline | Quiesc | Q-factor | -1.12E-07 | 3.75E-08 | -1.17E-07 | 1.53E-08 |
| 9 | ZT12_G3_OA1 | SSO | F0m | -11.2426 | 356.0905 | 20.44885 | 32.92058 |
| 10 | ZT12_G3_OA1 | SSO | F0 | 402.8885 | 139.1633 | 382.3274 | 12.86565 |
| 11 | ZT12_G3_OA1 | SSO | damp_ratio | 0.038395 | 0.230228 | 0.157269 | 0.021285 |
| 12 | ZT12_G3_OA1 | SSO | Q-factor | 7.09E-07 | 2.88E-06 | 4.68E-07 | 2.67E-07 |
| 13 | ZT12_G3_OA1 | Quiesc | F0m | -16.2579 | 354.9364 | 14.3172 | 19.74919 |
| 14 | ZT12_G3_OA1 | Quiesc | F0 | 548.7785 | 679.0663 | 544.5224 | 37.78426 |
| 15 | ZT12_G3_OA1 | Quiesc | damp_ratio | 0.254112 | 2.080702 | 0.218421 | 0.115773 |
| 16 | ZT12_G3_OA1 | Quiesc | Q-factor | 2.66E-07 | 5.46E-07 | 7.84E-08 | 3.04E-08 |
| 17 | ZT12_45_baseline | SSO | F0m | 25.55014 | 6.400079 | 22.4257 | 0.529674 |
| 18 | ZT12_45_baseline | SSO | F0 | 358.5642 | 84.87775 | 369.0642 | 7.024532 |
| 19 | ZT12_45_baseline | SSO | damp_ratio | -0.01237 | 0.176524 | -0.14721 | 0.014609 |
| 20 | ZT12_45_baseline | SSO | Q-factor | 6.51E-07 | 3.32E-07 | 7.24E-07 | 2.75E-08 |
| 21 | ZT12_45_baseline | Quiesc | F0m | 37.63418 | 0.523657 | 37.60586 | 0.058184 |
| 22 | ZT12_45_baseline | Quiesc | F0 | 419.6491 | 15.22066 | 424.1329 | 1.691184 |
| 23 | ZT12_45_baseline | Quiesc | damp_ratio | -0.05701 | 0.190704 | -0.19441 | 0.021189 |
| 24 | ZT12_45_baseline | Quiesc | Q-factor | -1.28E-07 | 4.36E-08 | -1.19E-07 | 4.84E-09 |
| 25 | ZT12_45_OA1 | SSO | F0m | 37.63535 | 484.1196 | 28.14991 | 28.93168 |
| 26 | ZT12_45_OA1 | SSO | F0 | 500.6864 | 315.8505 | 449.1201 | 18.87568 |
| 27 | ZT12_45_OA1 | SSO | damp_ratio | -0.00467 | 1.546803 | -0.15212 | 0.092439 |
| 28 | ZT12_45_OA1 | SSO | Q-factor | 6.65E-07 | 1.32E-06 | 5.08E-07 | 7.90E-08 |
| 29 | ZT12_45_OA1 | Quiesc | F0m | 198.9999 | 2016.157 | 1.544019 | 311.0997 |
| 30 | ZT12_45_OA1 | Quiesc | F0 | 691.3077 | 577.3947 | 402.9542 | 89.09394 |
| 31 | ZT12_45_OA1 | Quiesc | damp_ratio | 0.257348 | 0.546095 | 0.40434 | 0.084264 |
| 32 | ZT12_45_OA1 | Quiesc | Q-factor | -8.22E-07 | 1.38E-05 | 7.56E-07 | 2.13E-06 |
| 33 | ZT12_86_baseline | SSO | F0m | 27.84938 | 6.784535 | 25.84024 | 0.333845 |
| 34 | ZT12_86_baseline | SSO | F0 | 393.6869 | 21.31009 | 397.5879 | 1.048601 |
| 35 | ZT12_86_baseline | SSO | damp_ratio | 0.025358 | 0.240736 | -0.11342 | 0.011846 |
| 36 | ZT12_86_baseline | SSO | Q-factor | 5.02E-07 | 3.36E-07 | 4.46E-07 | 1.66E-08 |
| 37 | ZT12_86_OA1 | SSO | F0m | 29.66392 | 5.40886 | 30.01088 | 0.292906 |
| 38 | ZT12_86_OA1 | SSO | F0 | 331.4586 | 189.0484 | 382.8744 | 10.23755 |
| 39 | ZT12_86_OA1 | SSO | damp_ratio | 0.072516 | 0.257088 | -0.11133 | 0.013922 |
| 40 | ZT12_86_OA1 | SSO | Q-factor | 6.31E-07 | 3.31E-07 | 6.18E-07 | 1.79E-08 |

**Supplementary table 19: Wilcoxon signed-rank test on frequency-modulated sweep analysis in octopamine receptor mutant males (pairwise comparisons).**

|  | .y. | group1 | group2 | p | p.adj | p.format | p.signif | method |
| --- | --- | --- | --- | --- | --- | --- | --- | --- |
| 1 | F0m | Baseline\|wt | Baseline\|AGAP000045- | 1.34E-05 | 4.00E-05 | 1.30E-05 | **** | Wilcoxon |
| 2 | F0m | Baseline\|wt | Baseline\|AGAP002886- | 2.26E-16 | 2.90E-15 | 2.30E-16 | **** | Wilcoxon |
| 3 | F0m | Baseline\|wt | 1mM OA\|wt | 0.387475 | 0.77 | 0.39 | ns | Wilcoxon |
| 4 | F0m | Baseline\|wt | 1mM OA\|AGAP000045- | 7.52E-15 | 6.80E-14 | 7.50E-15 | **** | Wilcoxon |
| 5 | F0m | Baseline\|wt | 1mM OA\|AGAP002886- | 3.02E-27 | 4.20E-26 | < 2e-16 | **** | Wilcoxon |
| 6 | F0m | Baseline\|AGAP000045- | Baseline\|AGAP002886- | 2.12E-08 | 8.50E-08 | 2.10E-08 | **** | Wilcoxon |
| 7 | F0m | Baseline\|AGAP000045- | 1mM OA\|wt | 2.40E-09 | 1.40E-08 | 2.40E-09 | **** | Wilcoxon |
| 8 | F0m | Baseline\|AGAP000045- | 1mM OA\|AGAP000045- | 6.76E-09 | 3.40E-08 | 6.80E-09 | **** | Wilcoxon |
| 9 | F0m | Baseline\|AGAP000045- | 1mM OA\|AGAP002886- | 7.58E-14 | 6.10E-13 | 7.60E-14 | **** | Wilcoxon |
| 10 | F0m | Baseline\|AGAP002886- | 1mM OA\|wt | 2.80E-16 | 3.40E-15 | 2.80E-16 | **** | Wilcoxon |
| 11 | F0m | Baseline\|AGAP002886- | 1mM OA\|AGAP000045- | 6.20E-16 | 6.80E-15 | 6.20E-16 | **** | Wilcoxon |
| 12 | F0m | Baseline\|AGAP002886- | 1mM OA\|AGAP002886- | 3.18E-10 | 2.20E-09 | 3.20E-10 | **** | Wilcoxon |
| 13 | F0m | 1mM OA\|wt | 1mM OA\|AGAP000045- | 6.32E-16 | 6.80E-15 | 6.30E-16 | **** | Wilcoxon |
| 14 | F0m | 1mM OA\|wt | 1mM OA\|AGAP002886- | 1.36E-33 | 2.00E-32 | < 2e-16 | **** | Wilcoxon |
| 15 | F0m | 1mM OA\|AGAP000045- | 1mM OA\|AGAP002886- | 0.582645 | 0.77 | 0.58 | ns | Wilcoxon |
| 16 | f0 | Baseline\|wt | Baseline\|AGAP000045- | 2.30E-12 | 1.20E-11 | 2.30E-12 | **** | Wilcoxon |
| 17 | f0 | Baseline\|wt | Baseline\|AGAP002886- | 3.18E-41 | 3.80E-40 | < 2e-16 | **** | Wilcoxon |
| 18 | f0 | Baseline\|wt | 1mM OA\|wt | 1.26E-20 | 1.00E-19 | < 2e-16 | **** | Wilcoxon |
| 19 | f0 | Baseline\|wt | 1mM OA\|AGAP000045- | 2.81E-62 | 3.90E-61 | < 2e-16 | **** | Wilcoxon |
| 20 | f0 | Baseline\|wt | 1mM OA\|AGAP002886- | 3.78E-19 | 2.60E-18 | < 2e-16 | **** | Wilcoxon |
| 21 | f0 | Baseline\|AGAP000045- | Baseline\|AGAP002886- | 1.16E-32 | 1.20E-31 | < 2e-16 | **** | Wilcoxon |
| 22 | f0 | Baseline\|AGAP000045- | 1mM OA\|wt | 1.85E-06 | 5.60E-06 | 1.90E-06 | **** | Wilcoxon |
| 23 | f0 | Baseline\|AGAP000045- | 1mM OA\|AGAP000045- | 1.42E-58 | 1.90E-57 | < 2e-16 | **** | Wilcoxon |
| 24 | f0 | Baseline\|AGAP000045- | 1mM OA\|AGAP002886- | 4.34E-10 | 1.70E-09 | 4.30E-10 | **** | Wilcoxon |
| 25 | f0 | Baseline\|AGAP002886- | 1mM OA\|wt | 0.054732 | 0.11 | 0.055 | ns | Wilcoxon |
| 26 | f0 | Baseline\|AGAP002886- | 1mM OA\|AGAP000045- | 1.27E-40 | 1.40E-39 | < 2e-16 | **** | Wilcoxon |
| 27 | f0 | Baseline\|AGAP002886- | 1mM OA\|AGAP002886- | 1.52E-13 | 9.10E-13 | 1.50E-13 | **** | Wilcoxon |
| 28 | f0 | 1mM OA\|wt | 1mM OA\|AGAP000045- | 1.23E-21 | 1.10E-20 | < 2e-16 | **** | Wilcoxon |
| 29 | f0 | 1mM OA\|wt | 1mM OA\|AGAP002886- | 0.138722 | 0.14 | 0.139 | ns | Wilcoxon |
| 30 | f0 | 1mM OA\|AGAP000045- | 1mM OA\|AGAP002886- | 2.87E-63 | 4.30E-62 | < 2e-16 | **** | Wilcoxon |
| 31 | damp_ratio | Baseline\|wt | Baseline\|AGAP000045- | 0.110713 | 0.44 | 0.1107 | ns | Wilcoxon |
| 32 | damp_ratio | Baseline\|wt | Baseline\|AGAP002886- | 5.71E-07 | 6.90E-06 | 5.70E-07 | **** | Wilcoxon |
| 33 | damp_ratio | Baseline\|wt | 1mM OA\|wt | 1.07E-05 | 0.00011 | 1.10E-05 | **** | Wilcoxon |
| 34 | damp_ratio | Baseline\|wt | 1mM OA\|AGAP000045- | 0.038339 | 0.19 | 0.0383 | * | Wilcoxon |
| 35 | damp_ratio | Baseline\|wt | 1mM OA\|AGAP002886- | 6.91E-09 | 9.00E-08 | 6.90E-09 | **** | Wilcoxon |
| 36 | damp_ratio | Baseline\|AGAP000045- | Baseline\|AGAP002886- | 8.28E-07 | 9.10E-06 | 8.30E-07 | **** | Wilcoxon |
| 37 | damp_ratio | Baseline\|AGAP000045- | 1mM OA\|wt | 6.41E-05 | 0.00045 | 6.40E-05 | **** | Wilcoxon |
| 38 | damp_ratio | Baseline\|AGAP000045- | 1mM OA\|AGAP000045- | 0.417669 | 1 | 0.4177 | ns | Wilcoxon |
| 39 | damp_ratio | Baseline\|AGAP000045- | 1mM OA\|AGAP002886- | 0.001264 | 0.0076 | 0.0013 | ** | Wilcoxon |
| 40 | damp_ratio | Baseline\|AGAP002886- | 1mM OA\|wt | 3.53E-11 | 4.90E-10 | 3.50E-11 | **** | Wilcoxon |
| 41 | damp_ratio | Baseline\|AGAP002886- | 1mM OA\|AGAP000045- | 0.874183 | 1 | 0.8742 | ns | Wilcoxon |
| 42 | damp_ratio | Baseline\|AGAP002886- | 1mM OA\|AGAP002886- | 1.48E-11 | 2.20E-10 | 1.50E-11 | **** | Wilcoxon |
| 43 | damp_ratio | 1mM OA\|wt | 1mM OA\|AGAP000045- | 1.08E-05 | 0.00011 | 1.10E-05 | **** | Wilcoxon |
| 44 | damp_ratio | 1mM OA\|wt | 1mM OA\|AGAP002886- | 0.509422 | 1 | 0.5094 | ns | Wilcoxon |
| 45 | damp_ratio | 1mM OA\|AGAP000045- | 1mM OA\|AGAP002886- | 1.37E-05 | 0.00011 | 1.40E-05 | **** | Wilcoxon |
| 46 | Q | Baseline\|wt | Baseline\|AGAP000045- | 2.04E-06 | 1.60E-05 | 2.00E-06 | **** | Wilcoxon |
| 47 | Q | Baseline\|wt | Baseline\|AGAP002886- | 2.26E-07 | 2.00E-06 | 2.30E-07 | **** | Wilcoxon |
| 48 | Q | Baseline\|wt | 1mM OA\|wt | 5.40E-10 | 5.90E-09 | 5.40E-10 | **** | Wilcoxon |
| 49 | Q | Baseline\|wt | 1mM OA\|AGAP000045- | 0.928491 | 1 | 0.928 | ns | Wilcoxon |
| 50 | Q | Baseline\|wt | 1mM OA\|AGAP002886- | 3.34E-08 | 3.30E-07 | 3.30E-08 | **** | Wilcoxon |
| 51 | Q | Baseline\|AGAP000045- | Baseline\|AGAP002886- | 1.43E-14 | 2.10E-13 | 1.40E-14 | **** | Wilcoxon |
| 52 | Q | Baseline\|AGAP000045- | 1mM OA\|wt | 1.81E-10 | 2.20E-09 | 1.80E-10 | **** | Wilcoxon |
| 53 | Q | Baseline\|AGAP000045- | 1mM OA\|AGAP000045- | 0.590661 | 1 | 0.591 | ns | Wilcoxon |
| 54 | Q | Baseline\|AGAP000045- | 1mM OA\|AGAP002886- | 0.327217 | 0.98 | 0.327 | ns | Wilcoxon |
| 55 | Q | Baseline\|AGAP002886- | 1mM OA\|wt | 1.99E-11 | 2.60E-10 | 2.00E-11 | **** | Wilcoxon |
| 56 | Q | Baseline\|AGAP002886- | 1mM OA\|AGAP000045- | 0.014155 | 0.085 | 0.014 | * | Wilcoxon |
| 57 | Q | Baseline\|AGAP002886- | 1mM OA\|AGAP002886- | 1.48E-11 | 2.10E-10 | 1.50E-11 | **** | Wilcoxon |
| 58 | Q | 1mM OA\|wt | 1mM OA\|AGAP000045- | 5.80E-05 | 0.00041 | 5.80E-05 | **** | Wilcoxon |
| 59 | Q | 1mM OA\|wt | 1mM OA\|AGAP002886- | 0.026994 | 0.13 | 0.027 | * | Wilcoxon |
| 60 | Q | 1mM OA\|AGAP000045- | 1mM OA\|AGAP002886- | 0.042202 | 0.17 | 0.042 | * | Wilcoxon |
