## Supplemental Table 20 and 21 for "A novel beta-adrenergic like octopamine receptor modulates the audition of malaria mosquitoes and serves as insecticide target"

**Supplementary table 20: Summary of steady-state stiffness values extracted from force-step stimulation responses in octopamine receptor mutant male mosquitoes upon exposure to octopamine**

|  | condition | parameters | mean | sd | median | se |
| --- | --- | --- | --- | --- | --- | --- |
| 1 | ZT12_G3_Ringer2 | Steady-state stiffness | 229.4069 | 140.9107 | 180.2018 | 49.81945 |
| 2 | ZT12_45_Ringer2 | Steady-state stiffness | 160.6101 | 24.17638 | 159.6484 | 8.54764 |
| 3 | ZT12_2886_Ringer2 | Steady-state stiffness | 156.1379 | 25.8627 | 150.6974 | 9.775183 |
| 4 | ZT12_G3_OA1 | Steady-state stiffness | 488.5785 | 366.961 | 395.9188 | 129.7403 |
| 5 | ZT12_45_OA1 | Steady-state stiffness | 302.5479 | 345.3858 | 177.5663 | 122.1123 |
| 6 | ZT12_2886_OA1 | Steady-state stiffness | 145.6313 | 40.17464 | 136.9603 | 15.18459 |

**Supplementary table 21: Wilcoxon signed-rank test on force-step stimulation analysis in octopamine receptor mutant males (pairwise comparisons).**

|  | .y. | group1 | group2 | p | p.adj | p.format | p.signif | method |
| --- | --- | --- | --- | --- | --- | --- | --- | --- |
| 1 | value | wt baseline | wt OA1mM | 0.035556 | 0.39 | 0.0356 | * | Wilcoxon |
| 2 | value | wt baseline | AGAP002886- baseline | 0.020513 | 0.25 | 0.0205 | * | Wilcoxon |
| 3 | value | wt baseline | AGAP002886- OA1mM | 0.054079 | 0.54 | 0.0541 | ns | Wilcoxon |
| 4 | value | wt baseline | AGAP000045- baseline | 0.104895 | 0.73 | 0.1049 | ns | Wilcoxon |
| 5 | value | wt baseline | AGAP000045- OA1mM | 0.95913 | 1 | 0.9591 | ns | Wilcoxon |
| 6 | value | wt OA1mM | AGAP002886- baseline | 0.001243 | 0.017 | 0.0012 | ** | Wilcoxon |
| 7 | value | wt OA1mM | AGAP002886- OA1mM | 0.002176 | 0.028 | 0.0022 | ** | Wilcoxon |
| 8 | value | wt OA1mM | AGAP000045- baseline | 0.001088 | 0.016 | 0.0011 | ** | Wilcoxon |
| 9 | value | wt OA1mM | AGAP000045- OA1mM | 0.064957 | 0.58 | 0.065 | ns | Wilcoxon |
| 10 | value | AGAP002886- baseline | AGAP002886- OA1mM | 0.382867 | 1 | 0.3829 | ns | Wilcoxon |
| 11 | value | AGAP002886- baseline | AGAP000045- baseline | 0.866511 | 1 | 0.8665 | ns | Wilcoxon |
| 12 | value | AGAP002886- baseline | AGAP000045- OA1mM | 0.120591 | 0.73 | 0.1206 | ns | Wilcoxon |
| 13 | value | AGAP002886- OA1mM | AGAP000045- baseline | 0.396892 | 1 | 0.3969 | ns | Wilcoxon |
| 14 | value | AGAP002886- OA1mM | AGAP000045- OA1mM | 0.072106 | 0.58 | 0.0721 | ns | Wilcoxon |
| 15 | value | AGAP000045- baseline | AGAP000045- OA1mM | 0.234499 | 1 | 0.2345 | ns | Wilcoxon |
