## Supplemental Table 8 and 9 for "A novel beta-adrenergic like octopamine receptor modulates the audition of malaria mosquitoes and serves as insecticide target"

| condition | State | parameter | mean | sd | median | se |
| --- | --- | --- | --- | --- | --- | --- |
| ZT4_G3_Ringer2 | SSO | Frequency (Hz) | 350.0115 | 10.84086 | 351.1111 | 0.551785 |
| ZT4_G3_Ringer2 | SSO | Amplitude (Hz) | 519.4915 | 298.8118 | 409.0495 | 15.20912 |
| ZT4_G3_OA1 | SSO | Frequency (Hz) | 491.5491 | 70.91821 | 517.7778 | 4.284325 |
| ZT4_G3_OA1 | SSO | Amplitude (Hz) | 623.5885 | 430.3635 | 558.8202 | 25.99921 |
| ZT4_G3_OA10 | SSO | Frequency (Hz) | 412.5452 | 57.22712 | 397.7778 | 2.955196 |
| ZT4_G3_OA10 | SSO | Amplitude (Hz) | 414.074 | 129.6578 | 420.367 | 6.695499 |
| ZT12_G3_Ringer2 | SSO | Frequency (Hz) | 351.6738 | 23.56209 | 355.5556 | 0.842579 |
| ZT12_G3_Ringer2 | SSO | Amplitude (Hz) | 450.0125 | 235.6555 | 483.2234 | 8.427023 |
| ZT12_G3_OA1 | SSO | Frequency (Hz) | 407.3978 | 41.94953 | 397.7778 | 3.369468 |
| ZT12_G3_OA1 | SSO | Amplitude (Hz) | 386.1355 | 195.8682 | 390.6343 | 15.73251 |
| ZT12_G3_OA10 | SSO | Frequency (Hz) | 510.2116 | 25.81783 | 508.8889 | 3.983778 |
| ZT12_G3_OA10 | SSO | Amplitude (Hz) | 296.1106 | 201.9494 | 279.7756 | 31.16147 |

**Supplementary table 8: Summary of biophysical parameters extracted from antennal free fluctuations of SSO flagella in wildtype male mosquitoes upon exposure to octopamine**

**Supplementary table 9: Wilcoxon signed-rank test on free fluctuation analysis**

|  | .y. | group1 | group2 | p | p.adj | p.format | p.signif | method |
| --- | --- | --- | --- | --- | --- | --- | --- | --- |
| 1 | Frequency | ZT4\|Ringer2 | ZT4\|OA1 | 6.03E-106 | 8.40E-105 | <2e-16 | **** | Wilcoxon |
| 2 | Frequency | ZT4\|Ringer2 | ZT4\|OA10 | 1.63E-77 | 2.00E-76 | <2e-16 | **** | Wilcoxon |
| 3 | Frequency | ZT4\|Ringer2 | ZT12\|Ringer2 | 0.004813 | 0.014 | 0.0048 | ** | Wilcoxon |
| 4 | Frequency | ZT4\|Ringer2 | ZT12\|OA1 | 1.52E-63 | 1.50E-62 | <2e-16 | **** | Wilcoxon |
| 5 | Frequency | ZT4\|Ringer2 | ZT12\|OA10 | 1.37E-26 | 8.20E-26 | <2e-16 | **** | Wilcoxon |
| 6 | Frequency | ZT4\|OA1 | ZT4\|OA10 | 8.05E-43 | 7.20E-42 | <2e-16 | **** | Wilcoxon |
| 7 | Frequency | ZT4\|OA1 | ZT12\|Ringer2 | 1.10E-128 | 1.60E-127 | <2e-16 | **** | Wilcoxon |
| 8 | Frequency | ZT4\|OA1 | ZT12\|OA1 | 1.20E-31 | 9.60E-31 | <2e-16 | **** | Wilcoxon |
| 9 | Frequency | ZT4\|OA1 | ZT12\|OA10 | 0.366508 | 0.67 | 0.3665 | ns | Wilcoxon |
| 10 | Frequency | ZT4\|OA10 | ZT12\|Ringer2 | 6.57E-104 | 8.50E-103 | <2e-16 | **** | Wilcoxon |
| 11 | Frequency | ZT4\|OA10 | ZT12\|OA1 | 0.335243 | 0.67 | 0.3352 | ns | Wilcoxon |
| 12 | Frequency | ZT4\|OA10 | ZT12\|OA10 | 9.85E-22 | 4.90E-21 | <2e-16 | **** | Wilcoxon |
| 13 | Frequency | ZT12\|Ringer2 | ZT12\|OA1 | 8.85E-68 | 9.70E-67 | <2e-16 | **** | Wilcoxon |
| 14 | Frequency | ZT12\|Ringer2 | ZT12\|OA10 | 7.23E-28 | 5.10E-27 | <2e-16 | **** | Wilcoxon |
| 15 | Frequency | ZT12\|OA1 | ZT12\|OA10 | 3.27E-21 | 1.30E-20 | <2e-16 | **** | Wilcoxon |
| 16 | Amplitude | ZT4\|Ringer2 | ZT4\|OA1 | 0.004444 | 0.022 | 0.00444 | ** | Wilcoxon |
| 17 | Amplitude | ZT4\|Ringer2 | ZT4\|OA10 | 0.014661 | 0.039 | 0.01466 | * | Wilcoxon |
| 18 | Amplitude | ZT4\|Ringer2 | ZT12\|Ringer2 | 0.006153 | 0.025 | 0.00615 | ** | Wilcoxon |
| 19 | Amplitude | ZT4\|Ringer2 | ZT12\|OA1 | 1.56E-05 | 0.00012 | 1.60E-05 | **** | Wilcoxon |
| 20 | Amplitude | ZT4\|Ringer2 | ZT12\|OA10 | 1.27E-06 | 1.70E-05 | 1.30E-06 | **** | Wilcoxon |
| 21 | Amplitude | ZT4\|OA1 | ZT4\|OA10 | 4.46E-08 | 6.70E-07 | 4.50E-08 | **** | Wilcoxon |
| 22 | Amplitude | ZT4\|OA1 | ZT12\|Ringer2 | 2.85E-06 | 3.10E-05 | 2.80E-06 | **** | Wilcoxon |
| 23 | Amplitude | ZT4\|OA1 | ZT12\|OA1 | 1.75E-06 | 2.10E-05 | 1.80E-06 | **** | Wilcoxon |
| 24 | Amplitude | ZT4\|OA1 | ZT12\|OA10 | 6.33E-06 | 5.70E-05 | 6.30E-06 | **** | Wilcoxon |
| 25 | Amplitude | ZT4\|OA10 | ZT12\|Ringer2 | 4.54E-06 | 4.50E-05 | 4.50E-06 | **** | Wilcoxon |
| 26 | Amplitude | ZT4\|OA10 | ZT12\|OA1 | 0.136322 | 0.14 | 0.13632 | ns | Wilcoxon |
| 27 | Amplitude | ZT4\|OA10 | ZT12\|OA10 | 7.63E-07 | 1.10E-05 | 7.60E-07 | **** | Wilcoxon |
| 28 | Amplitude | ZT12\|Ringer2 | ZT12\|OA1 | 0.002111 | 0.013 | 0.00211 | ** | Wilcoxon |
| 29 | Amplitude | ZT12\|Ringer2 | ZT12\|OA10 | 0.000137 | 0.00096 | 0.00014 | *** | Wilcoxon |
| 30 | Amplitude | ZT12\|OA1 | ZT12\|OA10 | 0.012948 | 0.039 | 0.01295 | * | Wilcoxon |
