## Supplemental Table 10 and 11 for "A novel beta-adrenergic like octopamine receptor modulates the audition of malaria mosquitoes and serves as insecticide target"

**Supplementary table 10: Summary of biophysical parameters extracted from frequency-modulated sweep responses in wildtype male mosquitoes upon exposure to octopamine**

|  | condition | State | parameters | mean | sd | median | se |
| --- | --- | --- | --- | --- | --- | --- | --- |
| 1 | ZT4_G3_Ringer2 | SSO | F0m | 27.77139 | 12.05081 | 23.31792 | 0.734751 |
| 2 | ZT4_G3_Ringer2 | SSO | F0 | 353.9171 | 102.5214 | 343.1775 | 6.250839 |
| 3 | ZT4_G3_Ringer2 | SSO | damp_ratio | 0.053154 | 0.175631 | 0.146314 | 0.010708 |
| 4 | ZT4_G3_Ringer2 | SSO | Q-factor | 2.85E-07 | 3.02E-07 | 3.56E-07 | 1.84E-08 |
| 5 | ZT4_G3_OA1 | SSO | F0m | 12.03602 | 13.95092 | 13.59297 | 0.910058 |
| 6 | ZT4_G3_OA1 | SSO | F0 | 515.0869 | 79.40738 | 558.1755 | 5.179966 |
| 7 | ZT4_G3_OA1 | SSO | damp_ratio | -0.00659 | 0.169349 | -0.09079 | 0.011047 |
| 8 | ZT4_G3_OA1 | SSO | Q-factor | 8.41E-07 | 7.91E-07 | 6.36E-07 | 5.16E-08 |
| 9 | ZT4_G3_OA1 | Quiesc | F0m | -263.373 | 1109.224 | 10.04957 | 129.8249 |
| 10 | ZT4_G3_OA1 | Quiesc | F0 | 581.5119 | 338.982 | 434.1925 | 39.67484 |
| 11 | ZT4_G3_OA1 | Quiesc | damp_ratio | 0.282476 | 0.228976 | 0.236679 | 0.0268 |
| 12 | ZT4_G3_OA1 | Quiesc | Q-factor | 2.76E-06 | 8.73E-06 | 1.75E-07 | 1.02E-06 |
| 13 | ZT4_G3_OA10 | SSO | F0m | 28.42118 | 22.80354 | 38.7363 | 1.305727 |
| 14 | ZT4_G3_OA10 | SSO | F0 | 304.4006 | 361.6961 | 447.8948 | 20.71066 |
| 15 | ZT4_G3_OA10 | SSO | damp_ratio | -0.04861 | 0.189839 | -0.16058 | 0.01087 |
| 16 | ZT4_G3_OA10 | SSO | Q-factor | 8.24E-07 | 2.10E-06 | 3.36E-07 | 1.20E-07 |
| 17 | ZT4_G3_OA10 | Quiesc | F0m | 108.5403 | 557.6971 | 19.70692 | 52.00552 |
| 18 | ZT4_G3_OA10 | Quiesc | F0 | 629.632 | 191.6489 | 630.4844 | 17.87135 |
| 19 | ZT4_G3_OA10 | Quiesc | damp_ratio | 0.185879 | 0.27007 | 0.221907 | 0.025184 |
| 20 | ZT4_G3_OA10 | Quiesc | Q-factor | -9.36E-07 | 7.51E-06 | 6.45E-08 | 7.01E-07 |
| 21 | ZT12_G3_Ringer2 | SSO | F0m | 26.48756 | 10.41537 | 21.05205 | 0.423095 |
| 22 | ZT12_G3_Ringer2 | SSO | F0 | 314.5905 | 186.4026 | 349.9946 | 7.572089 |
| 23 | ZT12_G3_Ringer2 | SSO | damp_ratio | 0.028746 | 0.177441 | 0.158293 | 0.007208 |
| 24 | ZT12_G3_Ringer2 | SSO | Q-factor | 3.43E-07 | 2.16E-07 | 3.60E-07 | 8.76E-09 |
| 25 | ZT12_G3_Ringer2 | Quiesc | F0m | 28.55764 | 5.891399 | 26.0995 | 0.54466 |
| 26 | ZT12_G3_Ringer2 | Quiesc | F0 | 174.0825 | 324.3722 | 333.7481 | 29.98822 |
| 27 | ZT12_G3_Ringer2 | Quiesc | damp_ratio | 0.029724 | 0.196147 | 0.19101 | 0.018134 |
| 28 | ZT12_G3_Ringer2 | Quiesc | Q-factor | -5.93E-08 | 8.42E-08 | -2.55E-08 | 7.78E-09 |
| 29 | ZT12_G3_OA1 | SSO | F0m | -11.2426 | 356.0905 | 20.44885 | 32.92058 |
| 30 | ZT12_G3_OA1 | SSO | F0 | 402.8885 | 139.1633 | 382.3274 | 12.86565 |
| 31 | ZT12_G3_OA1 | SSO | damp_ratio | 0.038395 | 0.230228 | 0.157269 | 0.021285 |
| 32 | ZT12_G3_OA1 | SSO | Q-factor | 7.09E-07 | 2.88E-06 | 4.68E-07 | 2.67E-07 |
| 33 | ZT12_G3_OA1 | Quiesc | F0m | -16.2579 | 354.9364 | 14.3172 | 19.74919 |
| 34 | ZT12_G3_OA1 | Quiesc | F0 | 548.7785 | 679.0663 | 544.5224 | 37.78426 |
| 35 | ZT12_G3_OA1 | Quiesc | damp_ratio | 0.254112 | 2.080702 | 0.218421 | 0.115773 |
| 36 | ZT12_G3_OA1 | Quiesc | Q-factor | 2.66E-07 | 5.46E-07 | 7.84E-08 | 3.04E-08 |
| 37 | ZT12_G3_OA10 | SSO | F0m | 26.34493 | 16.7596 | 18.79456 | 2.83289 |
| 38 | ZT12_G3_OA10 | SSO | F0 | 549.1576 | 84.91681 | 502.9245 | 14.35356 |
| 39 | ZT12_G3_OA10 | SSO | damp_ratio | 0.174384 | 0.10123 | 0.185075 | 0.017111 |
| 40 | ZT12_G3_OA10 | SSO | Q-factor | 2.67E-07 | 5.78E-07 | 4.80E-07 | 9.77E-08 |
| 41 | ZT12_G3_OA10 | Quiesc | F0m | 22.88246 | 806.0903 | -0.22251 | 31.81385 |
| 42 | ZT12_G3_OA10 | Quiesc | F0 | 541.8071 | 708.4726 | 393.6804 | 27.96118 |
| 43 | ZT12_G3_OA10 | Quiesc | damp_ratio | 0.304689 | 1.981011 | 0.439472 | 0.078184 |
| 44 | ZT12_G3_OA10 | Quiesc | Q-factor | 3.80E-07 | 6.75E-06 | 4.10E-07 | 2.67E-07 |

**Supplementary table 11: Wilcoxon signed-rank test on frequency-modulated sweep analysis in males (pairwise comparisons).**

|  | .y. | group1 | group2 | p | p.adj | p.format | p.signif | method |
| --- | --- | --- | --- | --- | --- | --- | --- | --- |
| 1 | F0m | ZT4\|control\|SSO | ZT4\|1mM OA\|SSO | 3.15E-53 | 1.60E-51 | < 2e-16 | **** | Wilcoxon |
| 2 | F0m | ZT4\|control\|SSO | ZT4\|1mM OA\|Quiescent | 6.27E-33 | 2.60E-31 | < 2e-16 | **** | Wilcoxon |
| 3 | F0m | ZT4\|control\|SSO | ZT4\|10mM OA\|SSO | 1.57E-07 | 3.50E-06 | 1.60E-07 | **** | Wilcoxon |
| 4 | F0m | ZT4\|control\|SSO | ZT4\|10mM OA\|Quiescent | 7.06E-07 | 1.50E-05 | 7.10E-07 | **** | Wilcoxon |
| 5 | F0m | ZT4\|control\|SSO | ZT12\|control\|SSO | 0.02855 | 0.23 | 0.02855 | * | Wilcoxon |
| 6 | F0m | ZT4\|control\|SSO | ZT12\|control\|Quiescent | 0.000685 | 0.0082 | 0.00068 | *** | Wilcoxon |
| 7 | F0m | ZT4\|control\|SSO | ZT12\|1mM OA\|SSO | 5.72E-05 | 8.00E-04 | 5.70E-05 | **** | Wilcoxon |
| 8 | F0m | ZT4\|control\|SSO | ZT12\|1mM OA\|Quiescent | 3.06E-39 | 1.30E-37 | < 2e-16 | **** | Wilcoxon |
| 9 | F0m | ZT4\|control\|SSO | ZT12\|10mM OA\|SSO | 0.059736 | 0.36 | 0.05974 | ns | Wilcoxon |
| 10 | F0m | ZT4\|control\|SSO | ZT12\|10mM OA\|Quiescent | 3.32E-63 | 1.70E-61 | < 2e-16 | **** | Wilcoxon |
| 11 | F0m | ZT4\|1mM OA\|SSO | ZT4\|1mM OA\|Quiescent | 4.05E-05 | 0.00061 | 4.10E-05 | **** | Wilcoxon |
| 12 | F0m | ZT4\|1mM OA\|SSO | ZT4\|10mM OA\|SSO | 9.09E-50 | 4.40E-48 | < 2e-16 | **** | Wilcoxon |
| 13 | F0m | ZT4\|1mM OA\|SSO | ZT4\|10mM OA\|Quiescent | 0.000169 | 0.0022 | 0.00017 | *** | Wilcoxon |
| 14 | F0m | ZT4\|1mM OA\|SSO | ZT12\|control\|SSO | 1.04E-84 | 5.60E-83 | < 2e-16 | **** | Wilcoxon |
| 15 | F0m | ZT4\|1mM OA\|SSO | ZT12\|control\|Quiescent | 1.65E-49 | 7.80E-48 | < 2e-16 | **** | Wilcoxon |
| 16 | F0m | ZT4\|1mM OA\|SSO | ZT12\|1mM OA\|SSO | 3.85E-30 | 1.60E-28 | < 2e-16 | **** | Wilcoxon |
| 17 | F0m | ZT4\|1mM OA\|SSO | ZT12\|1mM OA\|Quiescent | 0.018034 | 0.18 | 0.01803 | * | Wilcoxon |
| 18 | F0m | ZT4\|1mM OA\|SSO | ZT12\|10mM OA\|SSO | 7.70E-10 | 2.10E-08 | 7.70E-10 | **** | Wilcoxon |
| 19 | F0m | ZT4\|1mM OA\|SSO | ZT12\|10mM OA\|Quiescent | 2.02E-22 | 7.10E-21 | < 2e-16 | **** | Wilcoxon |
| 20 | F0m | ZT4\|1mM OA\|Quiescent | ZT4\|10mM OA\|SSO | 1.17E-27 | 4.60E-26 | < 2e-16 | **** | Wilcoxon |
| 21 | F0m | ZT4\|1mM OA\|Quiescent | ZT4\|10mM OA\|Quiescent | 1.05E-05 | 0.00017 | 1.10E-05 | **** | Wilcoxon |
| 22 | F0m | ZT4\|1mM OA\|Quiescent | ZT12\|control\|SSO | 2.66E-35 | 1.10E-33 | < 2e-16 | **** | Wilcoxon |
| 23 | F0m | ZT4\|1mM OA\|Quiescent | ZT12\|control\|Quiescent | 1.40E-28 | 5.60E-27 | < 2e-16 | **** | Wilcoxon |
| 24 | F0m | ZT4\|1mM OA\|Quiescent | ZT12\|1mM OA\|SSO | 6.45E-23 | 2.30E-21 | < 2e-16 | **** | Wilcoxon |
| 25 | F0m | ZT4\|1mM OA\|Quiescent | ZT12\|1mM OA\|Quiescent | 0.007903 | 0.087 | 0.0079 | ** | Wilcoxon |
| 26 | F0m | ZT4\|1mM OA\|Quiescent | ZT12\|10mM OA\|SSO | 2.16E-12 | 6.90E-11 | 2.20E-12 | **** | Wilcoxon |
| 27 | F0m | ZT4\|1mM OA\|Quiescent | ZT12\|10mM OA\|Quiescent | 0.021742 | 0.2 | 0.02174 | * | Wilcoxon |
| 28 | F0m | ZT4\|10mM OA\|SSO | ZT4\|10mM OA\|Quiescent | 2.39E-10 | 6.90E-09 | 2.40E-10 | **** | Wilcoxon |
| 29 | F0m | ZT4\|10mM OA\|SSO | ZT12\|control\|SSO | 0.044202 | 0.31 | 0.0442 | * | Wilcoxon |
| 30 | F0m | ZT4\|10mM OA\|SSO | ZT12\|control\|Quiescent | 0.221555 | 0.7 | 0.22155 | ns | Wilcoxon |
| 31 | F0m | ZT4\|10mM OA\|SSO | ZT12\|1mM OA\|SSO | 4.25E-09 | 1.10E-07 | 4.30E-09 | **** | Wilcoxon |
| 32 | F0m | ZT4\|10mM OA\|SSO | ZT12\|1mM OA\|Quiescent | 1.64E-55 | 8.40E-54 | < 2e-16 | **** | Wilcoxon |
| 33 | F0m | ZT4\|10mM OA\|SSO | ZT12\|10mM OA\|SSO | 0.137974 | 0.69 | 0.13797 | ns | Wilcoxon |
| 34 | F0m | ZT4\|10mM OA\|SSO | ZT12\|10mM OA\|Quiescent | 2.97E-52 | 1.50E-50 | < 2e-16 | **** | Wilcoxon |
| 35 | F0m | ZT4\|10mM OA\|Quiescent | ZT12\|control\|SSO | 7.93E-07 | 1.60E-05 | 7.90E-07 | **** | Wilcoxon |
| 36 | F0m | ZT4\|10mM OA\|Quiescent | ZT12\|control\|Quiescent | 1.84E-10 | 5.50E-09 | 1.80E-10 | **** | Wilcoxon |
| 37 | F0m | ZT4\|10mM OA\|Quiescent | ZT12\|1mM OA\|SSO | 0.174533 | 0.7 | 0.17453 | ns | Wilcoxon |
| 38 | F0m | ZT4\|10mM OA\|Quiescent | ZT12\|1mM OA\|Quiescent | 4.01E-06 | 7.20E-05 | 4.00E-06 | **** | Wilcoxon |
| 39 | F0m | ZT4\|10mM OA\|Quiescent | ZT12\|10mM OA\|SSO | 0.288244 | 0.7 | 0.28824 | ns | Wilcoxon |
| 40 | F0m | ZT4\|10mM OA\|Quiescent | ZT12\|10mM OA\|Quiescent | 2.42E-10 | 6.90E-09 | 2.40E-10 | **** | Wilcoxon |
| 41 | F0m | ZT12\|control\|SSO | ZT12\|control\|Quiescent | 2.14E-15 | 7.30E-14 | 2.10E-15 | **** | Wilcoxon |
| 42 | F0m | ZT12\|control\|SSO | ZT12\|1mM OA\|SSO | 9.40E-09 | 2.30E-07 | 9.40E-09 | **** | Wilcoxon |
| 43 | F0m | ZT12\|control\|SSO | ZT12\|1mM OA\|Quiescent | 1.81E-78 | 9.60E-77 | < 2e-16 | **** | Wilcoxon |
| 44 | F0m | ZT12\|control\|SSO | ZT12\|10mM OA\|SSO | 1.13E-06 | 2.20E-05 | 1.10E-06 | **** | Wilcoxon |
| 45 | F0m | ZT12\|control\|SSO | ZT12\|10mM OA\|Quiescent | 2.37E-107 | 1.30E-105 | < 2e-16 | **** | Wilcoxon |
| 46 | F0m | ZT12\|control\|Quiescent | ZT12\|1mM OA\|SSO | 1.42E-11 | 4.40E-10 | 1.40E-11 | **** | Wilcoxon |
| 47 | F0m | ZT12\|control\|Quiescent | ZT12\|1mM OA\|Quiescent | 1.35E-41 | 6.20E-40 | < 2e-16 | **** | Wilcoxon |
| 48 | F0m | ZT12\|control\|Quiescent | ZT12\|10mM OA\|SSO | 8.55E-06 | 0.00015 | 8.60E-06 | **** | Wilcoxon |
| 49 | F0m | ZT12\|control\|Quiescent | ZT12\|10mM OA\|Quiescent | 2.42E-40 | 1.10E-38 | < 2e-16 | **** | Wilcoxon |
| 50 | F0m | ZT12\|1mM OA\|SSO | ZT12\|1mM OA\|Quiescent | 1.38E-25 | 5.20E-24 | < 2e-16 | **** | Wilcoxon |
| 51 | F0m | ZT12\|1mM OA\|SSO | ZT12\|10mM OA\|SSO | 0.30995 | 0.7 | 0.30995 | ns | Wilcoxon |
| 52 | F0m | ZT12\|1mM OA\|SSO | ZT12\|10mM OA\|Quiescent | 2.44E-25 | 9.00E-24 | < 2e-16 | **** | Wilcoxon |
| 53 | F0m | ZT12\|1mM OA\|Quiescent | ZT12\|10mM OA\|SSO | 6.77E-09 | 1.70E-07 | 6.80E-09 | **** | Wilcoxon |
| 54 | F0m | ZT12\|1mM OA\|Quiescent | ZT12\|10mM OA\|Quiescent | 5.41E-13 | 1.80E-11 | 5.40E-13 | **** | Wilcoxon |
| 55 | F0m | ZT12\|10mM OA\|SSO | ZT12\|10mM OA\|Quiescent | 2.80E-08 | 6.40E-07 | 2.80E-08 | **** | Wilcoxon |
| 56 | f0 | ZT4\|control\|SSO | ZT4\|1mM OA\|SSO | 1.99E-64 | 1.10E-62 | < 2e-16 | **** | Wilcoxon |
| 57 | f0 | ZT4\|control\|SSO | ZT4\|1mM OA\|Quiescent | 3.35E-25 | 1.40E-23 | < 2e-16 | **** | Wilcoxon |
| 58 | f0 | ZT4\|control\|SSO | ZT4\|10mM OA\|SSO | 4.16E-18 | 1.40E-16 | < 2e-16 | **** | Wilcoxon |
| 59 | f0 | ZT4\|control\|SSO | ZT4\|10mM OA\|Quiescent | 1.16E-46 | 5.90E-45 | < 2e-16 | **** | Wilcoxon |
| 60 | f0 | ZT4\|control\|SSO | ZT12\|control\|SSO | 0.189856 | 0.95 | 0.18986 | ns | Wilcoxon |
| 61 | f0 | ZT4\|control\|SSO | ZT12\|control\|Quiescent | 0.030401 | 0.3 | 0.0304 | * | Wilcoxon |
| 62 | f0 | ZT4\|control\|SSO | ZT12\|1mM OA\|SSO | 4.60E-09 | 8.70E-08 | 4.60E-09 | **** | Wilcoxon |
| 63 | f0 | ZT4\|control\|SSO | ZT12\|1mM OA\|Quiescent | 1.74E-43 | 8.70E-42 | < 2e-16 | **** | Wilcoxon |
| 64 | f0 | ZT4\|control\|SSO | ZT12\|10mM OA\|SSO | 6.44E-22 | 2.40E-20 | < 2e-16 | **** | Wilcoxon |
| 65 | f0 | ZT4\|control\|SSO | ZT12\|10mM OA\|Quiescent | 1.81E-13 | 4.20E-12 | 1.80E-13 | **** | Wilcoxon |
| 66 | f0 | ZT4\|1mM OA\|SSO | ZT4\|1mM OA\|Quiescent | 0.275351 | 0.95 | 0.27535 | ns | Wilcoxon |
| 67 | f0 | ZT4\|1mM OA\|SSO | ZT4\|10mM OA\|SSO | 6.39E-24 | 2.60E-22 | < 2e-16 | **** | Wilcoxon |
| 68 | f0 | ZT4\|1mM OA\|SSO | ZT4\|10mM OA\|Quiescent | 1.10E-22 | 4.40E-21 | < 2e-16 | **** | Wilcoxon |
| 69 | f0 | ZT4\|1mM OA\|SSO | ZT12\|control\|SSO | 3.32E-96 | 1.80E-94 | < 2e-16 | **** | Wilcoxon |
| 70 | f0 | ZT4\|1mM OA\|SSO | ZT12\|control\|Quiescent | 5.77E-39 | 2.80E-37 | < 2e-16 | **** | Wilcoxon |
| 71 | f0 | ZT4\|1mM OA\|SSO | ZT12\|1mM OA\|SSO | 1.33E-33 | 6.40E-32 | < 2e-16 | **** | Wilcoxon |
| 72 | f0 | ZT4\|1mM OA\|SSO | ZT12\|1mM OA\|Quiescent | 0.512352 | 0.95 | 0.51235 | ns | Wilcoxon |
| 73 | f0 | ZT4\|1mM OA\|SSO | ZT12\|10mM OA\|SSO | 0.191447 | 0.95 | 0.19145 | ns | Wilcoxon |
| 74 | f0 | ZT4\|1mM OA\|SSO | ZT12\|10mM OA\|Quiescent | 1.39E-09 | 2.80E-08 | 1.40E-09 | **** | Wilcoxon |
| 75 | f0 | ZT4\|1mM OA\|Quiescent | ZT4\|10mM OA\|SSO | 0.000107 | 0.0017 | 0.00011 | *** | Wilcoxon |
| 76 | f0 | ZT4\|1mM OA\|Quiescent | ZT4\|10mM OA\|Quiescent | 3.64E-06 | 6.20E-05 | 3.60E-06 | **** | Wilcoxon |
| 77 | f0 | ZT4\|1mM OA\|Quiescent | ZT12\|control\|SSO | 1.32E-27 | 5.80E-26 | < 2e-16 | **** | Wilcoxon |
| 78 | f0 | ZT4\|1mM OA\|Quiescent | ZT12\|control\|Quiescent | 3.30E-15 | 8.60E-14 | 3.30E-15 | **** | Wilcoxon |
| 79 | f0 | ZT4\|1mM OA\|Quiescent | ZT12\|1mM OA\|SSO | 3.09E-11 | 6.50E-10 | 3.10E-11 | **** | Wilcoxon |
| 80 | f0 | ZT4\|1mM OA\|Quiescent | ZT12\|1mM OA\|Quiescent | 0.442024 | 0.95 | 0.44202 | ns | Wilcoxon |
| 81 | f0 | ZT4\|1mM OA\|Quiescent | ZT12\|10mM OA\|SSO | 0.002701 | 0.032 | 0.0027 | ** | Wilcoxon |
| 82 | f0 | ZT4\|1mM OA\|Quiescent | ZT12\|10mM OA\|Quiescent | 0.000979 | 0.014 | 0.00098 | *** | Wilcoxon |
| 83 | f0 | ZT4\|10mM OA\|SSO | ZT4\|10mM OA\|Quiescent | 1.53E-29 | 6.90E-28 | < 2e-16 | **** | Wilcoxon |
| 84 | f0 | ZT4\|10mM OA\|SSO | ZT12\|control\|SSO | 1.40E-11 | 3.10E-10 | 1.40E-11 | **** | Wilcoxon |
| 85 | f0 | ZT4\|10mM OA\|SSO | ZT12\|control\|Quiescent | 5.69E-15 | 1.40E-13 | 5.70E-15 | **** | Wilcoxon |
| 86 | f0 | ZT4\|10mM OA\|SSO | ZT12\|1mM OA\|SSO | 0.003674 | 0.04 | 0.00367 | ** | Wilcoxon |
| 87 | f0 | ZT4\|10mM OA\|SSO | ZT12\|1mM OA\|Quiescent | 2.27E-14 | 5.50E-13 | 2.30E-14 | **** | Wilcoxon |
| 88 | f0 | ZT4\|10mM OA\|SSO | ZT12\|10mM OA\|SSO | 6.97E-07 | 1.30E-05 | 7.00E-07 | **** | Wilcoxon |
| 89 | f0 | ZT4\|10mM OA\|SSO | ZT12\|10mM OA\|Quiescent | 0.148656 | 0.89 | 0.14866 | ns | Wilcoxon |
| 90 | f0 | ZT4\|10mM OA\|Quiescent | ZT12\|control\|SSO | 4.81E-57 | 2.50E-55 | < 2e-16 | **** | Wilcoxon |
| 91 | f0 | ZT4\|10mM OA\|Quiescent | ZT12\|control\|Quiescent | 9.68E-33 | 4.50E-31 | < 2e-16 | **** | Wilcoxon |
| 92 | f0 | ZT4\|10mM OA\|Quiescent | ZT12\|1mM OA\|SSO | 4.17E-30 | 1.90E-28 | < 2e-16 | **** | Wilcoxon |
| 93 | f0 | ZT4\|10mM OA\|Quiescent | ZT12\|1mM OA\|Quiescent | 2.23E-19 | 7.80E-18 | < 2e-16 | **** | Wilcoxon |
| 94 | f0 | ZT4\|10mM OA\|Quiescent | ZT12\|10mM OA\|SSO | 0.048004 | 0.43 | 0.048 | * | Wilcoxon |
| 95 | f0 | ZT4\|10mM OA\|Quiescent | ZT12\|10mM OA\|Quiescent | 8.93E-17 | 2.70E-15 | < 2e-16 | **** | Wilcoxon |
| 96 | f0 | ZT12\|control\|SSO | ZT12\|control\|Quiescent | 1.50E-22 | 5.70E-21 | < 2e-16 | **** | Wilcoxon |
| 97 | f0 | ZT12\|control\|SSO | ZT12\|1mM OA\|SSO | 2.79E-17 | 8.60E-16 | < 2e-16 | **** | Wilcoxon |
| 98 | f0 | ZT12\|control\|SSO | ZT12\|1mM OA\|Quiescent | 1.65E-62 | 8.70E-61 | < 2e-16 | **** | Wilcoxon |
| 99 | f0 | ZT12\|control\|SSO | ZT12\|10mM OA\|SSO | 1.40E-22 | 5.50E-21 | < 2e-16 | **** | Wilcoxon |
| 100 | f0 | ZT12\|control\|SSO | ZT12\|10mM OA\|Quiescent | 1.65E-16 | 4.80E-15 | < 2e-16 | **** | Wilcoxon |
| 101 | f0 | ZT12\|control\|Quiescent | ZT12\|1mM OA\|SSO | 1.90E-15 | 5.30E-14 | 1.90E-15 | **** | Wilcoxon |
| 102 | f0 | ZT12\|control\|Quiescent | ZT12\|1mM OA\|Quiescent | 4.94E-21 | 1.80E-19 | < 2e-16 | **** | Wilcoxon |
| 103 | f0 | ZT12\|control\|Quiescent | ZT12\|10mM OA\|SSO | 5.73E-19 | 1.90E-17 | < 2e-16 | **** | Wilcoxon |
| 104 | f0 | ZT12\|control\|Quiescent | ZT12\|10mM OA\|Quiescent | 2.13E-15 | 5.70E-14 | 2.10E-15 | **** | Wilcoxon |
| 105 | f0 | ZT12\|1mM OA\|SSO | ZT12\|1mM OA\|Quiescent | 1.44E-24 | 6.10E-23 | < 2e-16 | **** | Wilcoxon |
| 106 | f0 | ZT12\|1mM OA\|SSO | ZT12\|10mM OA\|SSO | 8.96E-18 | 2.90E-16 | < 2e-16 | **** | Wilcoxon |
| 107 | f0 | ZT12\|1mM OA\|SSO | ZT12\|10mM OA\|Quiescent | 0.095369 | 0.76 | 0.09537 | ns | Wilcoxon |
| 108 | f0 | ZT12\|1mM OA\|Quiescent | ZT12\|10mM OA\|SSO | 0.12214 | 0.85 | 0.12214 | ns | Wilcoxon |
| 109 | f0 | ZT12\|1mM OA\|Quiescent | ZT12\|10mM OA\|Quiescent | 0.000149 | 0.0022 | 0.00015 | *** | Wilcoxon |
| 110 | f0 | ZT12\|10mM OA\|SSO | ZT12\|10mM OA\|Quiescent | 0.001852 | 0.024 | 0.00185 | ** | Wilcoxon |
| 111 | Q | ZT4\|control\|SSO | ZT4\|1mM OA\|SSO | 1.26E-26 | 5.90E-25 | < 2e-16 | **** | Wilcoxon |
| 112 | Q | ZT4\|control\|SSO | ZT4\|1mM OA\|Quiescent | 4.70E-14 | 1.50E-12 | 4.70E-14 | **** | Wilcoxon |
| 113 | Q | ZT4\|control\|SSO | ZT4\|10mM OA\|SSO | 0.006123 | 0.055 | 0.00612 | ** | Wilcoxon |
| 114 | Q | ZT4\|control\|SSO | ZT4\|10mM OA\|Quiescent | 5.22E-14 | 1.60E-12 | 5.20E-14 | **** | Wilcoxon |
| 115 | Q | ZT4\|control\|SSO | ZT12\|control\|SSO | 0.558653 | 1 | 0.55865 | ns | Wilcoxon |
| 116 | Q | ZT4\|control\|SSO | ZT12\|control\|Quiescent | 1.05E-05 | 0.00022 | 1.10E-05 | **** | Wilcoxon |
| 117 | Q | ZT4\|control\|SSO | ZT12\|1mM OA\|SSO | 9.20E-05 | 0.0016 | 9.20E-05 | **** | Wilcoxon |
| 118 | Q | ZT4\|control\|SSO | ZT12\|1mM OA\|Quiescent | 7.90E-11 | 2.40E-09 | 7.90E-11 | **** | Wilcoxon |
| 119 | Q | ZT4\|control\|SSO | ZT12\|10mM OA\|SSO | 0.336658 | 1 | 0.33666 | ns | Wilcoxon |
| 120 | Q | ZT4\|control\|SSO | ZT12\|10mM OA\|Quiescent | 4.46E-54 | 2.40E-52 | < 2e-16 | **** | Wilcoxon |
| 121 | Q | ZT4\|1mM OA\|SSO | ZT4\|1mM OA\|Quiescent | 5.45E-23 | 2.50E-21 | < 2e-16 | **** | Wilcoxon |
| 122 | Q | ZT4\|1mM OA\|SSO | ZT4\|10mM OA\|SSO | 8.12E-16 | 2.80E-14 | 8.10E-16 | **** | Wilcoxon |
| 123 | Q | ZT4\|1mM OA\|SSO | ZT4\|10mM OA\|Quiescent | 3.65E-21 | 1.50E-19 | < 2e-16 | **** | Wilcoxon |
| 124 | Q | ZT4\|1mM OA\|SSO | ZT12\|control\|SSO | 6.56E-31 | 3.20E-29 | < 2e-16 | **** | Wilcoxon |
| 125 | Q | ZT4\|1mM OA\|SSO | ZT12\|control\|Quiescent | 4.00E-35 | 2.00E-33 | < 2e-16 | **** | Wilcoxon |
| 126 | Q | ZT4\|1mM OA\|SSO | ZT12\|1mM OA\|SSO | 1.92E-20 | 7.90E-19 | < 2e-16 | **** | Wilcoxon |
| 127 | Q | ZT4\|1mM OA\|SSO | ZT12\|1mM OA\|Quiescent | 1.12E-17 | 4.00E-16 | < 2e-16 | **** | Wilcoxon |
| 128 | Q | ZT4\|1mM OA\|SSO | ZT12\|10mM OA\|SSO | 3.92E-08 | 1.00E-06 | 3.90E-08 | **** | Wilcoxon |
| 129 | Q | ZT4\|1mM OA\|SSO | ZT12\|10mM OA\|Quiescent | 2.29E-53 | 1.20E-51 | < 2e-16 | **** | Wilcoxon |
| 130 | Q | ZT4\|1mM OA\|Quiescent | ZT4\|10mM OA\|SSO | 2.91E-16 | 1.00E-14 | 2.90E-16 | **** | Wilcoxon |
| 131 | Q | ZT4\|1mM OA\|Quiescent | ZT4\|10mM OA\|Quiescent | 0.881944 | 1 | 0.88194 | ns | Wilcoxon |
| 132 | Q | ZT4\|1mM OA\|Quiescent | ZT12\|control\|SSO | 2.14E-15 | 7.10E-14 | 2.10E-15 | **** | Wilcoxon |
| 133 | Q | ZT4\|1mM OA\|Quiescent | ZT12\|control\|Quiescent | 3.83E-07 | 9.20E-06 | 3.80E-07 | **** | Wilcoxon |
| 134 | Q | ZT4\|1mM OA\|Quiescent | ZT12\|1mM OA\|SSO | 0.001347 | 0.018 | 0.00135 | ** | Wilcoxon |
| 135 | Q | ZT4\|1mM OA\|Quiescent | ZT12\|1mM OA\|Quiescent | 0.007616 | 0.061 | 0.00762 | ** | Wilcoxon |
| 136 | Q | ZT4\|1mM OA\|Quiescent | ZT12\|10mM OA\|SSO | 0.00282 | 0.031 | 0.00282 | ** | Wilcoxon |
| 137 | Q | ZT4\|1mM OA\|Quiescent | ZT12\|10mM OA\|Quiescent | 1.96E-05 | 0.00037 | 2.00E-05 | **** | Wilcoxon |
| 138 | Q | ZT4\|10mM OA\|SSO | ZT4\|10mM OA\|Quiescent | 1.47E-18 | 5.60E-17 | < 2e-16 | **** | Wilcoxon |
| 139 | Q | ZT4\|10mM OA\|SSO | ZT12\|control\|SSO | 0.000771 | 0.011 | 0.00077 | *** | Wilcoxon |
| 140 | Q | ZT4\|10mM OA\|SSO | ZT12\|control\|Quiescent | 1.48E-33 | 7.40E-32 | < 2e-16 | **** | Wilcoxon |
| 141 | Q | ZT4\|10mM OA\|SSO | ZT12\|1mM OA\|SSO | 2.93E-09 | 8.20E-08 | 2.90E-09 | **** | Wilcoxon |
| 142 | Q | ZT4\|10mM OA\|SSO | ZT12\|1mM OA\|Quiescent | 3.58E-20 | 1.40E-18 | < 2e-16 | **** | Wilcoxon |
| 143 | Q | ZT4\|10mM OA\|SSO | ZT12\|10mM OA\|SSO | 1.19E-05 | 0.00024 | 1.20E-05 | **** | Wilcoxon |
| 144 | Q | ZT4\|10mM OA\|SSO | ZT12\|10mM OA\|Quiescent | 1.21E-59 | 6.50E-58 | < 2e-16 | **** | Wilcoxon |
| 145 | Q | ZT4\|10mM OA\|Quiescent | ZT12\|control\|SSO | 1.70E-18 | 6.30E-17 | < 2e-16 | **** | Wilcoxon |
| 146 | Q | ZT4\|10mM OA\|Quiescent | ZT12\|control\|Quiescent | 7.89E-07 | 1.80E-05 | 7.90E-07 | **** | Wilcoxon |
| 147 | Q | ZT4\|10mM OA\|Quiescent | ZT12\|1mM OA\|SSO | 0.000231 | 0.0037 | 0.00023 | *** | Wilcoxon |
| 148 | Q | ZT4\|10mM OA\|Quiescent | ZT12\|1mM OA\|Quiescent | 0.023009 | 0.12 | 0.02301 | * | Wilcoxon |
| 149 | Q | ZT4\|10mM OA\|Quiescent | ZT12\|10mM OA\|SSO | 0.001984 | 0.024 | 0.00198 | ** | Wilcoxon |
| 150 | Q | ZT4\|10mM OA\|Quiescent | ZT12\|10mM OA\|Quiescent | 7.60E-08 | 1.90E-06 | 7.60E-08 | **** | Wilcoxon |
| 151 | Q | ZT12\|control\|SSO | ZT12\|control\|Quiescent | 6.94E-20 | 2.70E-18 | < 2e-16 | **** | Wilcoxon |
| 152 | Q | ZT12\|control\|SSO | ZT12\|1mM OA\|SSO | 5.40E-06 | 0.00012 | 5.40E-06 | **** | Wilcoxon |
| 153 | Q | ZT12\|control\|SSO | ZT12\|1mM OA\|Quiescent | 1.44E-22 | 6.30E-21 | < 2e-16 | **** | Wilcoxon |
| 154 | Q | ZT12\|control\|SSO | ZT12\|10mM OA\|SSO | 0.003177 | 0.032 | 0.00318 | ** | Wilcoxon |
| 155 | Q | ZT12\|control\|SSO | ZT12\|10mM OA\|Quiescent | 2.87E-89 | 1.60E-87 | < 2e-16 | **** | Wilcoxon |
| 156 | Q | ZT12\|control\|Quiescent | ZT12\|1mM OA\|SSO | 0.000466 | 0.007 | 0.00047 | *** | Wilcoxon |
| 157 | Q | ZT12\|control\|Quiescent | ZT12\|1mM OA\|Quiescent | 2.73E-08 | 7.40E-07 | 2.70E-08 | **** | Wilcoxon |
| 158 | Q | ZT12\|control\|Quiescent | ZT12\|10mM OA\|SSO | 3.22E-05 | 0.00058 | 3.20E-05 | **** | Wilcoxon |
| 159 | Q | ZT12\|control\|Quiescent | ZT12\|10mM OA\|Quiescent | 1.21E-26 | 5.80E-25 | < 2e-16 | **** | Wilcoxon |
| 160 | Q | ZT12\|1mM OA\|SSO | ZT12\|1mM OA\|Quiescent | 0.018242 | 0.12 | 0.01824 | * | Wilcoxon |
| 161 | Q | ZT12\|1mM OA\|SSO | ZT12\|10mM OA\|SSO | 0.587354 | 1 | 0.58735 | ns | Wilcoxon |
| 162 | Q | ZT12\|1mM OA\|SSO | ZT12\|10mM OA\|Quiescent | 3.27E-21 | 1.40E-19 | < 2e-16 | **** | Wilcoxon |
| 163 | Q | ZT12\|1mM OA\|Quiescent | ZT12\|10mM OA\|SSO | 0.01716 | 0.12 | 0.01716 | * | Wilcoxon |
| 164 | Q | ZT12\|1mM OA\|Quiescent | ZT12\|10mM OA\|Quiescent | 3.58E-24 | 1.60E-22 | < 2e-16 | **** | Wilcoxon |
| 165 | Q | ZT12\|10mM OA\|SSO | ZT12\|10mM OA\|Quiescent | 1.58E-09 | 4.60E-08 | 1.60E-09 | **** | Wilcoxon |
| 166 | Q | ZT4\|control\|SSO | ZT4\|1mM OA\|SSO | 1.26E-26 | 5.90E-25 | < 2e-16 | **** | Wilcoxon |
| 167 | Q | ZT4\|control\|SSO | ZT4\|1mM OA\|Quiescent | 4.70E-14 | 1.50E-12 | 4.70E-14 | **** | Wilcoxon |
| 168 | Q | ZT4\|control\|SSO | ZT4\|10mM OA\|SSO | 0.006123 | 0.055 | 0.00612 | ** | Wilcoxon |
| 169 | Q | ZT4\|control\|SSO | ZT4\|10mM OA\|Quiescent | 5.22E-14 | 1.60E-12 | 5.20E-14 | **** | Wilcoxon |
| 170 | Q | ZT4\|control\|SSO | ZT12\|control\|SSO | 0.558653 | 1 | 0.55865 | ns | Wilcoxon |
| 171 | Q | ZT4\|control\|SSO | ZT12\|control\|Quiescent | 1.05E-05 | 0.00022 | 1.10E-05 | **** | Wilcoxon |
| 172 | Q | ZT4\|control\|SSO | ZT12\|1mM OA\|SSO | 9.20E-05 | 0.0016 | 9.20E-05 | **** | Wilcoxon |
| 173 | Q | ZT4\|control\|SSO | ZT12\|1mM OA\|Quiescent | 7.90E-11 | 2.40E-09 | 7.90E-11 | **** | Wilcoxon |
| 174 | Q | ZT4\|control\|SSO | ZT12\|10mM OA\|SSO | 0.336658 | 1 | 0.33666 | ns | Wilcoxon |
| 175 | Q | ZT4\|control\|SSO | ZT12\|10mM OA\|Quiescent | 4.46E-54 | 2.40E-52 | < 2e-16 | **** | Wilcoxon |
| 176 | Q | ZT4\|1mM OA\|SSO | ZT4\|1mM OA\|Quiescent | 5.45E-23 | 2.50E-21 | < 2e-16 | **** | Wilcoxon |
| 177 | Q | ZT4\|1mM OA\|SSO | ZT4\|10mM OA\|SSO | 8.12E-16 | 2.80E-14 | 8.10E-16 | **** | Wilcoxon |
| 178 | Q | ZT4\|1mM OA\|SSO | ZT4\|10mM OA\|Quiescent | 3.65E-21 | 1.50E-19 | < 2e-16 | **** | Wilcoxon |
| 179 | Q | ZT4\|1mM OA\|SSO | ZT12\|control\|SSO | 6.56E-31 | 3.20E-29 | < 2e-16 | **** | Wilcoxon |
| 180 | Q | ZT4\|1mM OA\|SSO | ZT12\|control\|Quiescent | 4.00E-35 | 2.00E-33 | < 2e-16 | **** | Wilcoxon |
| 181 | Q | ZT4\|1mM OA\|SSO | ZT12\|1mM OA\|SSO | 1.92E-20 | 7.90E-19 | < 2e-16 | **** | Wilcoxon |
| 182 | Q | ZT4\|1mM OA\|SSO | ZT12\|1mM OA\|Quiescent | 1.12E-17 | 4.00E-16 | < 2e-16 | **** | Wilcoxon |
| 183 | Q | ZT4\|1mM OA\|SSO | ZT12\|10mM OA\|SSO | 3.92E-08 | 1.00E-06 | 3.90E-08 | **** | Wilcoxon |
| 184 | Q | ZT4\|1mM OA\|SSO | ZT12\|10mM OA\|Quiescent | 2.29E-53 | 1.20E-51 | < 2e-16 | **** | Wilcoxon |
| 185 | Q | ZT4\|1mM OA\|Quiescent | ZT4\|10mM OA\|SSO | 2.91E-16 | 1.00E-14 | 2.90E-16 | **** | Wilcoxon |
| 186 | Q | ZT4\|1mM OA\|Quiescent | ZT4\|10mM OA\|Quiescent | 0.881944 | 1 | 0.88194 | ns | Wilcoxon |
| 187 | Q | ZT4\|1mM OA\|Quiescent | ZT12\|control\|SSO | 2.14E-15 | 7.10E-14 | 2.10E-15 | **** | Wilcoxon |
| 188 | Q | ZT4\|1mM OA\|Quiescent | ZT12\|control\|Quiescent | 3.83E-07 | 9.20E-06 | 3.80E-07 | **** | Wilcoxon |
| 189 | Q | ZT4\|1mM OA\|Quiescent | ZT12\|1mM OA\|SSO | 0.001347 | 0.018 | 0.00135 | ** | Wilcoxon |
| 190 | Q | ZT4\|1mM OA\|Quiescent | ZT12\|1mM OA\|Quiescent | 0.007616 | 0.061 | 0.00762 | ** | Wilcoxon |
| 191 | Q | ZT4\|1mM OA\|Quiescent | ZT12\|10mM OA\|SSO | 0.00282 | 0.031 | 0.00282 | ** | Wilcoxon |
| 192 | Q | ZT4\|1mM OA\|Quiescent | ZT12\|10mM OA\|Quiescent | 1.96E-05 | 0.00037 | 2.00E-05 | **** | Wilcoxon |
| 193 | Q | ZT4\|10mM OA\|SSO | ZT4\|10mM OA\|Quiescent | 1.47E-18 | 5.60E-17 | < 2e-16 | **** | Wilcoxon |
| 194 | Q | ZT4\|10mM OA\|SSO | ZT12\|control\|SSO | 0.000771 | 0.011 | 0.00077 | *** | Wilcoxon |
| 195 | Q | ZT4\|10mM OA\|SSO | ZT12\|control\|Quiescent | 1.48E-33 | 7.40E-32 | < 2e-16 | **** | Wilcoxon |
| 196 | Q | ZT4\|10mM OA\|SSO | ZT12\|1mM OA\|SSO | 2.93E-09 | 8.20E-08 | 2.90E-09 | **** | Wilcoxon |
| 197 | Q | ZT4\|10mM OA\|SSO | ZT12\|1mM OA\|Quiescent | 3.58E-20 | 1.40E-18 | < 2e-16 | **** | Wilcoxon |
| 198 | Q | ZT4\|10mM OA\|SSO | ZT12\|10mM OA\|SSO | 1.19E-05 | 0.00024 | 1.20E-05 | **** | Wilcoxon |
| 199 | Q | ZT4\|10mM OA\|SSO | ZT12\|10mM OA\|Quiescent | 1.21E-59 | 6.50E-58 | < 2e-16 | **** | Wilcoxon |
| 200 | Q | ZT4\|10mM OA\|Quiescent | ZT12\|control\|SSO | 1.70E-18 | 6.30E-17 | < 2e-16 | **** | Wilcoxon |
| 201 | Q | ZT4\|10mM OA\|Quiescent | ZT12\|control\|Quiescent | 7.89E-07 | 1.80E-05 | 7.90E-07 | **** | Wilcoxon |
| 202 | Q | ZT4\|10mM OA\|Quiescent | ZT12\|1mM OA\|SSO | 0.000231 | 0.0037 | 0.00023 | *** | Wilcoxon |
| 203 | Q | ZT4\|10mM OA\|Quiescent | ZT12\|1mM OA\|Quiescent | 0.023009 | 0.12 | 0.02301 | * | Wilcoxon |
| 204 | Q | ZT4\|10mM OA\|Quiescent | ZT12\|10mM OA\|SSO | 0.001984 | 0.024 | 0.00198 | ** | Wilcoxon |
| 205 | Q | ZT4\|10mM OA\|Quiescent | ZT12\|10mM OA\|Quiescent | 7.60E-08 | 1.90E-06 | 7.60E-08 | **** | Wilcoxon |
| 206 | Q | ZT12\|control\|SSO | ZT12\|control\|Quiescent | 6.94E-20 | 2.70E-18 | < 2e-16 | **** | Wilcoxon |
| 207 | Q | ZT12\|control\|SSO | ZT12\|1mM OA\|SSO | 5.40E-06 | 0.00012 | 5.40E-06 | **** | Wilcoxon |
| 208 | Q | ZT12\|control\|SSO | ZT12\|1mM OA\|Quiescent | 1.44E-22 | 6.30E-21 | < 2e-16 | **** | Wilcoxon |
| 209 | Q | ZT12\|control\|SSO | ZT12\|10mM OA\|SSO | 0.003177 | 0.032 | 0.00318 | ** | Wilcoxon |
| 210 | Q | ZT12\|control\|SSO | ZT12\|10mM OA\|Quiescent | 2.87E-89 | 1.60E-87 | < 2e-16 | **** | Wilcoxon |
| 211 | Q | ZT12\|control\|Quiescent | ZT12\|1mM OA\|SSO | 0.000466 | 0.007 | 0.00047 | *** | Wilcoxon |
| 212 | Q | ZT12\|control\|Quiescent | ZT12\|1mM OA\|Quiescent | 2.73E-08 | 7.40E-07 | 2.70E-08 | **** | Wilcoxon |
| 213 | Q | ZT12\|control\|Quiescent | ZT12\|10mM OA\|SSO | 3.22E-05 | 0.00058 | 3.20E-05 | **** | Wilcoxon |
| 214 | Q | ZT12\|control\|Quiescent | ZT12\|10mM OA\|Quiescent | 1.21E-26 | 5.80E-25 | < 2e-16 | **** | Wilcoxon |
| 215 | Q | ZT12\|1mM OA\|SSO | ZT12\|1mM OA\|Quiescent | 0.018242 | 0.12 | 0.01824 | * | Wilcoxon |
| 216 | Q | ZT12\|1mM OA\|SSO | ZT12\|10mM OA\|SSO | 0.587354 | 1 | 0.58735 | ns | Wilcoxon |
| 217 | Q | ZT12\|1mM OA\|SSO | ZT12\|10mM OA\|Quiescent | 3.27E-21 | 1.40E-19 | < 2e-16 | **** | Wilcoxon |
| 218 | Q | ZT12\|1mM OA\|Quiescent | ZT12\|10mM OA\|SSO | 0.01716 | 0.12 | 0.01716 | * | Wilcoxon |
| 219 | Q | ZT12\|1mM OA\|Quiescent | ZT12\|10mM OA\|Quiescent | 3.58E-24 | 1.60E-22 | < 2e-16 | **** | Wilcoxon |
| 220 | Q | ZT12\|10mM OA\|SSO | ZT12\|10mM OA\|Quiescent | 1.58E-09 | 4.60E-08 | 1.60E-09 | **** | Wilcoxon |
