## Supplemental Table 16 and 17 for "A novel beta-adrenergic like octopamine receptor modulates the audition of malaria mosquitoes and serves as insecticide target"

**Supplementary table 16: Summary of biophysical parameters extracted from antennal free fluctuations of SSO flagella in octopamine receptor mutant male mosquitoes upon exposure to octopamine**

| condition | State | parameter | mean | sd | median | se |
| --- | --- | --- | --- | --- | --- | --- |
| ZT12_G3_OA1 | SSO | Frequency | 407.3978 | 41.94953 | 397.7778 | 3.369468 |
| ZT12_G3_OA1 | SSO | Amplitude | 386.1355 | 195.8682 | 390.6343 | 15.73251 |
| ZT12_45_OA1 | SSO | Frequency | 442.7131 | 77.18562 | 428.8889 | 4.217101 |
| ZT12_45_OA1 | SSO | Amplitude | 642.7155 | 388.6638 | 622.8234 | 21.23497 |
| ZT12_2886_OA1 | SSO | Frequency | 349.041 | 14.16409 | 344.4444 | 0.665487 |
| ZT12_2886_OA1 | SSO | Amplitude | 754.3368 | 352.145 | 728.0937 | 16.54521 |

**Supplementary table 17: Wilcoxon signed-rank test on free fluctuation analysis in octopamine receptor mutant male mosquitoes**

|  | .y. | group1 | group2 | p | p.adj | p.format | p.signif | method |
| --- | --- | --- | --- | --- | --- | --- | --- | --- |
| 1 | Frequency | G3\|baseline | AGAP000045-\|baseline | 0.225387 | 0.23 | 0.22539 | ns | Wilcoxon |
| 2 | Frequency | G3\|baseline | AGAP002886-\|baseline | 3.65E-64 | 4.70E-63 | < 2e-16 | **** | Wilcoxon |
| 3 | Frequency | G3\|baseline | G3\|OA1 | 7.19E-42 | 7.20E-41 | < 2e-16 | **** | Wilcoxon |
| 4 | Frequency | G3\|baseline | AGAP000045-\|OA1 | 1.19E-45 | 1.30E-44 | < 2e-16 | **** | Wilcoxon |
| 5 | Frequency | G3\|baseline | AGAP002886-\|OA1 | 1.77E-16 | 7.10E-16 | < 2e-16 | **** | Wilcoxon |
| 6 | Frequency | AGAP000045-\|baseline | AGAP002886-\|baseline | 3.91E-32 | 2.70E-31 | < 2e-16 | **** | Wilcoxon |
| 7 | Frequency | AGAP000045-\|baseline | G3\|OA1 | 9.28E-34 | 7.40E-33 | < 2e-16 | **** | Wilcoxon |
| 8 | Frequency | AGAP000045-\|baseline | AGAP000045-\|OA1 | 1.99E-40 | 1.80E-39 | < 2e-16 | **** | Wilcoxon |
| 9 | Frequency | AGAP000045-\|baseline | AGAP002886-\|OA1 | 1.77E-17 | 8.80E-17 | < 2e-16 | **** | Wilcoxon |
| 10 | Frequency | AGAP002886-\|baseline | G3\|OA1 | 1.91E-08 | 5.70E-08 | 1.90E-08 | **** | Wilcoxon |
| 11 | Frequency | AGAP002886-\|baseline | AGAP000045-\|OA1 | 1.29E-24 | 7.80E-24 | < 2e-16 | **** | Wilcoxon |
| 12 | Frequency | AGAP002886-\|baseline | AGAP002886-\|OA1 | 2.44E-114 | 3.70E-113 | < 2e-16 | **** | Wilcoxon |
| 13 | Frequency | G3\|OA1 | AGAP000045-\|OA1 | 0.000768 | 0.0015 | 0.00077 | *** | Wilcoxon |
| 14 | Frequency | G3\|OA1 | AGAP002886-\|OA1 | 7.01E-54 | 8.40E-53 | < 2e-16 | **** | Wilcoxon |
| 15 | Frequency | AGAP000045-\|OA1 | AGAP002886-\|OA1 | 1.88E-85 | 2.60E-84 | < 2e-16 | **** | Wilcoxon |
| 16 | Amplitude | G3\|baseline | AGAP000045-\|baseline | 1.94E-53 | 2.70E-52 | < 2e-16 | **** | Wilcoxon |
| 17 | Amplitude | G3\|baseline | AGAP002886-\|baseline | 2.58E-23 | 2.80E-22 | < 2e-16 | **** | Wilcoxon |
| 18 | Amplitude | G3\|baseline | G3\|OA1 | 0.930438 | 1 | 0.93044 | ns | Wilcoxon |
| 19 | Amplitude | G3\|baseline | AGAP000045-\|OA1 | 4.63E-19 | 4.60E-18 | < 2e-16 | **** | Wilcoxon |
| 20 | Amplitude | G3\|baseline | AGAP002886-\|OA1 | 6.85E-55 | 1.00E-53 | < 2e-16 | **** | Wilcoxon |
| 21 | Amplitude | AGAP000045-\|baseline | AGAP002886-\|baseline | 3.31E-10 | 2.30E-09 | 3.30E-10 | **** | Wilcoxon |
| 22 | Amplitude | AGAP000045-\|baseline | G3\|OA1 | 7.70E-36 | 1.00E-34 | < 2e-16 | **** | Wilcoxon |
| 23 | Amplitude | AGAP000045-\|baseline | AGAP000045-\|OA1 | 0.000627 | 0.0025 | 0.00063 | *** | Wilcoxon |
| 24 | Amplitude | AGAP000045-\|baseline | AGAP002886-\|OA1 | 0.883121 | 1 | 0.88312 | ns | Wilcoxon |
| 25 | Amplitude | AGAP002886-\|baseline | G3\|OA1 | 1.51E-12 | 1.40E-11 | 1.50E-12 | **** | Wilcoxon |
| 26 | Amplitude | AGAP002886-\|baseline | AGAP000045-\|OA1 | 0.285161 | 0.86 | 0.28516 | ns | Wilcoxon |
| 27 | Amplitude | AGAP002886-\|baseline | AGAP002886-\|OA1 | 3.41E-09 | 2.00E-08 | 3.40E-09 | **** | Wilcoxon |
| 28 | Amplitude | G3\|OA1 | AGAP000045-\|OA1 | 1.44E-10 | 1.20E-09 | 1.40E-10 | **** | Wilcoxon |
| 29 | Amplitude | G3\|OA1 | AGAP002886-\|OA1 | 3.04E-30 | 3.70E-29 | < 2e-16 | **** | Wilcoxon |
| 30 | Amplitude | AGAP000045-\|OA1 | AGAP002886-\|OA1 | 0.000448 | 0.0022 | 0.00045 | *** | Wilcoxon |
